# Transcriptional Network Orchestrating Regional Patterning of Cortical Progenitors

**DOI:** 10.1101/2020.11.03.366914

**Authors:** Athéna R Ypsilanti, Kartik Pattabiraman, Rinaldo Catta-Preta, Olga Golonzhka, Susan Lindtner, Ke Tang, Ian Jones, Armen Abnousi, Ivan Juric, Ming Hu, Yin Shen, Diane E Dickel, Axel Visel, Len A Pennachio, Michael Hawrylycz, Carol Thompson, Hongkui Zeng, Iros Barozzi, Alex S Nord, John Rubenstein

## Abstract

We uncovered a transcription factor (TF) network that regulates cortical regional patterning. Screening the expression of hundreds of TFs in the developing mouse cortex identified 38 TFs that are expressed in gradients in the ventricular zone (VZ). We tested whether their cortical expression was altered in mutant mice with known patterning defects (*Emx2, Nr2f1* and *Pax6)*, which enabled us to define a cortical regionalization TF network (CRTFN). To identify genomic programming underlying this network, we performed TF ChIP-seq and chromatin-looping conformation to identify enhancer-gene interactions. To map enhancers involved in regional patterning of cortical progenitors, we performed assays for epigenomic marks and DNA accessibility in VZ cells purified from wild-type and patterning mutant mice. This integrated approach has identified a CRTFN and VZ enhancers involved in cortical regional patterning.

## INTRODUCTION

Cortical development is an intricate choreography that integrates finely tuned molecular states such as gradients of transcription factors (TFs) and the chromatin landscape with a variety of cellular processes (cell type specification, proliferation, differentiation and migration). Together, these developmental processes result in the establishment of an adult cerebral cortex, and associated pallial structures, which are organized into discrete areas and laminae distinguished by specific molecular signatures (Ortiz et al, 2020). The earliest steps in the generation of cortical areas (regionalization process) is regulated in mouse by FGF-signaling and by TFs such as ARX, DMRT3, DMRT5, EMX2, LEF1, LHX2, NR2F1, NR2F2, PAX6, PBX1 and SP8; TFs which exhibit graded expression patterns along the rostral/caudal and dorsal/ventral axes in the neuroepithelium of the developing pallium (O’Leary et al., 2007; Arai and Pierani, 2014; Desmaris et al., 2019; Saulnier et al., 2013; Golonzhka et al., 2015). These TFs are crucial for early patterning of the cortex and their loss causes shifts in cortical regional structures (Bishop et al., 2000; Mallamaci et al., 2000; Bishop et al. 2003; Armentano et al, 2007). As the cortex develops, regional identity is then translated from the VZ to secondary progenitors of the subventricular zone (SVZ) and then to neurons of the cortical plate (CP) (Greig et al., 2013).

To understand cortical patterning, one needs to comprehend the regulatory architecture of individual genes that play a role in regionalization. Work on the regulatory control of individual TF genes identified enhancer elements that control gene expression of specific patterning TFs such as *Emx2* (Theil et al, 2002; Suda et al, 2010) and *Pax6* (Kammandel et al., 1999; McBride et al. 2011; Mi et al. 2013). A larger scale effort identified E14.5 pallial enhancers by ChIP-Seq for p300 (an enhancer-associated co-activator) and showed activity for 8 enhancers (Wenger et al, 2010). Similarly, we performed a large-scale identification of forebrain putative regulatory elements (pREs) and found 145 pREs active in the forebrain at E11.5 (Visel et al, 2013). This work showed that many cortical pREs had sharp boundaries of activity in the neuroepithelium. In a follow-up study, we discovered that TF binding at individual pREs enabled them to integrate information from TF gradients in the developing pallium to generate discrete regional domains prenatally and postnatally (Pattabiraman et al, 2014). Individual enhancers with discrete neuroepithelial activity can together encode regional expression gradients, and thus individual pRE activity may reflect the highest resolution of transcriptional differences within the developing cortex and serve as a protomap of cortical areas (Rakic, 1988).

Herein, we built on our previous work with the aim to elucidate components of the transcriptional network that orchestrates regionalization of the cortical neuroepithelium. We focused on 38 cortical progenitor TFs that have graded RNA distribution along the rostrocaudal (RC) and/or dorsoventral (DV) axes. We refer to these 38 TFs as the Cortical Regionalization TF Network (CRTFN) (Figure 1E). Towards defining the transcriptional logic of cortical patterning, we assessed changes in CRTFN expression in the context of known cortical patterning defects by using *Emx2, Nr2f1* and *Pax6* loss-of-function mouse mutants. We found that rostroventral patterning involves *Pax6* promoting the expression of *Nr2f1* and *Nr2f2*, which in turn promotes *Lmo3* and *Npas3*. We show that loss of *Npas3* function also alters rostroventral cortical patterning.

**Figure 1.**
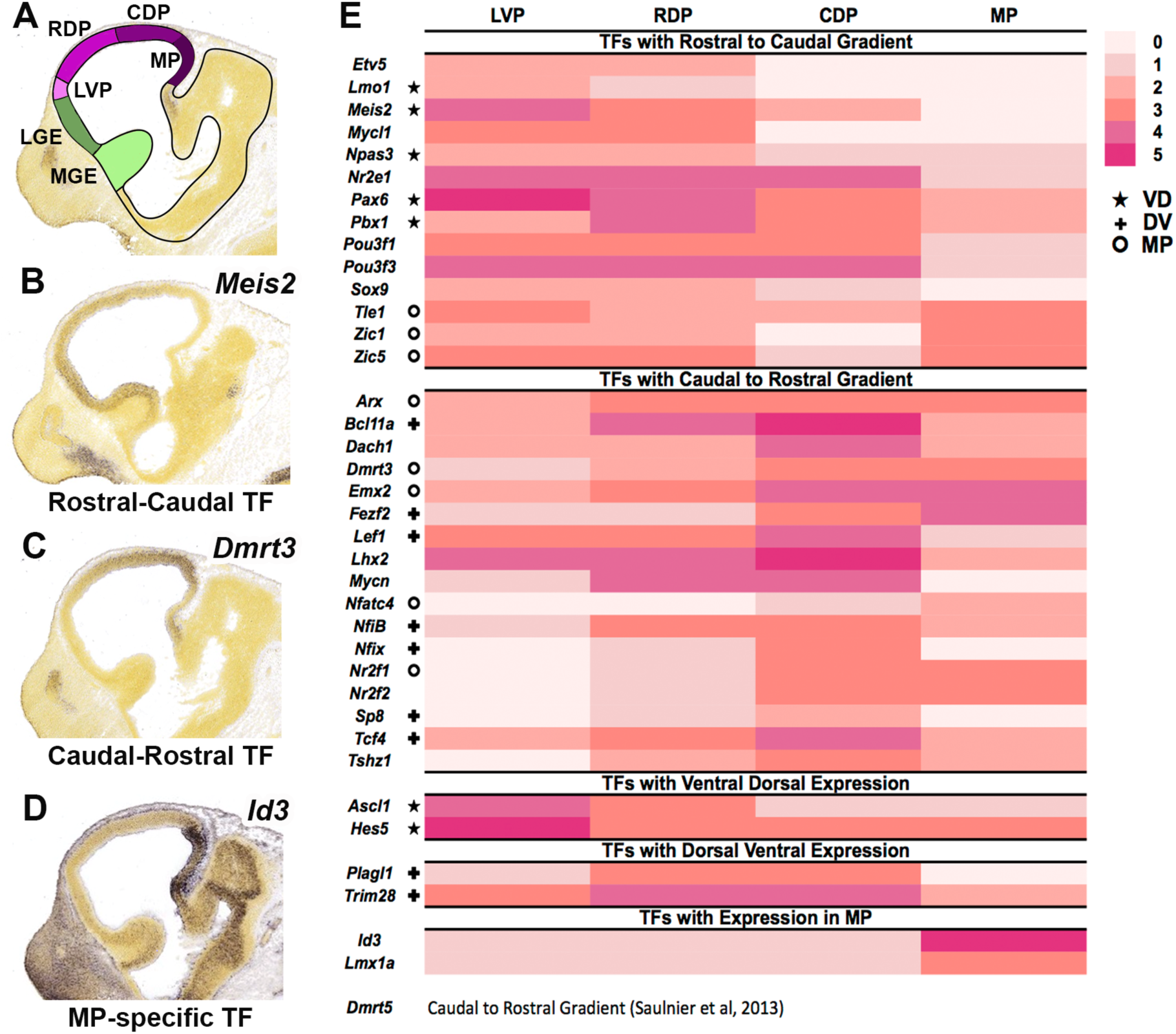
Annotation of TF expression in the Cortical Regionalization TF Network (CRTFN) in the E11.5 cortex. (A) Schema of sagittal view of E11.5 mouse brain. The pallium is in four shades of purple corresponding to regional subdivisions (LVP: laterovental pallium, RVP: rostrovental pallium, CDP: caudodoral pallium, MP: medial pallium). (B-D) ISH analysis of *Meis2* (B), *Dmrt3* (C) and *Id3* (D). (E) Heatmap of cortical expression levels (0-5: pink to red scale) of CRTFN TFs in the four pallial subdivisions. **★** indicates that the TF also has ventrodorsal gradient. **✚** indicates that the TF also has dorsoventral gradient. ° indicates that the TF also has MP expression. Abbreviations: LGE: lateral ganglionic eminence, MGE: medial ganglionic eminence, MP: medial pallium.

To identify the genomic REs tying together the CRTFN we performed TF chromatin immunoprecipitation and DNA sequencing (ChIP-seq) for EMX2, LHX2, NR2F1, PAX6, and PBX1. This analysis, coupled to histone ChIP-seq and ATAC-seq, identified ∼2000 pREs in the CRTFN. Combinatorial TFs binding was tightly associated with the activity of cortical enhancers. Furthermore, coordinated binding of EMX2, LHX2, NR2F1, PAX6, and PBX1 provided evidence for a combinatorial signature regulating enhancer activity along the rostroventral-caudodorsal axes. Finally, by assessing epigenetic changes of the 2000 CRTFN pREs in *Emx2, Nr2f1* and *Pax6* mutants, we identified enhancers likely to participate in cortical patterning. Together, this study establishes the foundations of the transcriptional circuitry underlying cortical regionalization.

## RESULTS

### Pallial Expression of 698 TFs at E11.5

To systematically identify TFs that could participate in cortical regional patterning through their activity in neural progenitors, we used the Allen Brain Developing Mouse Atlas to study the RNA expression of 698 TFs based on *in situ* hybridization (ISH) analyses on sagittal sections of E11.5 mouse embryos.

We found that 270 TFs showed pallial expression, whereas pallial expression of 428 TFs was not detected. We described the pallial expression of all TFs based on their regional and laminar expression patterns, using topological schemata that define pallial expression in lateroventral, rostrodorsal, caudodorsal and medial pallial regions (LVP, RDP, CDP, MP, respectively) (Figure 1A, Table S1). Herein, we focus on progenitor zone expression, as we wished to concentrate on mechanisms that operate in progenitors and control pallial regional patterning. We assessed the level of expression based on both the intensity and the density of expression (0-5 scale). Figure 1E shows TFs that were expressed in gradients in progenitors at E11.5 in the LVP, RDP, CDP and MP. In this analysis, we identified 218 TFs whose telencephalic expression was specific to the ventricular zone of the pallium and/or showed a regional or graded pattern within the pallium, with an expression intensity of 2 or more. There were just a few TFs with regional expression boundaries: 10 largely restricted to the DP, 6 largely restricted to the MP (i.e. *Id3*, Figure 1D) and 1 largely restricted to the LVP. More commonly, TFs were broadly expressed across the pallium (n=134), with many showing either rostrocaudal (RC, n= 30) or caudorostral (CR, n= 31) gradients. For instance, *Dmrt3* showed a CR gradient and *Meis2* showed a RC gradient at E11.5 (Figure 1B, C).

We hypothesize that our screen and subsequent analysis identified most of the TFs that are candidate VZ regulators of pallial patterning at E11.5. We will refer to these TFs as the Cortical Regionalization TF Network (CRTFN). This screen also provides evidence that there are few TFs with restricted intrapallial expression domains except in the MP, supporting the idea that most pallial subdivisions are not generated by highly restricted TF expression within pallial progenitors.

### *Emx2, Nr2f1 and Pax6* Regulate Gradients of CRTFN Expression at E11.5

Next, we investigated how CRTFN TFs are regulated by TFs with known functions in cortical patterning. We studied the expression of 31 TFs in *Emx2, Nr2f1* and *Pax6* mouse mutants at E11.5 (Figures 2, S6). We chose to study the 31 TFs based on the following three criteria: 1) TFs previously not well-known to be expressed in the E11.5 pallium; 2) TFs whose neurodevelopmental functions were poorly known; and 3) TFs with clear E11.5 expression gradients.

**Figure 2.**
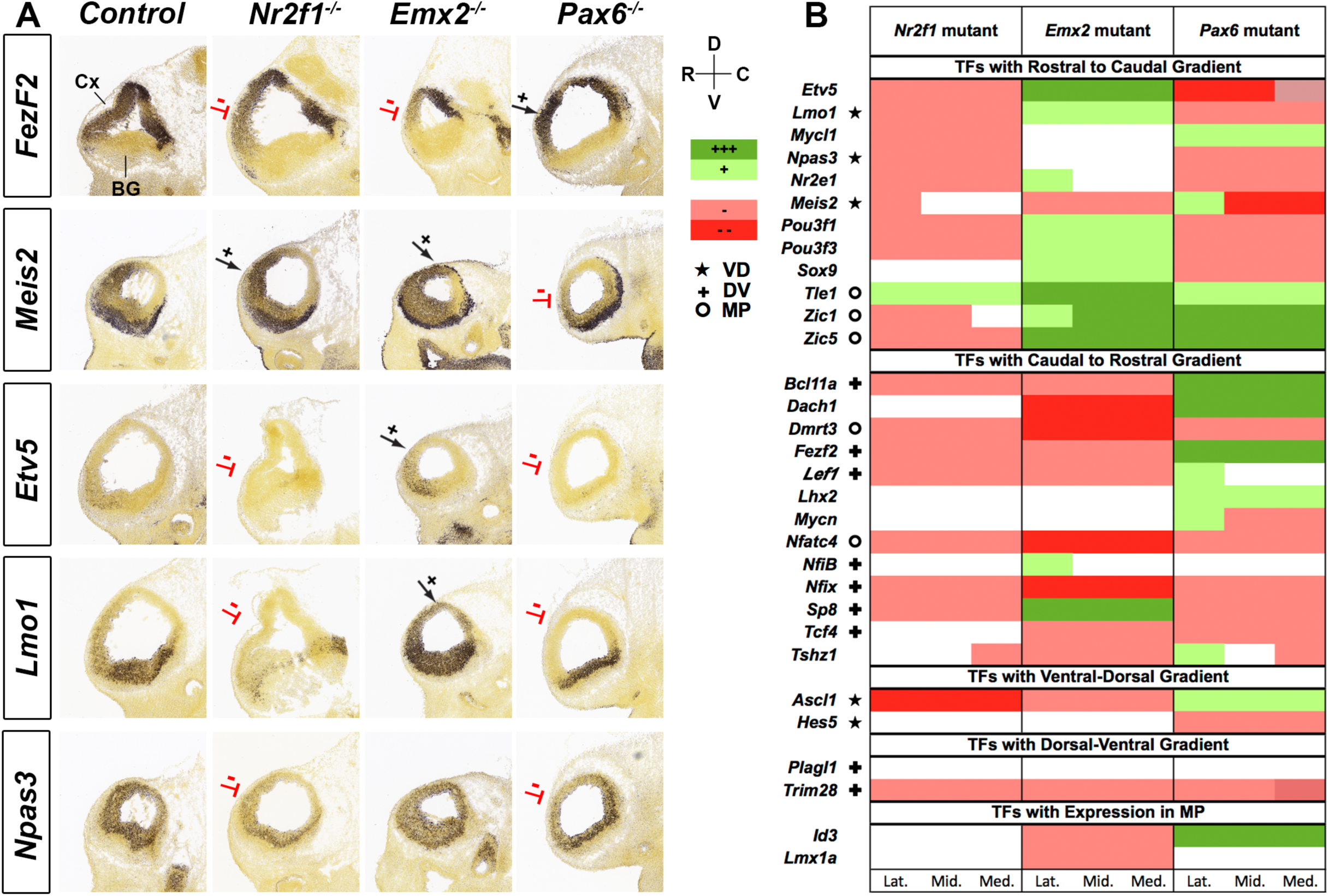
Graded expression changes of CRTFN TFs in *Nr2f1, Emx2* and *Pax6* mutants. *(A)*ISH shows changes in one caudorostral TF (*Fezf2)* and four rostrocaudal TFs (*Meis2, Etv5, Lmo1* and *Npas3)* in E11.5 sagittal sections of *Nr2f1, Emx2* and *Pax6* mutants. Red lines denote reduced expression, whereas black arrows indicate increased expression. The RC and DV axes are depicted on the top-right of the sections. *(B)*Heatmap showing changes in expression of 31 TFs in *Nr2f1*^*-/-*^, *Emx2*^*-/-*^ and *Pax6*^*-/-*^ using a 5-level qualitative scale: increased expression (dark green +++ or light green +); no change (white); decreased expression (light red - or dark red --). TFs are categorized according to their gradient in WT (RC, CR, VD or DV) or their expression in a restricted region (MP). Changes are annotated in Lateral (Lat.), Middle (Mid.) and Medial (Med.) sagittal sections. **★** indicates that the TF also has ventrodorsal gradient. **✚** indicates that the TF also has dorsoventral gradient. ° indicates that the TF also has MP expression. Abbreviations: Cx: cortex. BG: basal ganglia.

The phenotypic descriptions of the effects of the *Emx2, Nr2f1* and *Pax6* mutations on the expression of the 31 TFs are organized based on changes of their RC, CR or regional expression. We annotated the expression changes based on the independent assessment of 3 experts using a 5-level qualitative expression scale: increased expression (green; ++ or +); no change; decreased expression (red; -- or -)(Fig. 2B). The expression changes were assessed at lateral, middle and medial levels of sagittal sections at E11.5 and the expression change at each level was denoted in Figure 2. Representative sections for each TF in the three mutants can be found in Figure S5. All but one TF (*Plagl1*) showed differential RNA expression in at least one of the 3 mutants.

During cortical patterning, EMX2 and NR2F1 promote caudal identity, whereas PAX6 promotes rostral identity (O’Leary et al., 2007; Cadwell et al., 2019). Considering the regional patterning role of these TFs, we hypothesized that genetic ablation would lead to loss of regional transcriptional activation along the rostral-caudal axis. Consistent with this, genes with CR gradients (i.e. increased caudal expression relative to rostral) were more likely to show decreased expression in *Emx2*^*-/-*^ (9/13) and in *Nr2f1*^*-/-*^ (8/13) than in *Pax6*^*-/-*^ (5/13). Likewise, genes with RC gradients showed reduced expression in *Pax6*^*-/-*^ (8/12) and increased expression in *Emx2*^*-/-*^ (9/12) (Figure 2, S6). However, unlike in *Emx2*^*-/-*^, loss of *Nr2f1* led to a decrease in expression of RC TFs (9/12).

Although *Nr2f1* and *Emx2* are expressed in CR gradients, they have opposite dorsoventral (DV) gradients. *Emx2* has a DV gradient, whereas *Nr2f1*, like *Pax6*, has a ventrodorsal (VD) gradient (Figure S1A). Thus, perhaps the TF expression differences between the *Emx2* and *Nr2f1* mutants might in part relate to their differences in DV patterning functions. Indeed, in the *Nr2f1* knockout, several down-regulated RC TFs (including *Npas3*) were also down-regulated in the *Pax6* mutant (Figure 2B). This led us to explore whether *Pax6 and Nr2f1* have similar roles in regulating the ventral patterning of the cortex.

### *Nr2f1/2* and *Pax6* regulate patterning of the rostral ventrolateral pallium

The loss of function experiments described above suggest that TFs with rostral-ventral gradients (such as *Npas3*) are similarly regulated by *Pax6* and *Nr2f1*. Loss-of-function analyses show that *Pax6* has prominent roles in promoting LVP properties (Yun et al., 2001). In *Pax6* mutants we found that LVP expression of both *Nr2f1* and *Nr2f2* were strongly reduced at E12.5 (Figure 3A,B), suggesting that patterning of the ventral pallium by *Pax6* involves *Nr2f1* and *Nr2f2*. Previous work did not find LVP defects in *Nr2f1*^*-/-*^ (Faedo et al., 2008). However, because *Nr2f1* and *Nr2f2* expression are very similar, they may compensate for each other. Therefore, we analyzed *Nr2f1/2* conditional mutants (cKOs; *Nr2f1*^*-/-*^; *Nr2f2*^*f/f*^; *Emx1-cre*). We first assessed whether they co-regulated *Pax6* expression in the LVP at E11.5, but detected no change in PAX6 immunostaining (Figure 3C). We then examined rostroventral cortex phenotypes at later ages in both *Pax6*^*-/-*^ and *Nr2f1/2* cKOs (Figure 3H). In the rostroventral cortex at E16.5, ventral cortex markers, *Nurr1* and *Npas3*, were strongly reduced in both the *Pax6*^*-/-*^ and *Nr2f1/2* cKOs. (Figure 3E,F). *Nurr1* expression was maintained ventrocaudally in the *Pax6* mutant, but was lost in the *Nr2f1/2* cKOs (Figure 3F). Among other ventral cortex markers, *Lmo3* exhibited reduction in expression in *Pax6*^*-/-*^ and not the *Nr2f1/2* cKOs (Figure 3D). This suggests that *Pax6* can also work through a *Nr2f1/2*-independent pathway to specify the ventral cortex.

**Figure 3.**
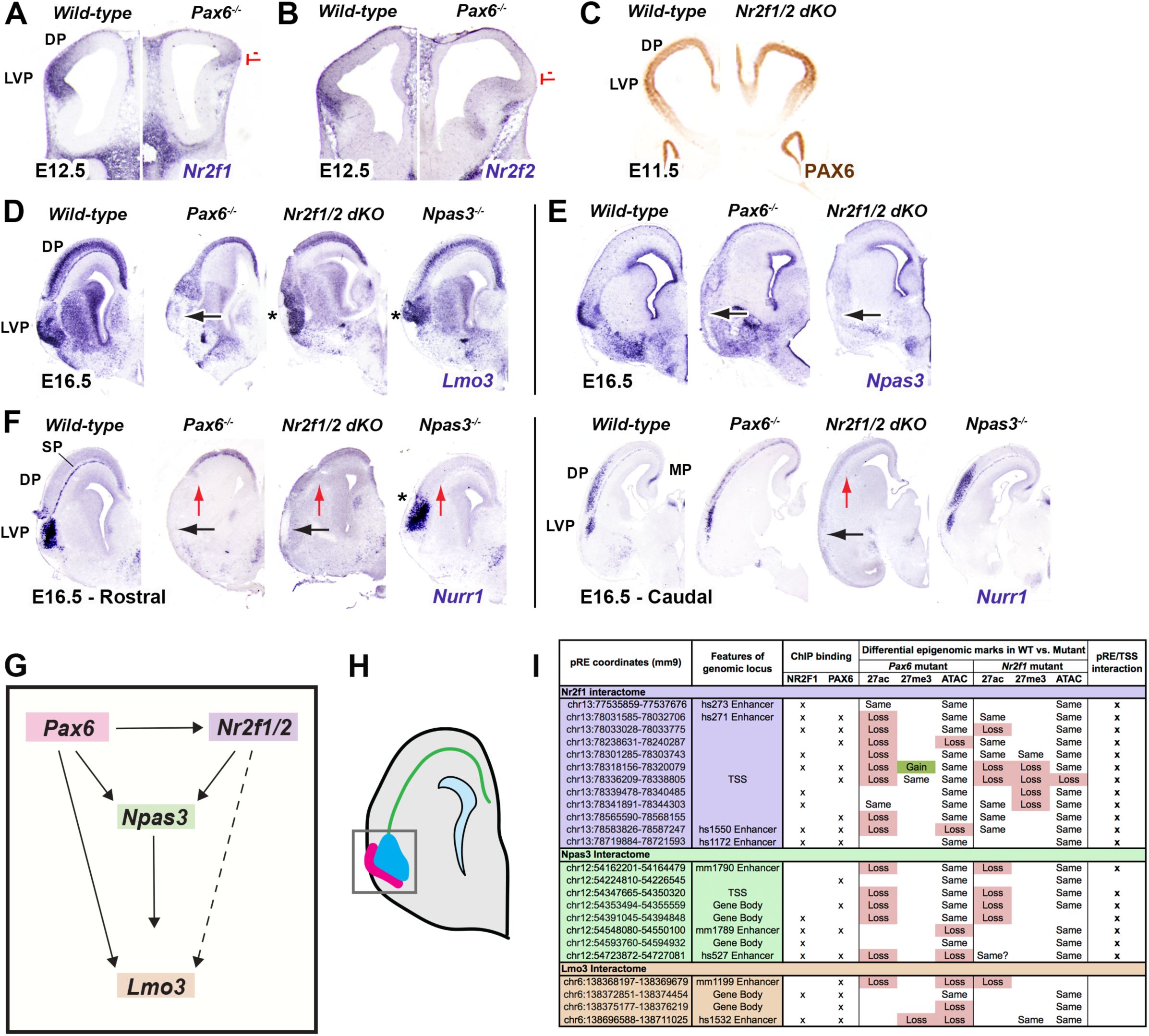
*Pax6*-*Nr2f1/2*-*Npas3* pathway promotes patterning of the rostral lateroventral pallium (R LVP) and subplate. (A,B) At E12.5, *Nr2f1 and Nr2f2* ISH expression is decreased in the LVP and DP of coronal sections in *Pax6*^*-/-*^ (red lines). (C)At E11.5, PAX6 immunohistochemistry shows no change in the *Nr2f1/2* dKO. (D-F) E16.5, ISH shows changes of R LVP markers (*Lmo3, Npas3* and *Nurr1)* in *Pax6*^*-/-*^, *Nr2f1/2* dKO and *Npas3*^*-/-*^ in rostral sections. Caudal sections are also depicted for *Nurr1* ISH. Black arrow: Loss of expression in the R LVP; Star (*): abnormal morphology of the R LVP; Red arrow: loss of the subplate (SP). (G) Schematic of transcriptional control of R LVP development. (H) Depiction of the R LVP at E16.5. Piriform cortex (*Lmo3*^*+*^ and *Npas3*^*+*^) is pink. Endopiriform cortex and claustrum (*Nurr1*^*+*^) are blue. Subplate is green. (I)Table of regulatory elements (pREs) around the *Nr2f1, Npas3* and *Lmo3* loci, that may participate in R LVP patterning Shown are NR2F1 and PAX6 ChIP-seq peaks, differential epigenomic peaks in the *Nr2f1*^*-/-*^ *and Pax6*^*-/-*^ and pRE/TSS interactions (see Figures 4, 6, 7).

We next examined whether some of the rostroventral TFs that were coregulated by *Pax6* and *Nr2f1* could in turn regulate rostroventral cortical development. To that end, we examined the rostroventral cortical phenotypes in the *Npas3* null mice (Erbel-Sieler et al., 2004). We found that the rostroventral cortex in *Npas3*^-/-^ appeared dysmorphic based on *Lmo3* and *Nurr1* expression (Figure 3D,F). Moreover, as in *Pax6*^-/-^, *Npas3*^-/-^ maintained ventrocaudal *Nurr1* expression. The core components likely to participate in the transcriptional regulation of patterning the rostroventral cortex (Figure 3H) are schematized in Figure 3G (characterization of *Nr2f1, Npas3* and *Lmo3* genomic loci in Figure 3I will be explained hereafter).

### Analysis of EMX2, LHX2, NR2F1, PAX6 and PBX1 binding to genomic regions associated with the Cortical Regionalization TF Network (CRTFN)

Next, we investigated whether PAX6, EMX2 and NR2F1 proteins bind to regulatory regions of the CRTFN genes (Table S2). Our goal was to determine whether they directly regulate these genes and to identify pREs for the CRTFN. We performed chromatin immunoprecipitation-DNA sequencing experiments (ChIP-seq) using PAX6, EMX2 and NR2F1 specific antibodies on E12.5 cortical tissue (Pattabiraman et al., 2014 and here). We also chose to include ChIP-seq data for two other patterning cortical TFs that are expressed in progenitor cells, namely LHX2 and PBX1 (Golonzhka et al., 2015), because the mutants for these TFs exhibit patterning defects (Chou et al., 2009; Shetty et al., 2013; Golonzhka et al., 2015). These five TFs are expressed in gradients in the pallial VZ: RC for *Pax6* and *Pbx1* and CR for *Emx2, Lhx2* and *Nr2f1* (Figure S1A). A description of the ChIP-seq for PAX6, PBX1 and NR2F1 was previously published (Pattabiraman et al., 2014; Golonzhka et al., 2015). The LHX2 antibody was previously used for ChIP-seq (Monahan et al., 2017). The EMX2 antibody was generated for this study; its specificity was tested by immunostaining in wild-type mice as well as by performing ChIP-seq with an EMX2 blocking peptide (EMX2 Rep2, Figure 4A). Representative ChIP-seq peaks are depicted around the *Pax6* locus in Figure 4A. TF ChIP peaks are shown around the *Pax6* locus and called peaks are annotated beneath the genomic tracks. Overlapping TF ChIP-seq peaks were merged into discrete pREs (highlighted in pink). We assessed the reproducibility of the ChIP-seq experiments by pairwise Pearson genome-wide read count correlation of biological replicates run individually, as depicted in Figure S1C. For each TF, ChIP binding sites were located predominantly at distal rather than proximal genomic loci (Figure S1B). We defined the size of each genomic locus by defining its boundaries by: 1) the distal point of contact for promoter-enhancer looping obtained by a chromatin conformation assay (see below) 2) ± 100kb from the limits of the gene body.

**Figure 4.**
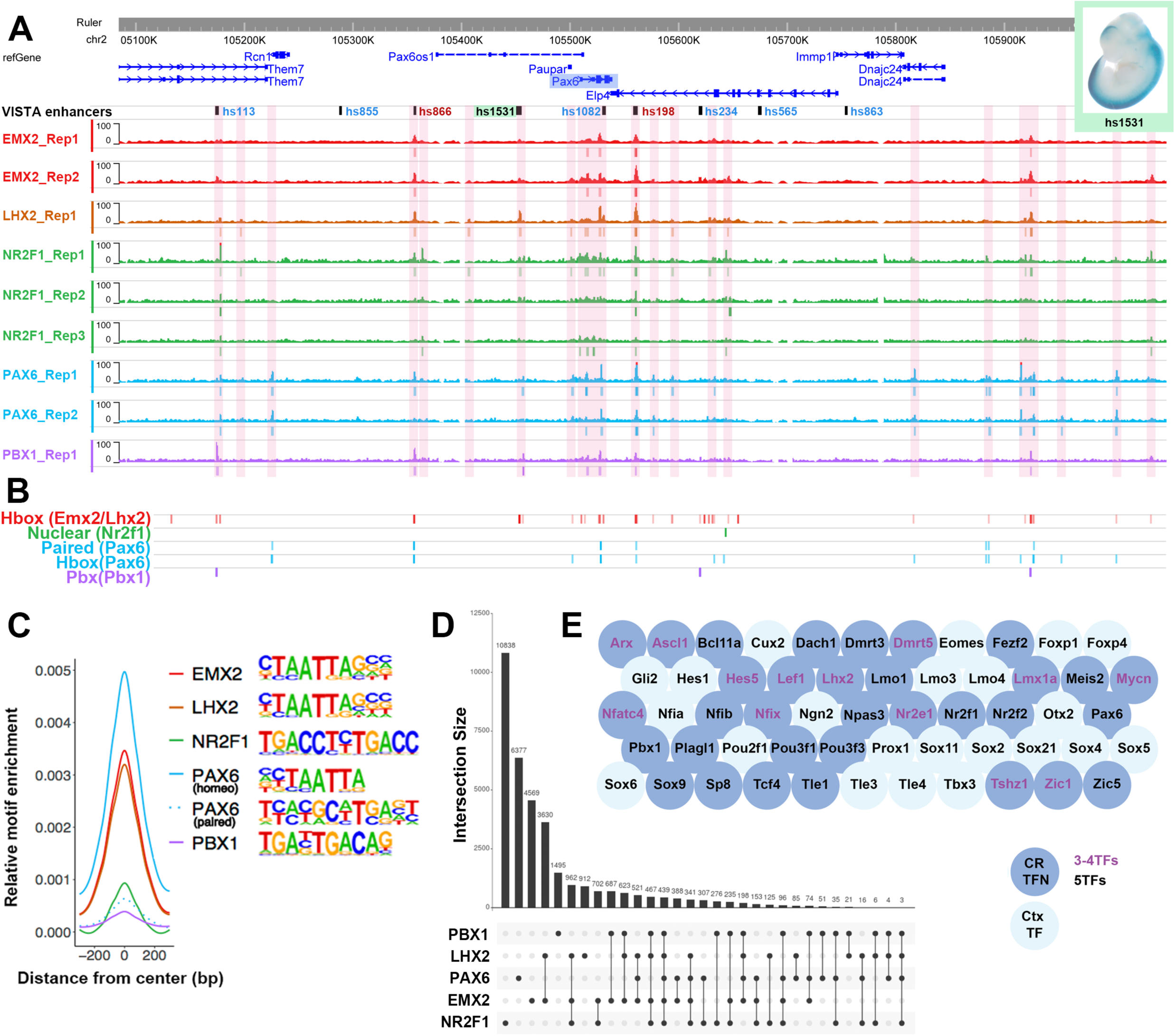
Combinatorial binding of EMX2, LHX2, NR2F1, PAX6, and PBX1 predicts genes in the CRTFN. (A) EMX2, LHX2, NR2F1, PAX6 and PBX1 ChIP-seq coverage at the *Pax6* locus at E12.5. pREs are highlighted in pink. VISTA enhancers are also indicated in green (cortical activity; i.e. hs1531, wholemount with positive blue LacZ staining, top-right), in blue (non-cortical activity; i.e. hs113) and in red (no activity at E11.5; i.e. hs198). (B) Genomic location of the EMX2, LHX2, NR2F1, PAX6 and PBX1 primary binding motifs in the *Pax6* locus. (C) Centered distribution of EMX2, LHX2, NR2F1, PAX6 and PBX1 motifs across ChIP-seq peaks. Motifs that were strongly enriched within ChIP peaks were homeobox motifs in EMX2, LHX2, PAX6 and PBX1, the paired motif in PAX6 and the nuclear receptor motif in NR2F1. Primary DNA binding sequences are indicated to the right. (D) Number of peaks for each TF alone or in combination. Y axis shows number of peaks. X axis indicates single and combinatorial binding at pREs. (E) pREs with combinatorial binding of 3 or 4 (Purple letters) or 5 TFs (Black letters) are enriched around genomic loci for TFs from the CRTFN (blue circles, 30/35 TFs) as well as other TFs important in cortical development (light blue circles).

We used Homer (Heinz et al., 2010) to perform *de novo* motif discovery and enrichment analysis. We identified enriched sequence motifs including the primary binding motifs for EMX2, LHX2, PAX6 and PBX1 (homeoboxes), NR2F1 (nuclear receptor), and PAX6 (paired domain) (Figure 4C, S2E) (Epstein et al., 1994; Czerny and Busslinger, 1995; Iler et al., 1995; Montemayor et al., 2010; Golonzhka et al., 2015). These putative primary motifs were enriched in the center of pREs (Figure S1E, S1I) supporting direct TF binding. Beyond the primary motifs associated with TFs profiled here, many additional motifs were enriched within pREs, including motifs for TFs that are crucial for cortical development (i.e. SOX motifs), as well as for basal ganglia development (i.e. NKX and DLX motifs) (Figure S1F). Notably, when NR2F1 was bound at loci in conjunction with other TFs, the nuclear NR2F1 motif was rare, indicating that NR2F1 is likely to bind indirectly to such pREs (Figure S1I).

Many pREs exhibited combinatorial TF binding. Our hypothesis was that regions bound by multiple TFs are important for regulating pallial regional patterning in the developing cortical VZ. Thus, we looked for loci which were bound by all five TFs. At the whole genome level, these pREs were enriched at loci associated with transcriptional activity/DNA binding (GO molecular and cellular function) and genes with functions in forebrain and neuronal development (GO Biological Process and Mouse Phenotype, Figure S1D). These pREs were predominantly found at distal loci (Figure S1G). They were highly enriched around TF genes that are expressed in the embryonic pallium (blue bubbles in Figure 4E) as well as non-TFs that are implicated in cortical development (e.g. *BMPs, Ephrins, Semaphorins*, Figure S1H). Of particular note, pREs with combinatorial binding were enriched around TFs that we identified as being a part of the CRTFN (dark blue bubbles in Figure 4E). Considering the full set of pREs, we found 1236 unique TF binding sites associated with genes in the network (see Table S2), and 33/38 CRTFN genomic loci had at least one pRE co-bound by 3-5 TFs. In the CRTFN, we found 439 genomic regions where there was co-binding of 5TFs (Figure 4D). We hypothesize that these pREs are likely to participate in regulating TF gene expression during patterning of the cortical VZ.

### pREs bound by multiple TFs are more likely to be active enhancers

To assess the activity of pREs defined by ChIP binding, we explored whether they are located in known enhancers that are active at E11.5 (VISTA enhancers – https://enhancer.lbl.gov; Visel et al., 2013). These enhancers were primarily ascertained by screens of evolutionarily conserved sequences or p300 ChIP-seq, and thus represent an unbiased set with regard to the binding patterns of the TFs profiled here. We classified VISTA enhancers that are located around CRTFN genes based on where they are active at E11.5: Pallial (n=32), Subpallial (n=25), and Non-telencephalic (n=70). We also identified Inactive enhancers (n=121, Table S3). We found that TF binding (1-5 TFs) was present on 94% of Pallial, 64% of Subpallial, 56% of Non-telencephalic and, 33% of Inactive enhancers (Figure S4C). This indicates that nearly all genomic loci from this unbiased set of validated pallial enhancers exhibit binding of at least one of the CRTFN TFs.

Strikingly, 56% of the VISTA enhancers with Pallial activity were bound by four or five TFs. In comparison, only 20% of Subpallial, 8% of Non-telencephalic and 8% of Inactive VISTA enhancers showed this pattern (Figure 5C; Table S3). Thus, binding by 4-5 TFs appears to be a potent indicator of enhancer activity. Furthermore, we estimate that there are 115 such loci associated with the CRTFN. Thus, CRTFN pREs bound by multiple TFs are likely to have pallial activity (Figure S4C).

**Figure 5.**
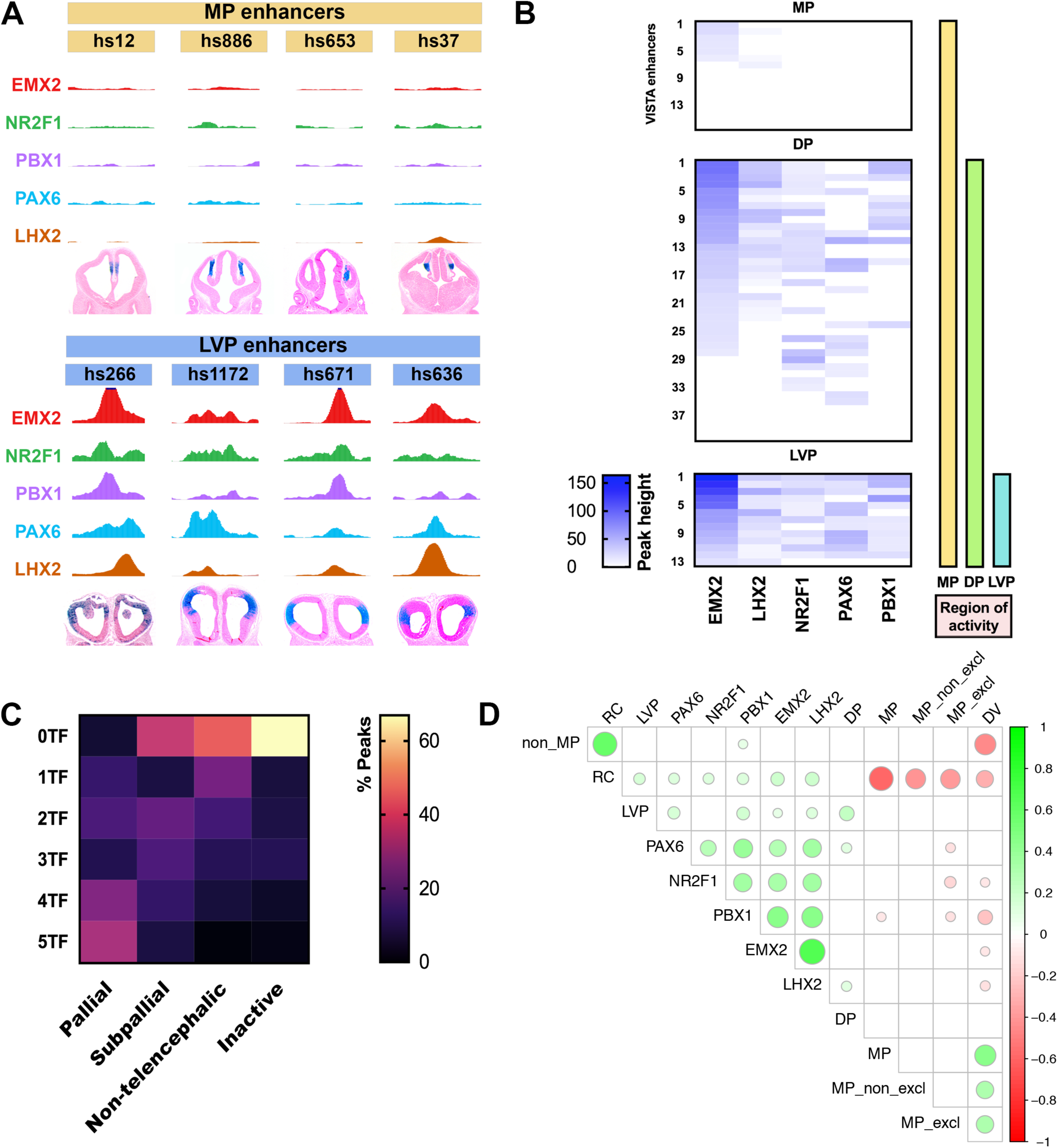
Combinatorial binding of EMX2, LHX2, NR2F1, PAX6, and PBX1 predicts gradients of enhancer activity in the pallium. (A)LVP enhancers (hs266, hs1172, hs671 and hs636) are enriched for ChIP-seq peaks but MP enhancers (hs12, hs886, hs653 and hs37) lack peaks. (B)Heatmap showing enrichment (peak height) of EMX2, LHX2, NR2F1, PAX6 and PBX1 binding over VISTA enhancers with MP exclusive activity (n=16), MP + DP activity (n=40) and MP + DP + LVP activity (n=13) symbolized by the yellow, green and blue vertical bars. Each row represents a distinct VISTA enhancer and the coverage of TF peaks at that locus. (C) Heatmap of combinatorial TF binding (percentage enrichment; 0-5 TFs) on VISTA enhancers that have pallial, subpallial, non-telencephalic, and no activity (inactive). (D) Correlalogram of computational modeling showing the different combinations of TF binding and their predictions of gradients of activity (p < 0.01). Green circles indicate correlation and red circles indicate anti-correlation. The size of the circles is associated with the strength of the correlation/anti-correlation. For instance, RC enhancer activity is correlated with PAX6, NR2F1, PBX1, EMX2 and LHX2 TF ChIP-Seq.

We hypothesized that TF binding would predict regional pRE activity. To test this, we leveraged the information provided by TF combinatorial binding and applied it in the context of cortical gradients and/or cortical subregions.

### Expression gradients are regulated by the combinatorial binding of EMX2, LHX2, NR2F1, PAX6, and PBX1

We were interested in determining whether combinatorial binding of TFs was predictive of the activity pattern of pREs and/or of their cognate genes. We looked at VISTA enhancers that specifically had cortical expression patterns and classified their expression in cortical subregions: LVP, DP and MP (as in Figure 1A, see Table S4). Next, we correlated the ChIP-seq peak heights, normalized for coverage depth differences across TFs, and found association between ChIP-seq signal strength and cortical regional activity (Figure 5B). While ChIP-seq signal was present across a large proportion of regions, LVP active enhancers appeared to exhibit a particularly strong ChIP-seq signal (Figure 5A,B).

Enhancers with LVP activity had the most robust peaks (>25% percentile for peak enrichment, 100%, n=13, Figure 5A,B). In contrast, peaks over enhancers whose activity was exclusively in the MP displayed no TF peaks in ∼56% of cases (n=12) or small peaks (<25% percentile for peak enrichment) in ∼44% of cases (n=6). Peaks that were over enhancers with DP activity (88% peaks, n=25), or in mixed regions (DP+MP – 100% peaks, n=14), displayed more heterogeneity in peak height but were often between both extremes observed in LVP and MP VISTA enhancers. TF binding and peak height enrichment for each individual VISTA enhancers, that have regional activity, is shown in Figure 5B. Thus quantitative enrichment from ChIP-seq suggests stronger TF-pRE interactions at LVP and DP pallial enhancers.

Some VISTA enhancers had spatial activity gradients along the pallial RC and DV axes. For instance, hs798 has both CR and DV gradients, whereas hs636 has RC and VD gradients (Figure S2B). To model our findings, we examined the correlation of TF binding, enhancer spatial activity gradients, and regionally-defined enhancer activity. We found that peak presence was correlated with RC and LVP enhancer activity (Figure 5D, S2C). In contrast, the presence of EMX2, LHX2, NR2F1 and PBX1, was anti-correlated with MP and DV enhancers (p < 0.01 for all correlations). This implies that the binding of these 5 TFs promotes activity of enhancers with RC and/or LVP activity, but not enhancers with MP activity. These results provide a gene regulatory logic where TF-regulatory element interactions generate spatial gradients across the developing pallium.

### Profiling the epigenomics of cortical progenitors

To link TF binding with pRE activity, we profiled chromatin state in wild-type cortical VZ cells isolated using the FlashTag method (Telley et al., 2016) (Figure 6A, S3A, S3B). From these VZ cells, we made nuclei and performed a chromatin accessibility assay (ATAC-seq), and histone ChIP-seq with antibodies to histone modifications associated with active pRE state (H3K27ac) and repressed pRE state (H3K27me3) (Lindtner et al., 2019; Markenscoff-Papadimitriou et al., 2020). In CRTFN loci, we found 749 ATAC, 556 H3K27ac and 564 H3K27me3 peaks. Examples of VZ epigenomic peaks are illustrated in the *Lef1* locus, with pREs highlighted in pink (Figure 6B).

**Figure 6.**
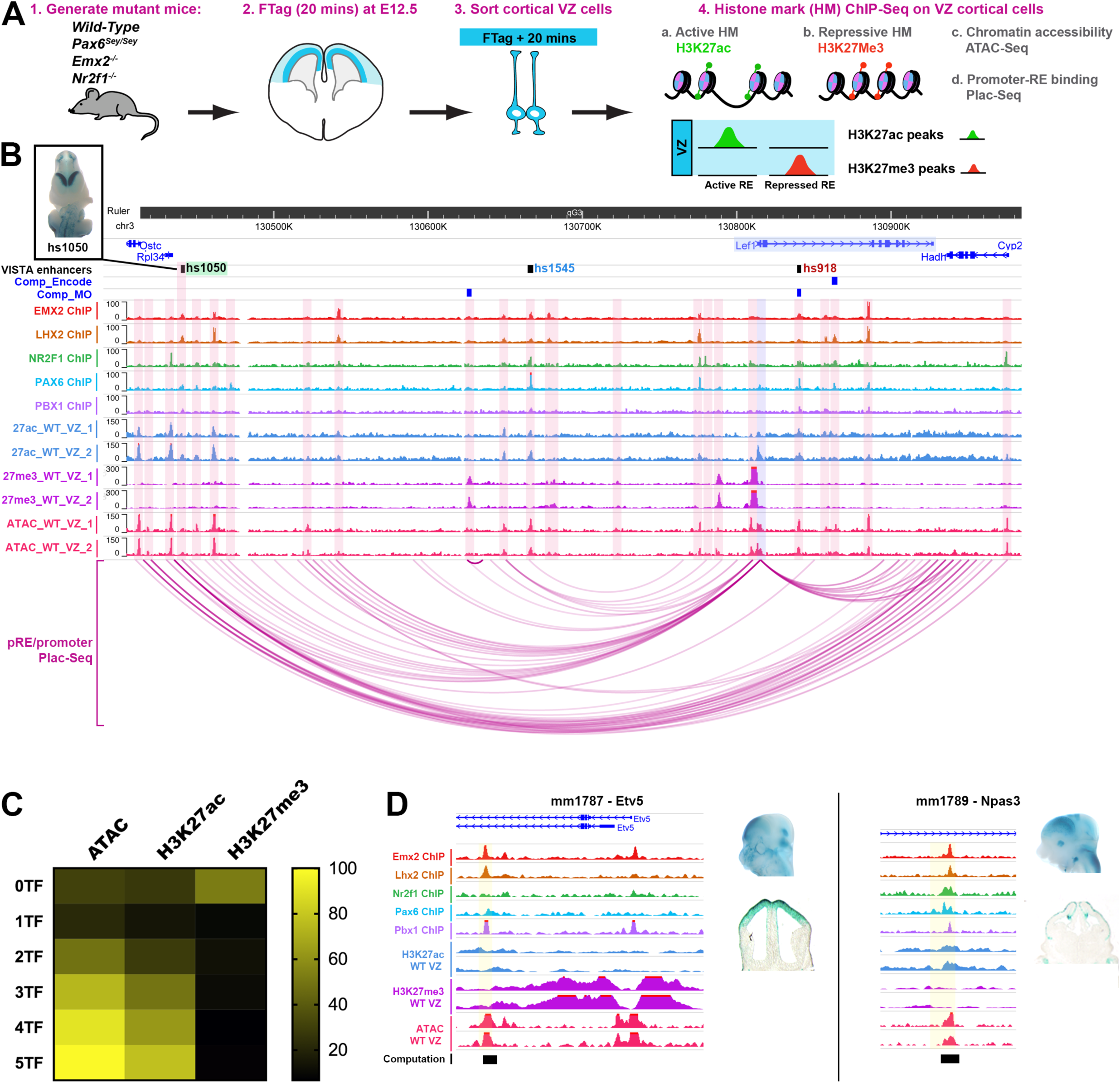
pRE landscape around TF genes in CRTFN in cortical progenitors. (A) Method for VZ cell purification for epigenomic experiments (see STAR methods). (B) Coverage of TF ChIPs, VZ histone marks (active: H3K27ac; repressive: H3K27me3) and VZ ATAC-seq in the *Lef1* locus at E12.5. pRE-promoter interactions are derived from PLAC-seq (purple arcs below) and computationally (blue boxes above, Comp_Encode and Comp_MO; Table S5).. VISTA enhancers are also indicated (green = cortical activity, blue = non-cortical activity, red = inactive).. (C) Heatmap of percentage enrichment of epigenomic marks over pREs with differential combinatorial TF binding (0-5 TFs). (D) Enrichment of TF and epigenomic marks over newly identified cortical enhancers near CRTFN genes: mm1787 (*Etv5*) and mm1789 (*Npas3*).

In general, there was a high degree of overlap of TF binding, H3K27ac and ATAC peaks; H3K27me3 peaks were less likely to be centered on TF, ATAC and H3K27ac peaks. The proportion of overlap with TF binding (1-5 TFs) was 40% for ATAC peaks, 26% for H3K27ac and 10% for H3K27me3 (Figure 6C). Interestingly, as TF binding increased from 0 to 5 TFs, the likelihood of a pRE having an ATAC or H3K27ac peak increased proportionally (Figure 6C). All pREs where 5 TFs were bound had ATAC peaks, and ∼78% had H3K27ac (active) marks. An inverse relationship was found between H3K27me3 marks and TF binding; only ∼10% of 5 TF-bound pREs had a H3K27me3 mark. On the other hand, 60% pREs with no TF peaks had H3K27me3 binding. Complete genomic tracks for each of the 38 TF loci are shown in Figure S5.

Overall, active marks and repressive marks were evenly represented across the 2083 pREs identified in the CRTFN (27% for each histone modification). At individual genomic loci though, there was a range of acetylated and repressed pREs (Figure 6D, S3C). Certain loci were enriched for H3K27me3 (i.e. *Emx2, Dmrt3* and Sp8) whereas others were highly enriched for H3K27ac (i.e. *NfiB, Bcl11a and Lhx2*) (Figure 6D, S3C, S4B). In general, genes with MP expression had pREs that were significantly more repressed than pREs in other genes (acetylation/methylation ratio is 0.38 vs. 2.80, p<0.05, unpaired two-tailed t-test, Figure S4B, S3C). Thus, the analysis of histone modifications in the CRTFN on purified VZ cells has provided evidence for which regulatory elements are active and repressed during cortical VZ patterning.

### Promoter-enhancer binding around 35 TFs of the CRTFN

To define promoter-enhancer units, and thereby gain information about the looping structure of the chromatin, we performed proximity ligation assisted chromatin-immunoprecipitation (PLAC)-seq on E12.5 cortices (Fang et al., 2016). We dissociated ∼6 million cortical cells, immunoprecipitated promoters using a known promoter epigenomic mark (H3K4me3), and then identified chromosomal interactions for those promoters (Juric et al., 2019). The identification of enhancer-promoter units that surpassed an FDR < 0.01 yielded a total of 39,655 interactions using 5kb bin pairs and 118,701 interactions with 10kb bin pairs. Over 90% of these interactions were located at distal loci (at least 2500 bp away from the TSS of a gene).

Enhancer-promoter loops show points of contact between TF gene promoters and multiple pREs (see *Lef1* locus, Figure 6B, S6). PLAC-Seq loops terminated at regions of chromatin accessibility, TF binding and H3K27Ac, as well as at VISTA enhancers with cortical activity (i.e. hs1050) (Figure 6B). The PLAC-Seq data allowed us to assign some of the 2083 CRTFN pREs to specific genes based on interaction between the pREs and the promoter region of genes. Within the CRTFN, we identified 619 pRE-gene interactions (using 5kb bin pairs) and 945 pRE-gene interactions (using 10kb bin pairs, Table S5). In addition, we complemented the PLAC-seq results with previously computationally derived enhancer-gene associations based on correlation between the putative enhancer activity and the expression level of the genes residing in the same topologically-associating domain (Table S5) (Osterwalder et al., 2018; ENCODE Project Consortium et al., 2020). Thus, we defined the interactome of cis-regulatory elements around the TSS for each CRTFN genomic locus.

### Epigenomic marks are good predictors of pRE activity

We tested whether chromatin state was predictive of cortical progenitor activity for pREs by examining open chromatin and histone marks on VISTA Enhancers in Table S3. As with TF binding, Pallial VISTA enhancers were more likely to have ATAC and H3K27ac peaks than Subpallial, Non-telencephalic or Inactive enhancers (Figure S4C). In contrast, H3K27me3 was not enriched on cortical VISTA enhancers compared to other enhancers (active in other tissues or inactive).

Next, we tested the enhancer activity of additional pREs in the CRTFN by selecting a set of loci with 3-5 TF binding and ATAC epigenomic marks. We tested 18 pREs using a transgenic reporter assay at E12.5 (Visel et al., 2013). We found that 15/18 pREs had positive enhancer activity at E12.5 with at least 3 embryos showing the same pattern of activity (Table S6). Of those, 5/15 had activity in the cortex and an additional 8/15 had activity in surrounding regions such as the pretectum or subpallium (Table S6). Two of these enhancers are shown in Figure 6E and displayed activity in the developing DP (mm1787 near *Etv5*) as well as the CDP and MP (mm1789 near *Npas3*) (Figure 6E). Thus, integrating TF and histone ChIP with ATAC data provides an efficient approach to identify REs that are active in cortical progenitors.

### Identification of pREs that are sensitive to changes in cortical patterning in Emx2-/-, Nr2f1-/- and Pax6-/-

Having identified 2083 CRTFN pREs in cortical progenitor cells, we next sought evidence for which of these may participate in regulating regional patterning. To this end, we looked for changes in pRE chromatin state in *Emx2*^*-/-*^, *Nr2f1*^*-/-*^ and *Pax6*^*-/-*^, an approach we previously applied in *Nkx2-1* and *Dlx1/2* mutants (Sandberg et al., 2016; Lindtner et al., 2019). We used the Flash-Tag/FACS method to purify cortical VZ progenitors from E12.5 WT and mutant littermates. After FACs sorting, we conducted ATAC-seq and histone ChIP-seq (H3K27ac and H3K27me3). We annotated the data as: Gain (gain of enrichment in a peak in the mutant), Loss or No change In the CRTFN, we found 204 pREs (12% of pREs) that were affected in the mutant cortex (“differential pREs”) (Table S7).

Of these 204 differential pREs, we could associate over 80% with a TF gene promoter either by PLAC-seq or by computational analysis (Table S7). The annotated changes in epigenetic state of differential pREs for each mutant are presented in Figure 7G. *Pax6*^-/-^ showed a 69% loss of H3K27ac marks (81/118) and no increases; and a balanced loss and gain of H3K27me3 (12% and 11%, respectively). Consistent with histone changes, loss of chromatin accessibility (25%) occurred more frequently than an increase in chromatin accessibility (2%). *Emx2*^-/-^ showed a 44% loss and a 17% gain of H3K27ac; and a 42% loss and 47% gain of H3K27me3. Chromatin accessibility was more often lost (25%) than gained (7%). Finally, *Nr2f1*^-/-^ showed a 45% loss of H3K27ac and an 11% loss of H3K27me3, and no gains in either mark. Chromatin accessibility only showed a change in 3% of peaks (Figure 7E).

**Figure 7.**
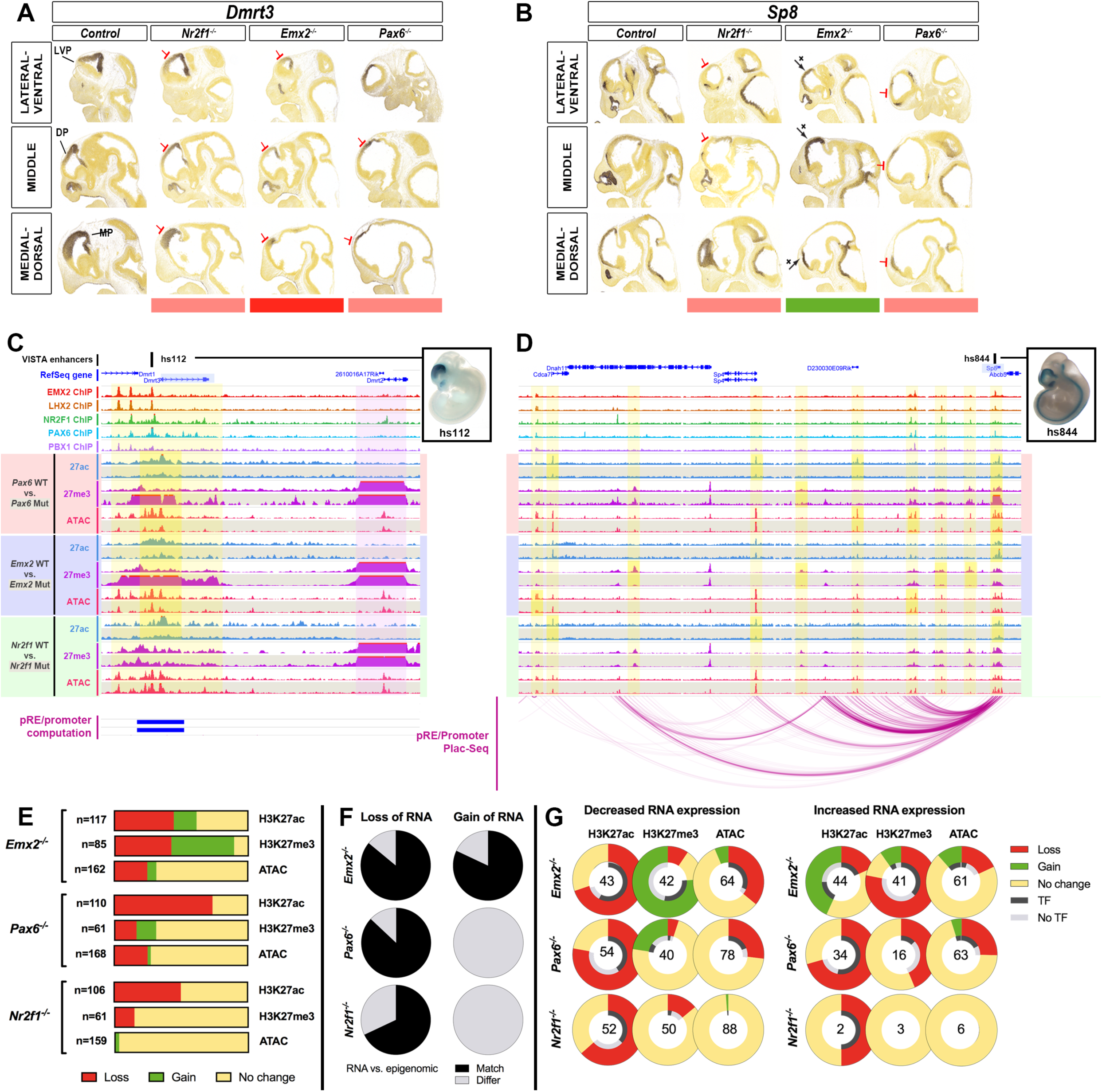
pREs show epigenetic changes in *Emx2*^-/-^, *Nr2f1*^*-/-*^, and *Pax6*^*-/-*^. (A-B) ISH for *Dmrt3* and *Sp8* in WT, *Nr2f1*^*-/-*^, *Emx2*^*-/-*^ and *Pax6*^*-/-*^ at E11.5. Red and green bars below indicate decrease and increase in expression. (C-D) TF ChIPs and VZ epigenomic marks (H3K27ac, H3K27me3, ATAC-seq) at *Dmrt3* and *Sp8* loci at E12.5 in Wild-Type, *Pax6*^*-/-*^ (red*), Emx2*^*-/-*^ (blue) and *Nr2f1*^*-/-*^ (green). WT tracks (shaded white) are above mutant tracks (shaded grey). pRE-promoter interactions are derived from PLAC-seq (purple arcs below *Sp8* locus) and computationally (blue boxes below *Dmrt3* locus). Two cortical VISTA enhancers (hs112 and hs844) are shown above the genomic loci (black squares); pictures of embryo wholemounts are to the right and show their cortical activity (blue stain). Yellow vertical bars highlight regions of differential epigenomic marks in mutants, and a purple vertical bar highlights *Dmrt2*, a region next to *Dmrt3* that is unchanged. (E)CRTFN pREs with differential enrichment of epigenomic marks in the *Emx2*^*-/-*^, *Pax6*^*-/-*^ and *Nr2f1*^*-/-*^ were tabulated. Loss (red), Gain (green) and No change (yellow) were recorded for individual pREs and tabulated across all peaks for individual epigenomic marks. (F)Proportion of loci showing a match (black) or a mismatch (grey) between the RNA ISH analysis and the differential H3K27ac/H3K27me3 ratio in the TF locus, in *Emx2*^*-/-*^, *Pax6*^*-/-*^ and *Nr2f1*^*-/-*^. For *Nr2f1*^*-/-*^, there is not enough information to make comparisons (only one TF that shows a gain in RNA expression; that locus has 2 pREs). (G)pREs in TFs loci were classified as having either a decrease, an increase or no change in RNA expression in the *Emx2*^*-/-*^, *Pax6*^*-/-*^ and *Nr2f1*^*-/-*^. We show the changes in epigenomic marks in the TF genes’ interactome according to whether the TF gene showed increased or decreased RNA expression. The percent change compared to wild-type littermates is plotted in the outer ring (Loss in red; No change in yellow; Gain in green). For pREs that changed we assessed whether we found a TF peak at that pRE (i.e. in the case of *Pax6*^*-/-*^, we looked for a PAX6 ChIP peak). The proportion of differential pREs with or without the cognate TF binding ChIP-seq reads is presented in the inner ring (Black for presence of the TF binding peak; Grey for the absence of the TF binding peak). Numbers of differential pREs assessed for each TF gene locus are shown in the center of the rings.

We next checked whether these epigenomic changes were consistent with changes in RNA expression within the CRTFN (Figure 7F). In genes exhibiting a decrease in RNA expression, we expect a loss of H3K27ac and a gain of H3K27me3 (Amberg et al., 2019). In *Emx2*^*-/-*^, 14 TFs had reduced RNA expression and 86% of these genomic loci exhibited an overall loss of active and/or gain of repressive marks on pREs (Table S7). In *Pax6*^*-/-*^, 15 TF genes had a reduction in RNA expression and 87% of these genomic loci showed an overall loss of active and/or gain of repressive marks. Finally, in *Nr2f1*^*-/-*^, 19 TF genes had a reduction in RNA expression and 13 (68%) of these genomic loci showed an overall loss of active marks at pREs. This indicates that, for a given TF gene, reduction of RNA expression in a mutant is correlated with a reduction of the H3K27ac/H3K27me3 ratio in the pREs of that locus.

In contrast, for genes showing an increase in RNA expression, we expect a loss of H3K27me3 and/or a gain of H3K27ac (increased H3K27ac/H3K27me3 ratio). *Emx2*^*-/-*^ met this expectation; 9/11 TF genes that had increased RNA expression, also had pREs that lost H3K27me3 and/or gained H3K27ac. On the other hand, in *Pax6*^*-/-*^ increased RNA expression was not associated with an increased H3K27ac/H3K27me3 ratio.

The summary of changes in epigenomic profiles for pREs based on changes in ISH RNA expression in *Emx2*^*-/-*^, *Nr2f1*^*-/-*^ and *Pax6*^*-/-*^ are presented in Figure 7G. The schema is organized according to Loss, Gain and No change in RNA expression in the three mutants. The outer ring depicts percent changes for each epigenomic mark in pREs in the mutants (Loss, No change, Gain). The inner ring depicts the presence (Black) or absence (Grey) of the cognate TF ChIP binding peak (i.e. PAX6 in *Pax6*^*-/-*^). The number in the center of the rings represents the number of pREs investigated. Overall, we suggest that loss of H3K27ac is associated with reduced RNA expression and with TF protein binding to pREs in the *Emx2*^*-/-*^, *Nr2f1*^-/-^ and *Pax6*^*-/-*^.

Next we show two gene loci (*Dmrt3* and *Sp8*) in Figure 7 (C-D) with the cognate change in RNA expression presented in Figure 7 (A-B) (see Figure S5 for all 38 TF loci in the CRTFN). In the genomic loci of these 2 TFs, we highlighted pREs that showed epigenomic changes in yellow. *Dmrt3 RNA* expression is downregulated in all mutants. At the *Dmrt3* locus, hs112 showed a decreased H3K27ac/H3K27me3 ratio and a loss of chromatin accessibility in both *Emx2*^*-/-*^ and *Pax6*^*-/-*^. *Sp8* is downregulated in *Pax6*^-/-^ but increased in *Emx*2^-/-^. pRE epigenetic changes were in synchrony with this. For instance, the H3K27ac/H3K27me3 ratio on hs844 was reduced in *Pax6*^*-/-*^ and increased in *Emx2*^*-/-*^. Other pREs around the *Sp8* locus showed epigenetic changes in only one or two mutants and were unchanged in the other knockouts.

We identified 20-120 pREs per locus with 10% showing epigenetic changes in the TF mutants. Understanding how individual pREs regulate transcription at a single gene locus will require investigating these pREs individually and in combination across normal and abnormal development.

## DISCUSSION

We have identified the key elements (transcription factors and target genomic regulatory elements) in a transcriptional network that regulates early steps in cortical regional patterning. This Cortical Regionalization TF network (CRTFN) is comprised of TFs and their pREs. We tested the validity of this network by characterizing the impact of genetic ablation of three key members (*Emx2, Nr2f1, Pax6*) on the expression of TF genes and the epigenomic state of pREs. Within the CRTFN, 30/31 of the TF genes were sensitive to genetic ablation of at least one of these three TFs. pREs were identified using TF ChIP-seq for EMX2, LHX2, NR2F1, PAX6 and PBX1, and epigenomic assays for chromatin accessibility, active pREs (H3K27ac) and repressive pREs (H3K27me3). pREs were further assessed by PLAC-seq defined promoter-enhancer looping or computationally derived enhancer-gene associations. We found that pREs with combinatorial binding of different TFs were enriched at enhancers displaying cortical activity (VISTA enhancers), and that their binding was correlated with activity in the LVP and rostrocaudal (RC) gradients, and anti-correlated with activity in the MP and dorsoventral (DV) gradients. We also showed that five novel CRTFN pREs had cortical activity in transient transgenic assays. ∼10% pREs were sensitive to loss of EMX2, NR2F1 or PAX6; these showed a change in the H3K27ac/H3K27me3 ratio. Taken together, our integrated approach has identified a TF network involved in regional patterning of the cortex.

### Defining a CRTFN

We carried out an unbiased screen to discover TFs that have informative patterns in the E11.5 pallium using the Developing Mouse Brain Atlas (http://developingmouse.brain-map.org). TFs at E11.5 were expressed either in gradients (∼70) or homogeneously across cortical regions (∼150). We did not find TFs with sharp intrapallial boundaries except for 6 whose expression was restricted to the MP. This suggests that cortical areas do not originate from progenitor domains with spatially restricted TF expression, but rather from a process that integrates information from TF gradients to create sharp boundaries.

We focused our analysis on testing the sensitivity of 31 CRTFN TFs with graded expression patterns in *Emx2*^*-/-*^, *Nr2f1*^*-/-*^ *and Pax6*^*-/-*^. Caudorostral (CR) TFs showed decreased expression in *Emx2*^*-/-*^ and *Nr2f1*^*-/-*^; whereas RC TFs showed decreased expression in *Nr2f1*^*-/-*^ and *Pax6*^*-/-*^. Patterning TFs often act in concert to regulate regional patterning of pallial subregions (Muzio et al., 2002a). Here, we present evidence that *Nr2f1* and *Pax6* act together to co-regulate the development of the rostroventral cortex. We found that *Pax6* promotes *Nr2f1/2* expression and that, together, these TFs co-regulate other rostroventral TFs including *Lmo3* and *Npas3* which in turn plays a role in patterning the rostral piriform cortex and adjacent structures. Of note, the rostral subplate is also reduced in the *Pax6*^*-/-*^, *Nr2f1/2 cKO* and *Npas3*^-/-^ (Fig. 3), suggesting that its development is linked to the rostroventral cortex (Saito et al, 2019).

Thus, we defined CRTFN TFs that are likely to play a role in cortical regional patterning. Further, it enabled us to identify a *Pax6-Nr2f1-Npas3-Lmo3* pathway that regulates patterning of the rostroventral cortex.

### The epigenome of CRTFN pREs predicts their spatial gradients of cortical activity

Having defined the key TFs playing a role in cortical patterning, we aimed to uncover the regulatory network governing the expression of TF gene gradients. To this end, we identified putative regulatory elements (pREs) of the CRTFN.

Combinatorial binding of EMX2, LHX2, NR2F1, PAX6 and PBX1 was a strong predictor of pRE activity in the developing cortex (Figure 5C). Biochemical and computational chromatin conformation analysis provided evidence that at least 45% of pREs interacted with a given TF gene’s promoter, and defined the cis-regulatory interactome for each CRTFN gene. 94% of cortical Vista enhancers in the CRTFN had binding of at least one TF. Moreover, combinatorial binding of TFs was strongly enriched in Pallial Vista enhancers compared to Subpallial, Non-telencephalic and Inactive enhancers.

Here we uncovered some of the transcriptional logic underlying differences in enhancer regional activity using two different methods: 1) degree of TF binding (EMX2, LHX2, NR2F1, PAX6 and PBX1) on VISTA enhancers with regional activity; 2) an unbiased computational approach (Fig. 5). We found that enhancers active in the most dorsal caudal pallium (MP) had the least binding, whereas enhancers active in the most rostral ventral pallium (LVP) had the most binding. This bias is not due to expression of the TFs (*Emx2, Lhx2, Nr2f1* and *Pax6* are expressed in the MP), nor is it entirely due to a bias in cell numbers since LVP and MP are both small structures in comparison to the DP, which had a level of TF binding intermediate to that of the LVP and MP. We propose that there is a distinct set of MP TFs, such as those found in our screen, *Id3* and *Lmx1a*, that promote the activity of MP enhancers (Figure 1; Fregoso et al., 2019). Thus, we have shown that the combination of EMX2, LHX2, NR2F1, PAX6 and PBX1 binding in pREs provided a signature that was predictive of pRE regional activity and cortical gradients.

Towards establishing an epigenetic metric for a gene’s activity, we calculated the H3K27Ac/H3K27me3 ratio within the interactomes’ of each CRTFN TF. We found that pREs within the genetic locus of TFs with expression in the MP had a lower H3K27Ac/H3K27me3 ratio than that of non-MP TF gene loci (Figure S4). We suggest that pREs of MP genes are repressed in the DP and LVP, and we predict that a focused analysis on the VZ of the MP would find a H3K27Ac/H3K27me3 > 1.

To gain insights into the function of the pREs, we assessed their epigenomic state. We restricted our study to VZ cells to bypass issues relating to cellular heterogeneity. We found that VZ ATAC and H3K27ac peaks were proportionally enriched at pREs that have increased co-binding by TFs (Figure S4C). This indicates that pREs with combinatorial TF binding are accessible in the VZ and are likely to be active. We estimated that CRTFN gene loci have between 20-100 pREs, thus underscoring the complexity of regulatory landscapes (Table S2)

This analysis identified ∼2000 pREs in the CRTFN based upon their chromatin state and binding by TFs important for cortical patterning. Co-binding of multiple TFs, as well as the presence of ATAC and H3K27ac marks, is correlated with enhancer activity. Moreover, our results suggest that binding of TFs can predict enhancer spatial activity gradients, thus uncovering a transcriptional logic to cortical regional patterning.

### pREs of the CRTFN respond to loss of EMX2, NR2F1 or PAX6 through changes of H3K27ac/H3K27me3 ratios rather than changes of chromatin accessibility

At the gene locus level, transcriptional regulation requires the integration of RE function. In the CRTFN each TF gene can have 20-100 pREs. Our analysis uncovered complexity in the properties and regulation of individual pREs within the CRTFN when we probed their epigenomic profiles in *Emx2*^*-/-*^, *Nr2f1*^*-/-*^ and *Pax6*^*-/-*^.

Enhancers are spatial and temporal integrators of combinatorial TF binding and chromatin state (Nord et al., 2015). While the majority of pREs did not exhibit epigenetic changes in the three mutants, we did find 204 pREs that were differentially regulated, of which >80% were linked to the TSS of the TF gene (Table S7). This suggests that these pREs are responsive to ablation of these specific TFs and likely important for regional patterning of the cortical neuroepithelium.

Loss of EMX2, NR2F1 and PAX6 had different effects on distinct pREs. In some cases, changes in the epigenetic state of a pRE was similar in two or three mutants (38%). For instance, *hs112* in *Dmrt3* showed loss of H3K27ac in the three mutants, gain of H3K27me3 in *Pax6*^*-/-*^ *and Emx2*^*-/-*^ and loss of chromatin accessibility in *Pax6*^*-/-*^ (Figure 7). Other pREs (∼9%) responded differently in different mutants. For instance, *hs844* in the *Sp8* locus showed a reduced H3K27ac/H3K27me3 ratio in the *Pax6*^*-/-*^ but an increased ratio in the *Emx2*^*-/-*^. In 52% of occurrences, pREs showed changes in only one TF mutant, reflecting the specificity of pRE regulation. While histone modifications were sensitive marks of pRE epigenetic changes, the ATAC-seq assay of chromatin accessibility showed few changes in all three mutants (Figure 7G). Thus, while ATAC-seq was effective at identifying active pREs (Figure 6C), it was not an effective tool to assess which were modified in the VZ of *Emx2*^*- /-*^, *Nr2f1*^*-/-*^ and *Pax6*^*-/-*^.

At the gene locus level, each TF gene had between 1-13 differentially regulated pREs in the three mutants. In 93% of cases, all differentially regulated pREs in the same locus showed a similar change in their H3K27ac/H3K27me3 ratio in a given mutant, such as at the *Dmrt3* locus in *Pax6*^*-/-*^ and *Emx2*^*-/-*^. Perhaps, some of these pREs can compensate for one another making their regulatory programs more robust to mutations, such as in the *Arx* loci (Dickel et al., 2018). However, as for *Arx* enhancers, such redundant enhancers could also have distinct functions.

Integration of discrete RE function across a locus leads to overall transcriptional patterns of the target genes. We found that genes showing reduced cortical RNA had coherent changes in epigenomic marks in the pREs of mutants (Figure 7H,I). Likewise, genes showing increased RNA had coherent changes in epigenomic marks in the *Emx2*^*-/-*^ mouse but surprisingly not in the *Pax6*^*-/-*^. Understanding why *Pax6*^*-/-*^ showed increased RNA may require probing different epigenomic marks and/or investigating indirect mechanisms of transcriptional activation. The most pronounced epigenomic change that we detected was the loss of H3K27ac marks in *Pax6*^*-*/-^ (Figure 7G). PAX6 binds several chromatin modifiers, including p300, a transcriptional co-activator and histone acetyltransferase (Hussain and Habener, 1999); perhaps this accounts for the reduction in the H3K27ac marks in *Pax6*^*-*/-^. Thus, further integration of TFs and chromatin regulators will be crucial in understanding the epigenetic code that regulates cortical development.

Around 10% of CRTFN pREs were epigenetically changed in *Emx2*^*-/-*^, *Nr2f1*^*-/-*^ and *Pax6*^*-/-*^. These were enriched for promoter looping. We propose that these pREs are the core set of enhancers regulating cortical patterning.

### Propagating the ventricular zone protomap into regionalization of the cortical laminae

Here we define TFs within the CRTFN that exhibit regional gradients in the developing cortical neuroepithelium. We demonstrate via genomic interaction mapping and genetic knockout that expression patterns of these TFs are regulated, at least in part, via the interaction of specific TFs with target REs. Considering the lack of TFs that are restricted to specific cortical VZ regions, pREs must encode regional activity via integrating the effects of combinatorial TF binding. Based on this, we hypothesize that the VZ enhancers identified here may serve as a protomap of cortical regions by integrating TF signalling. We previously postulated that enhancers serve as protein-binding modules that translate gradients of TFs in cortical progenitors into region-specific expression in cortical neurons (Pattabiraman et al. 2014; Ypsilanti et al, 2016). Our hypothesis is that information encoded in the VZ by gradients of TFs will be encoded in the regulatory elements of genes important in cell specification. Along these lines, *Pax6* was shown to control regionalization through a downstream cascade beginning with the induction of *Tbr2* in the intermediate progenitors and propagated by *Tbr1* to the cortical plate (Bedogni et al., 2010; Elsen et al., 2013). Likewise, *Pbx1, Nr2f1 and Lhx2* function in immature cortical neurons to control cortical regionalization (Golonzhka et al, 2015, Alfano et al, 2014). Moreover, at early postnatal stages, refinement of the somatosensory cortical neurons (co-expressing *Ctip2* and *Satb2*) was found to be regulated by the modulation of epigenetic mechanisms in a time and area-specific manner by *Lmo4* (Harb et al, 2016). Here, we restricted our focus to studying the transcriptional control of patterning in the VZ. Future work will further elucidate the TF network (TFs and cis-regulatory elements) that plays a role in the propagation of the VZ protomap to the overlying cortical regions.

## SUPPLEMENTARY FIGURE LEGENDS

**Figure S1. Characterization of EMX2, LHX2, NR2F1, PAX6, and PBX1 combinatorial binding**

(A)ISH cortical expression gradients of TFs used for ChIP at E11.5. *Lhx2* has a caudorostral (CR) gradient. *Emx2 has a CR and dorsoventral (DV) gradient. Nr2f1* has a CR and ventrodorsal (VD) gradient. *Pax6* and *Pbx1* have a rostrocaudal (RC) and VD gradient.

(B)Proximal (near promoter) vs. Distal binding for each TF ChIP-seq

(C)Heatmap showing pairwise Pearson Correlation for genome-wide coverage values for each TF ChIP-seq replicate.

(D)Enrichment of functional annotation terms (GO) for genomic loci showing combinatorial binding of 5 TFs by ChIP-seq.

(E) Relative motif enrichment for primary binding DNA motifs of EMX2, LHX2, NR2F1, PAX6 and PBX1 ChIP-seq across all pREs.

(F) Motif enrichment of all *de novo* motifs in pREs of the TF ChIPs.

(G) Counts of distance from TSS for loci showing combinatorial binding of 5TFs by ChIP-seq.

(H)Non-TF genes important in cortical development showing co-binding of 5TFs by ChIP-seq. Light green are genes with known functions during cortical development; Dark green are chromatin modifiers; Orange are TFs with expression in other brain regions.

(I)Analysis showing percentage of peaks with motif enrichment of primary binding motifs (described in Figure 4C) for each unique or combinatorial binding of TFs by ChIP-seq. Abbreviations: Hbox: homeobox.

**Figure S2. Using VISTA enhancers to model how combinatorial binding of EMX2, LHX2, NR2F1, PAX6, and PBX1 predicts cortical activity and graded expression in the developing pallial VZ**.

(A)Plot showing how likely VISTA enhancer loci bound by the TFs are forebrain-active as opposed to active in other brain regions and elsewhere (heart as a surrogate). Main plot shows the frequency of occurrence, while the upper one depicts the comparison of mean number of enhancers hit by sampling, showing significant difference among different enhancer classes (p < 0.01).

(B)Examples of cortical VISTA enhancers with graded patterns of activity. The wholemounts and sections for each example are placed along the central grid according to their annotated gradient of activity in the developing pallium. hs1035 has a DV and RC gradient; hs798 has a DV and CR gradient; hs636 has a RC and VD gradient; and finally, hs1172 has a CR and VD gradient.

(C)Correlalogram showing the expansion of the modeling presented in Fig. 4D showing the different combinations of TF binding and their predictions of gradients of activity (p < 0.01).

**Figure S3. Epigenomic profiling of CRTFN genes (Cortical Regionalization TF Network) in the cortical VZ**.

(A) Histology of Flash-Tag staining of the cortical VZ at E12.5; note that the Flash-Tag labeled VZ cells do not overlap with TBR2 immunostained SVZ progenitors.

(B)Example FACs plots showing data for Flash-Tag (CFSE) positive progenitor cells prepared from the E12.5 cortex. The histogram of FITC counts shows a bi-modal FITC negative population (VZ^-^) and a FITC positive population (VZ^+^). The dot plot depicting FSC vs. FITC shows the gating which was used to collect the FITC^+^ (VZ^+^) population. Black bar above VZ^+^ correspond to the cells collected for further analysis.

(C) Overview of epigenomic enrichment in pREs with differential combinatorial TF binding (0-5 TFs) for CRTFN TF gene loci. Number of pREs at each locus are in brackets (n=#pREs). Percentage of total pREs are indicated by a teal histogram bar. Percentage of pREs with the following epigenetic marks are indicated by the following colors: ATAC = yellow; H3K27ac = dark green; H3K27me3 = red. See Figure S4A which summarizes this data for each TF locus.

**Figure S4. Epigenomic profiling of the cortical VZ shows a diminished H3K27ac/H3K27me3 ratio over pREs for MP TFs**

(A) Table showing percentage enrichment of TF binding and epigenomic marks of pREs at each locus of the Cortical Regionalization TF Network. A H3K27ac/H3K27me3 ratio is calculated for each locus.

(B) Statistical analysis shows that the H3K27ac/H3K27me3 ratio is significantly lower for pREs in genomic loci of TFs with MP expression (red) compared to pREs for the non-MP TFs (purple) (p<0.05, unpaired two-tailed t-test).

(C)Plot of genomic features (TF binding, Epigenomic marks, H3K27ac, H3K27me3, ATAC) over VISTA enhancers that have pallial (dark green), subpallial (light green), non-telencephalic (yellow) and no activity (red) at E11.5.

**Figure S5. CRTFN transcriptional network in *Emx2***^***-/-***^, ***Nr2f1***^***-/-***^ **and *Pax6***^***-/-***^

For each of the 38 TFs in the CRTFN, we show:

-ISH of TF gene in wild-type and mutant mice at E11.5.

-pallial VISTA enhancer wholemount (and sections if available) showing activity in the cortex at E11.5.

-Table with a comprehensive list of pREs that shows differential regulation in *Pax6*^*-/-*^, *Emx2*^*-/-*^ and *Nr2f1*^*-/-*^ around the interactome of the relevant TF. Each pRE in this table is highlighted by a yellow column in the genome tracks below. The highlighted columns have orange numbers matched to the row number in this table.

-TF’s genomic loci showing:

1. RefSeq genes around the TF gene (highlighted in purple).
2. Genomic position of VISTA enhancers with cortical activity (Green), activity in other regions (Blue) or no activity (Red) at E11.5.
3. Computationally derived enhancer-gene associations based on correlation between the putative enhancer activity (using Encode data: Comp_Encode or data generate here: Comp_MO) and the expression level of the genes residing in the same topologically-associating domain.
4. TF ChIP binding for EMX2, LHX2, NR2F1, PAX6 and PBX1; replicates are shown.
5. Epigenomic marks in VZ of Wild-Type (Shaded in white) and *Pax6*^*-/-*^, *Emx2*^*-/-*^ and *Nr2f1*^*-/-*^ littermates (shaded in grey). For each experiment, H3K27ac tracks are in blue, H3K27me3 tracks are in purple, and ATAC-seq tracks are in red. Replicates are shown.
6. pRE-promoter interactions derived from PLAC-seq are shown by arcs (magenta) using two statistical stringencies (5k and 10k).

pREs with differential epigenomic marks in any one mutant are highlighted in yellow columns. These columns are labelled by orange numbers that correspond to the rows in the table above.

**Figure S6**.

Bar plot showing proportion of pREs for CRTFN genomic loci that are sensitive (light green) or insensitive (dark green) to the effects of *Pax6*^*-/-*^, *Emx2*^*-/-*^ and *Nr2f1*^*-/-*^.

## TABLE LEGENDS

**Table S1**: Annotation of TF expression in pallium at E11.5. Expression levels were assessed on a scale of 0-5 for density (D) and intensity (I) of ISH staining in subregions of the pallium: LVP, RDP, CDP, MP. When possible, VZ/SVZ and Mantle zone expression were assessed (blank means levels of staining in these regions could not be assessed).

**Table S2**: Genomic coordinates of CRTFN pRES annotated for TF ChIP-seq peaks (EMX2, LHX2, NR2F1, PAX6 and PBX1), histone ChIP-seq peaks (H3K27ac, H3K27me3) and ATAC-seq peaks. Each row is a pRE. Column A is the TF gene and column B-D is the interactome of a given TF gene. Column H is the size of the pRE. Numbers in column I are the number of TFs co-bound to a given pRE. Numbers in columns (J-Q) indicates how many replicates have a peak. Number of replicates for each ChIP-seq/ATAC-seq is indicated in parenthesis in the title of column. Column R annotates the presence of a VISTA enhancer.

**Table S3**: Classification of CRTFN VISTA enhancers as Pallial, Subpallial, Non-telencephalic and Inactive at E11.5. Table shows VISTA enhancer name, genomic coordinates, region of activity and VISTA annotation of other regions of activity, TF and histone ChIP-seq peaks. Table includes: enhancer name, genomic coordinates, size of enhancer, relevant region of activity of enhancer (P for Pallial, SP for Subpallial, N for Non-telencephalic and I for Inactive), other regions of activity of enhancer, TF ChIP-seq, WT histone marks (H3K27ac and H3K27me3) and ATAC-seq peaks. In rows, x indicates presence of that particular feature.

**Table S4:** Classification of gradients of activity in CRTFN VISTA enhancers. Table shows VISTA enhancer name, genomic coordinates, flanking genes, cortical gradient of activity (RC = rostrocaudal, CR = caudorostral, DV = dorsoventral, VD = ventrodorsal) and a detailed analysis of their subregional cortical expression of activity when sections were available (columns J-R). Number 1 in column indicates presence of that particular feature.

**Table S5:** PLAC-seq and computational pREs-promoter interactions in the interactome of CRTFN genes. Tab 1 maps the interactions derived from PLAC-seq in CRTFN interactomes using a 5k stringency. Computationally derived enhancer-gene associations are also indicated. This computational analysis is based on correlation between the putative enhancer activity (using Encode data: Comp_Encode or data generated here: Comp_MO) and the expression level of the genes residing in the same topologically-associating domain. Tab 2 is the same as Tab 1 with a stringency for PLAC-seq of 10k. Footnotes and abbreviations for Tab 1-2 is in Tab 3.

**Table S6:** Coordinates and activity annotations of newly tested VISTA enhancers. Each row is a newly tested VISTA enhancer and includes information about its coordinates, its activity overall, its cortical activity (column F), annotations of other regions of activity, annotations for presence of TF ChIP-seq, histone and ATAC ChIP-seq peaks, and pRE-promoter interactions. x and xx indicate presence of peaks in 1 or 2 replicates respectively.

**Table S7:** Coordinates and annotations (Gain, Loss, No Change) of pREs that show changes in histone marks and/or chromatin accessibility in *Pax6*^*-/-*^, *Emx2*^*-/-*^ and *Nr2f1*^*-/-*^. This table shows for each interactome: coordinates of sensitive pREs (column B); primary DNA binding motifs (column C-G); TF ChIP-Seq (column H-L); Gain, Loss and No Change in Histone marks and ATAC Seq in Pax6, Emx2 and Nr2f1 mutants (Columns M-U); Plac-Seq and computation pRE-promoter binding (Column V-W); Genomic features of a given pRE.

## ACKNOWLEDGMENTS

This work was supported by grants to JLRR from Nina Ireland, the National Institute of Neurological Disorders and Stroke (NINDS R01 NS34661), and the National Institute of Mental Health (NIMH R37 MH049428). ARY was supported by a postdoctoral fellowship from the Fondation Fyssen (Paris, France). ASN and RCP were supported by National Institute of General Medical Sciences (NIGMS R35 GM119831). Mutant mice were given by M and S Tsai (*Nr2f*) and S McKnight (*Npas3*). RC-P was supported by a Science without Borders Fellowship from CNPq (Brazil). MH and IJ are partially supported by NIH grant U54DK107977. This study was supported in part by the HDFCCC Laboratory for Cell Analysis Shared Resource Facility through a grant from NIH (P30CA082103). I.B. is funded through an Imperial College Research Fellowship. Research conducted at E.O. Lawrence Berkeley National Laboratory was supported by NIH grant R01HG003988 (to LAP) and performed under Department of Energy Contract DE-AC02-05CH11231, University of California.

## AUTHOR CONTRIBUTIONS

## Conceptualization

ARY, KP, JLRR

## Methodology and Investigation

ARY (TF ChIP-seq, Histone ChIP-seq, ATAC-seq, ISH, bioinformatics), KP (ISH), RC-P (bioinformatics, modeling), SL (TF ChIP-seq), OG (TF ChIP-seq), DED, LAP, and AV (enhancers in mice), ARY, IJ and YS (PLAC-seq), AA, IJ and MH (MAPS-interaction calling), IB and MH (Computation ??), LT and HZ (ISH of 31TFs in patterning mutants), Ke-Tang (Nr2f2 flox allele).

Writing – Original Draft, AY, KP, RC-P, ASN and JLRR; Writing – Review & Editing, AY, KP, RC-P, DED, ASN, IB and JLRR

Funding Acquisition: JLRR and ASN Supervision: JLRR and ASN

## STAR Methods

### Key Resources Table

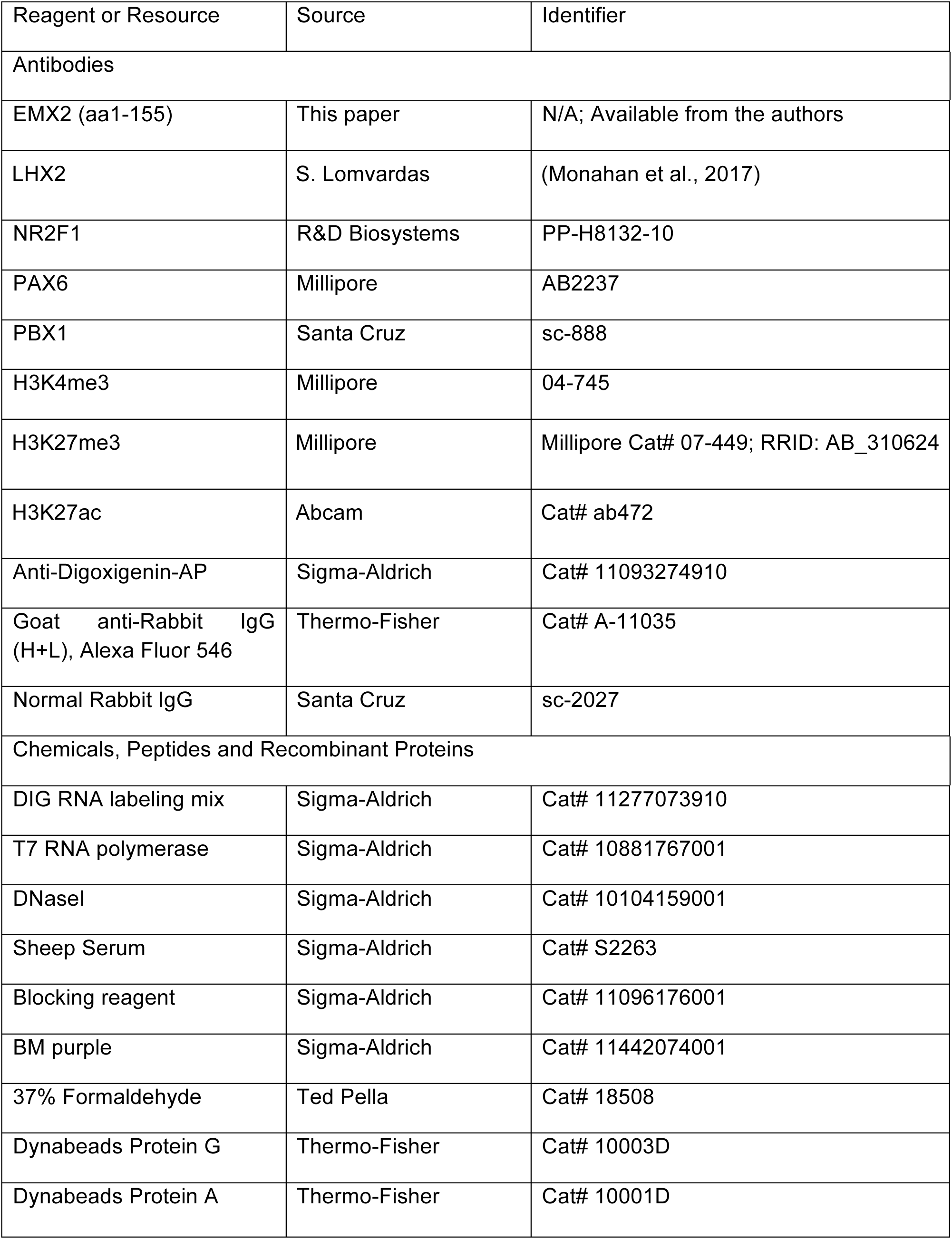
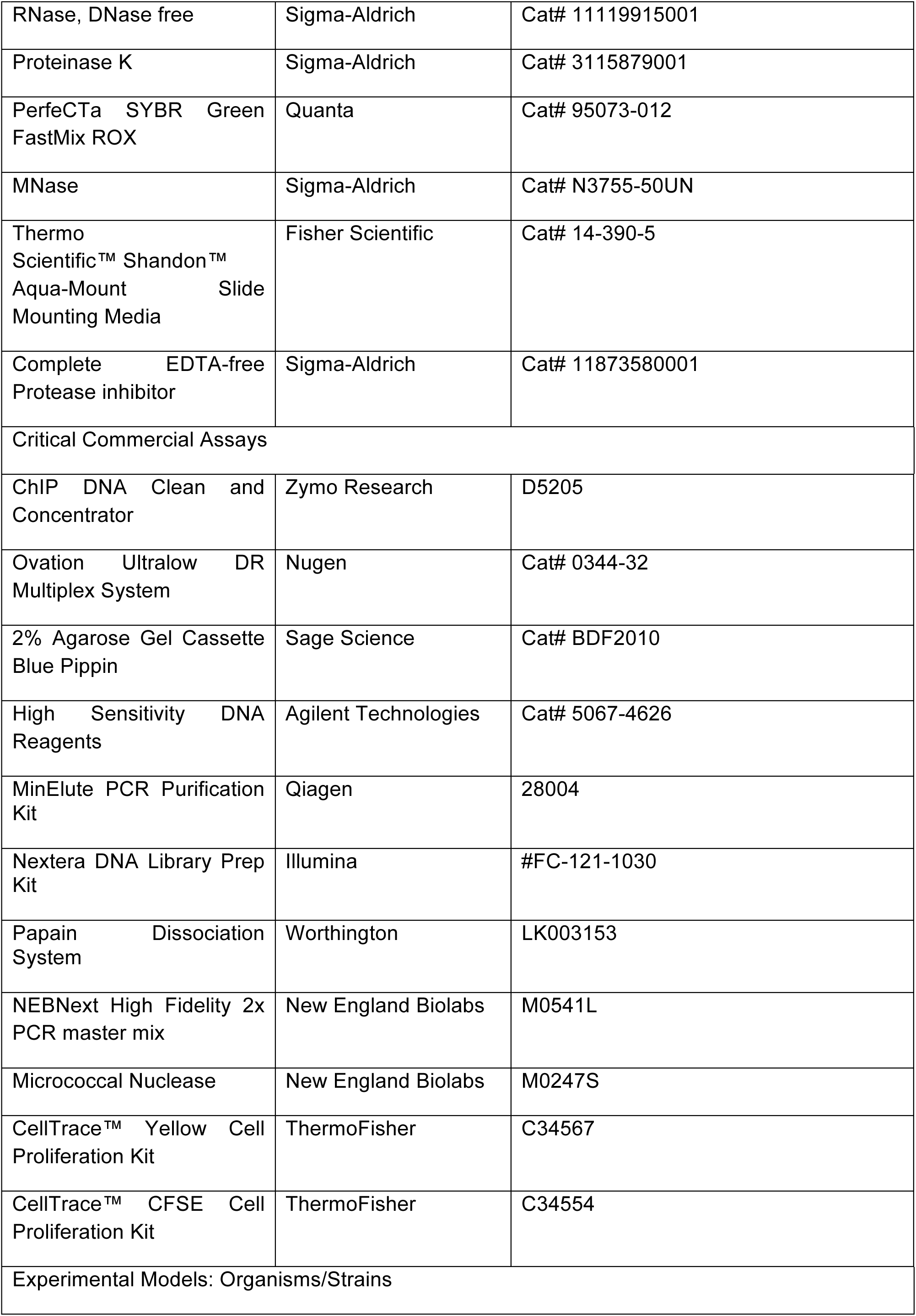
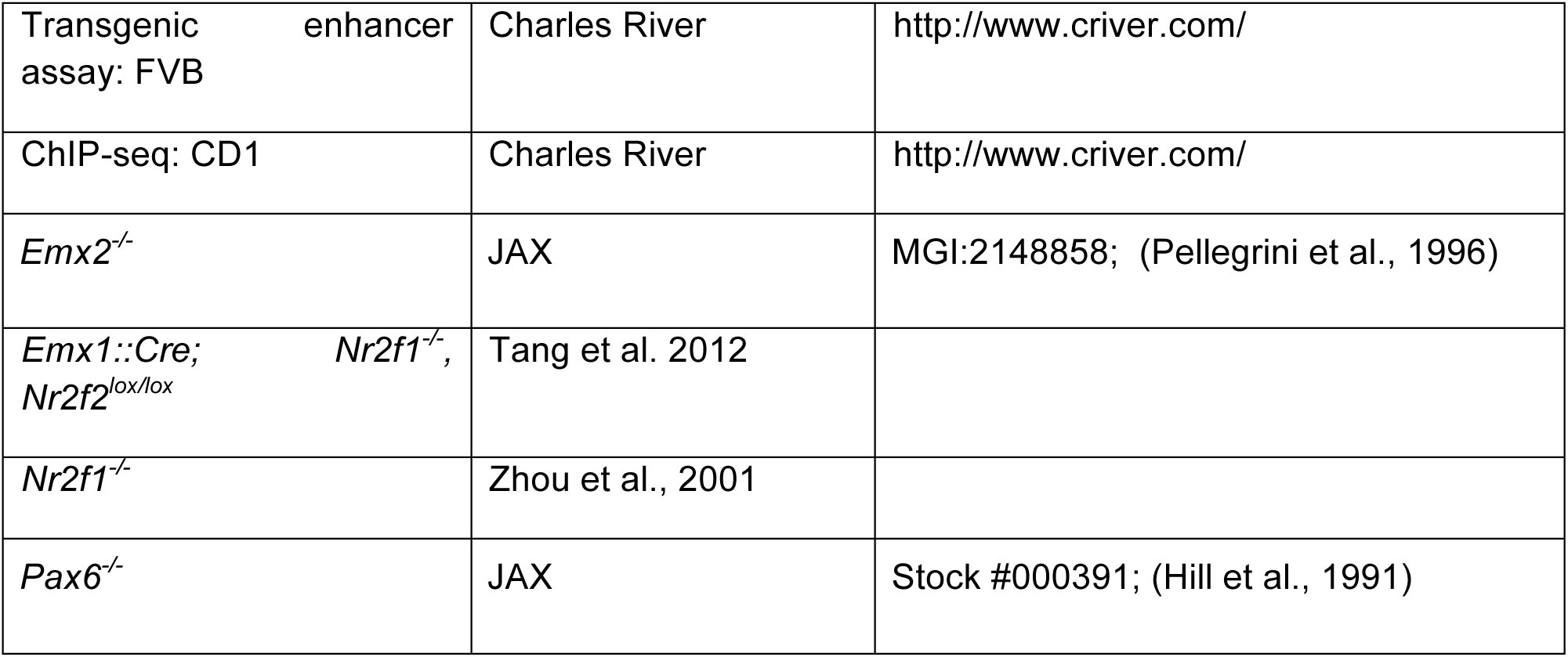

#### Contact for Reagent and Resource Sharing

Further information and requests for resources should be directed to and will be addressed by the Lead Contacts, John L. R. Rubenstein and Alex S. Nord.

### Experimental Model and Subject Details

#### Mice and genotyping

All procedures and animal care were approved and performed in accordance with National Institutes of Health and the University of California San Francisco Laboratoy Animal Research Center (LARC) guidelines. Mice strains that were used have been previously reported: *Pax6*^*Sey/Sey*^ (Hill et al., 1991), *Emx2*^*-/-*^ (Pellegrini et al., 1996), *NR2F1*^*-/-*^ (Zhou et al., 2001), *Emx1::Cre; NR2F1*^*-/-*^; *NR2F2*^*fl/fl*^ (Gorski et al., 2002; Tang et al., 2012) and *Npas3*^*-/-*^ (Erbel-Sieler et al., 2004). All mice were genotyped as previously reported.

### Method Details

#### Cortical expression analysis

The Developing Mouse Brain database of the Allen Brain Institute has generated *in situ hybridization* catalogs for hundreds of proteins in the mouse embryonic and postnatal brain. Three separate investigators annotated expression patterns of the TFs in the cortex at E11.5 by annotating both the density and the intensity of the *ISH* staining in the ventricular zone and the subventricular zone/mantle zone of the lateral ventral pallium (LVP), rostral dorsal pallium (RDP), caudal dorsal pallium (CDP) and medium pallium (MP). These annotations were then computationally mined to identify novel gradient patterns and/or regional expression.

### Histology

Brains were fixed, cryopreserved and embedded as previously described (Pattabiraman et al., 2014). After cryostat-sectioning, brain sections were stained as described (Zhao et al., 2008). In situ hybridization was performed as previously described (Pattabiraman et al., 2014). For details on how probes were generated, see ABA website.

#### Emx2 antibody production

A guinea-pig Emx2 antibody was generated by Genscript. It was raised against a peptide of the N-terminus of mouse Emx2 (aa 1-155) which specifically excluded the homeodomain of Emx2.

#### TF Chromatin Immunoprecipitation (ChIP)

Dissected cortices from embryos from multiple litters were dissociated and crosslinked at room temperature for 10 min in 1% formaldehyde (EMX2, LHX2) or 20 minutes in 1.5% formaldehyde (NR2F1, PAX6, PBX1) before being quenched for 5 mins in glycine (2.5mM) and washed gently with 1xPBS. Nuclei were extracted by lysing the fixed cells in a hypotonic buffer (50 mM Tris pH 7.5 / 0.5% NP40 / 0.25% Sodium Deoxycholate / 0.1% SDS / 150 mM NaCl). The crosslinked chromatin was sheared in dilution buffer (0.01% SDS, 1.1% Triton X-100, 1.2mM EDTA, 16.7mM Tris-HCl, pH 8.1, 167mM NaCl) using a bioruptor (Diagenode) for either 18 rounds (for samples fixed in 1% PFA) or 40 rounds (1.5% PFA) (1 round = 30 s on/45 min off at high intensity) and incubated at 4°C overnight with 5 mg of antibody or either 20× blocking peptide or a control IgG which were used as negative controls as appropriate. Protein/antibody complexes were collected using Dynabeads (20 µL protein A + 20 µL protein G) before being washed in three wash buffers: low salt wash buffer (0.1% SDS, 1% Triton X-100, 2mM EDTA, 20mM Tris-HCl, pH 8.1, 150mM NaCl), high salt wash buffer (0.1% SDS, 1% Triton X-100, 2mM EDTA, 20mM Tris-HCl, pH 8.1, 500mM NaCl); and LiCl wash buffer (0.25M LiCl, 1% IGEPAL CA630, 1% deoxycholic acid (sodium salt), 1mM EDTA, 10mM Tris, pH 8.1and TE). Chromatin was eluted in Elution buffer (1%SDS, 10mM Codium bicarbonate buffer) at 65^0^C for 10 min. Eluted chromatin was reverse crosslinked overnight at 65^0^C with NaCl (500mM), then subsequently treated with RNase (10 µg/ 200 µl reaction, 15 min at 37^▪^C) and Proteinase K (10 µg/ 200 µl reaction, 60 min at 55^▪^C). DNA was then cleaned using a ChIP DNA Clean & Concentrator kit (Zymo Research). ChIP experiments were then validated by ChIP-qPCR using a 7900HT Fast Real-Time PCR System (Applied Biosystems) with SYBR Green qPCR SuperMix (Invitrogen, Cat. No. 11760-100).

ChIP-seq libraries were prepared using the Ovation Ultralow DR Multiplex System (NuGEN). Generated libraries were size selected using Blue Pippin (centered around 300 bp), QC tested on a Bioanalyzer (Agilent) and sequenced as single-end 50 nucleotide reads on a HiSeq 4000 (Illumina) at the Center for Advanced Technology at UCSF (http://cat.ucsf.edu/).

#### ChIP-seq Computational Analysis

Clustering, base calling, and quality metrics were performed using standard Illumina software. Sequencing libraries were analyzed for overall quality and were filtered, and reads were mapped to the mouse genome (mm9) using Burrows-Wheeler Alignment (BWA) (Li et al., 2009).

#### Pairwise Pearson Correlation

Pearson correlation between aligned read counts of pairs of TFs was determined using DeepTools (Ramírez et al., 2016) to show association between TFs and between different TF replicates.

#### Motif analysis

*De novo* motif discovery and enrichment was performed using HOMER version 4.9 (Heinz et al., 2010) in the called peaks for each individual TF, using standard parameters, 300 bp up- and downstream of TF peaks. We compared the significant motifs discovered with the JASPAR (Fornes et al., 2020) database. Motif enrichment was determined for all motifs present in the HOMER known motif database with p-value < 10^−100^. We established the average motif coverage enrichment plot around 300 bp of peaks for each TF using custom R scripts.

#### Gene Ontology analysis

Gene Ontology analyses were conducted using the GREAT algorithm.

#### VISTA enhancer annotations

Relevant VISTA enhancers were annotated by at least two experts in the field. Their region of activity was ascertained on whole-mount embryos and where necessary, using sections. Gradients of activity were only defined when they were clear and when consensus was obtained between at least two observers.

#### Gradient modelling

We established the model associating TF binding, enhancer spatial activity gradients, and enhancer regional activity by determining the pairwise correlation of among the different factors. The TF binding combinations and other factors were intersected using custom R scripts, and the correlation matrix and plot with R ggcorrplot package. Only correlations with p < 0.01 was shown. Non-relevant associations were manually removed.

#### Histone mark ChIP

We used Cell Trace Yellow Cell Proliferation Kit (ThermoFisher, #C34567) or Cell Trace CFSE Cell Proliferation Kit (ThermoFisher, #C34554) to label the ventricular zone of Pax6 and Emx2 mutants and their control littermates at E12.5. FlashTag labeling was conducted by injecting 0.3 µl of 10 mM of a carboxyfluorescein succinimidyl ester (CellTrace Yellow or Cell Trace CFSE, ThermoFisher) bilaterally into the fourth ventricle of E12.5 embryos (Telley et al., 2016). Gentle manual pressure was then applied to the exterior of the embryonic head to promote even mixing of the dyes. After 20 minutes, the cortices were then dissected and papain-treated (Papain dissociation system, Worthington Biochemical Corporation) for 15-30 minutes at 37°C with rotation. After inhibiting the papain, dissociated cortical cells were resuspended in 4% FBS/1x-PBS and the singlet FTag-positive population was sorted using the BD FACS Aria III Cell Sorter at Helen Diller Cancer Building (UCSF). Approximately 200,000 cells were used for native ChIP as described in (Sandberg et al., 2016). Wild-type cells and mutant littermates were always injected, sorted and processed side by side using the same number of nuclei. Basically, nuclei were extracted from the sorted cells and digested for 8 mins with micrococcal nuclease (MNase, Sigma). Mono and di-nucleosomes were combined and used for ChIP of two epigenomic marks: H3 acetylated lysine 27 (H3K27ac, Abcam, ab472) and H3 trimethyl lysine-27 (H3K27me3, Active motif, 39157). After immunoprecipatation, DNA and libraries were prepared as for TF ChIP as described above.

#### ATAC-seq

ATAC-seq was performed on around 80,000 sorted nuclei. Basically, we fluorescently labeled the VZ of wild-type and mutant littermates using the FlashTag procedure as indicated above. After making nuclei, the pellet was resuspended in 25uL of Tagment DNA buffer and 2uL of enzyme (Tagment DNA Enzyme, Nextera DNA Library Prep Kit, 15028211, Illumina). Tagmentation was performed at 37°C for 30 mins without shaking. Samples were then purified using MinElute columns (Qiagen), PCR-amplified for 8-10 cycles using the NEB Next High Fidelity 2x PCR Master Mix (NEB, 72 °C 5 min, 98 °C 30 s, (98 °C 10 s, 63 °C 30 s, 72 °C 60 s) per cycle, held at 72 °C). The generated amplified libraries were purified on Ampure XP Bead (Beckmann Coulter) and bioanalyzed. Sequencing was carried out on a HiSeq4000 (50 bp PE, Illumina).

#### PLAC-seq

PLAC-seq libraries for E12.5 cortex were prepared similar to the previously published protocol (Fang et al., 2016). 3 to 7 million cells were used for each library. If the cells appeared aggregated, they were dissociated with gentle MACS dissociator or dounce homogenizer. Each PLAC-seq library was prepared using DpnII as the restriction enzyme and Dynabeads M-280 sheep anti-rabbit IgG (Invitrogen #11203D) mixed with 5ug of H3K4me3 (04-745, Millipore) for the chromatin immunoprecipitation step. Finally, libraries were prepared with the Illumina Tru-seq adaptors and the final libraries were sent for paired-end sequencing on the HiSeq X Ten (150 bp reads).

#### PLAC-seq data analysis using the MAPS pipeline

We applied the MAPS pipeline (Juric et al., 2019) to detect statistically significant H3K4me3-centric long-range chromatin interactions from PLAC-seq data. We only analyzed intra-chromosomal interactions for autosomal chromosomes, and identified chromatin interactions at 10Kb bin resolution between 20Kb and 1Mb. We first mapped the raw paired-end reads (i.e. fastq file) to the mm10 reference genome using bwa mem. Next, we applied filtering steps to remove PCR duplicates, low quality reads (MAPQ <= 30) and chimeric reads (Juric et al., 2019). We then split the remaining mapped reads into short-range (distance between pair ends within 1Kb) and long-range reads (distance between pair ends between 1Kb and 1Mb). We used the short-range and long-range reads to measure protein immunoprecipitation (IP) efficiency and detect long-range chromatin interactions, respectively. In addition, we collected ChIP-seq peaks called by MACS2 from cortical cells. Among all 10Kb bin pairs, we only examined those 10Kb bin pairs in which at least one 10Kb bin overlapped with MACS2 ChIP-seq peaks, since these 10Kb bin pairs are enriched by H3K4me3 antibody during the PLAC-seq experiment (Zhang et al., 2008). We fitted a positive Poisson regression model on all selected 10Kb bin pairs with more than one raw count, taking into consideration multiple bias factors including 1D genomic distance between two interacting bins, restriction enzyme cut site frequency, GC content, mappability score, and H3K4me3 antibody IP efficiency measured by the number of short-range reads in each bin. After modeling fitting, we obtained expected contact frequency, p-value and false discovery rate (FDR) for each 10Kb bin pairs. Finally, we defined a 10Kb bin pair as a statistically significant long-range chromatin interaction if the raw contact frequency >= 12, normalized contact frequency (defined as observed contact frequency/expected contact frequency) >= 2, and FDR < 0.01. We further merged adjacent significant chromatin interactions together, and defined those isolated significant chromatin interactions as singletons. We applied a more stringent FDR threshold (FDR < 0.0001) among those singletons to reduce potential false positives.

#### Enhancer-Gene maps (definitions and annotation strategy)

Associations were previously defined by correlation between the H3K27ac profile of putative enhancers and the total-RNA profile of annotated genes across embryonic development. Two different maps were used, based on a dataset of 29 (Osterwalder et al., 2018) or 66 (Gorkin et al., 2020) samples representing time-courses considering up to 12 tissues and 7 time points. Given a list of genomic regions, a custom C++ script was used to annotate each one of these regions to the overlapping putative enhancers in these two maps (if any) and to associate them to the computationally inferred target gene.

#### Validated enhancers from the VISTA Enhancer Browser (derivation and annotation strategy)

Human and murine validated elements available on August 25, 2017 on the VISTA enhancer browser (http://enhancer.lbl.gov) (Visel et al., 2007). Human regions (hg19) were lifted to the mouse genome (mm10) using liftOver (Speir et al., 2016). This was run with default parameters except for *minmatch* that was set to 0.1 for mouse to human conversions and to 0.95 for mapping between mm9 and mm10. After that, the same script and strategy outlined in the previous paragraph were used to annotated a given list of genomic intervals to the regions in VISTA.

#### PLAC-seq data annotation

Using the same script and strategy outlined in the previous paragraphs, each end of the interaction was annotated to any overlapping: (1) VISTA element; (2) the putative target gene, as inferred separately from the two maps described in the previous paragraph; (3) TF-binding events, as inferred by ChIP-seq; in case of multiple overlapping peaks, the region was assigned the peaks with the highest enrichment score; (4) the closest gene on the linear genome, using the TSS of RefSeq genes as landmark. Coordinates of RefSeq genes were downloaded from the UCSC genome browser (Speir et al., 2016) on May 30, 2015. Merging of the resulting annotations was performed using the statistical computing environment R v3 (http://www.r-project.org).

#### Defining the interactome for loci in the Cortical Regionalization TF Network

We defined genomic loci based on the farthest PLAC-seq interaction between promoters and pREs for each loci. In cases in which this chromatin binding domain was restricted to only one side (upstream or downstream) of the gene body (i.e. *Bcl11a* or *Dmrt3*, Figure S5), we added a 100kb buffer on the other side of the TSS or 5’UTR.

#### Transgenic enhancer assays

Transgenic assays were performed according to published methods (Kothary et al., 1989; Pennacchio et al., 2006) and the VISTA enhancer browser can be consulted here: https://enhancer.lbl.gov. A summary of the methodology can be found in Lindtner et al, 2019.

