## Supplemental Figure 1 for "Transcriptional Network Orchestrating Regional Patterning of Cortical Progenitors"

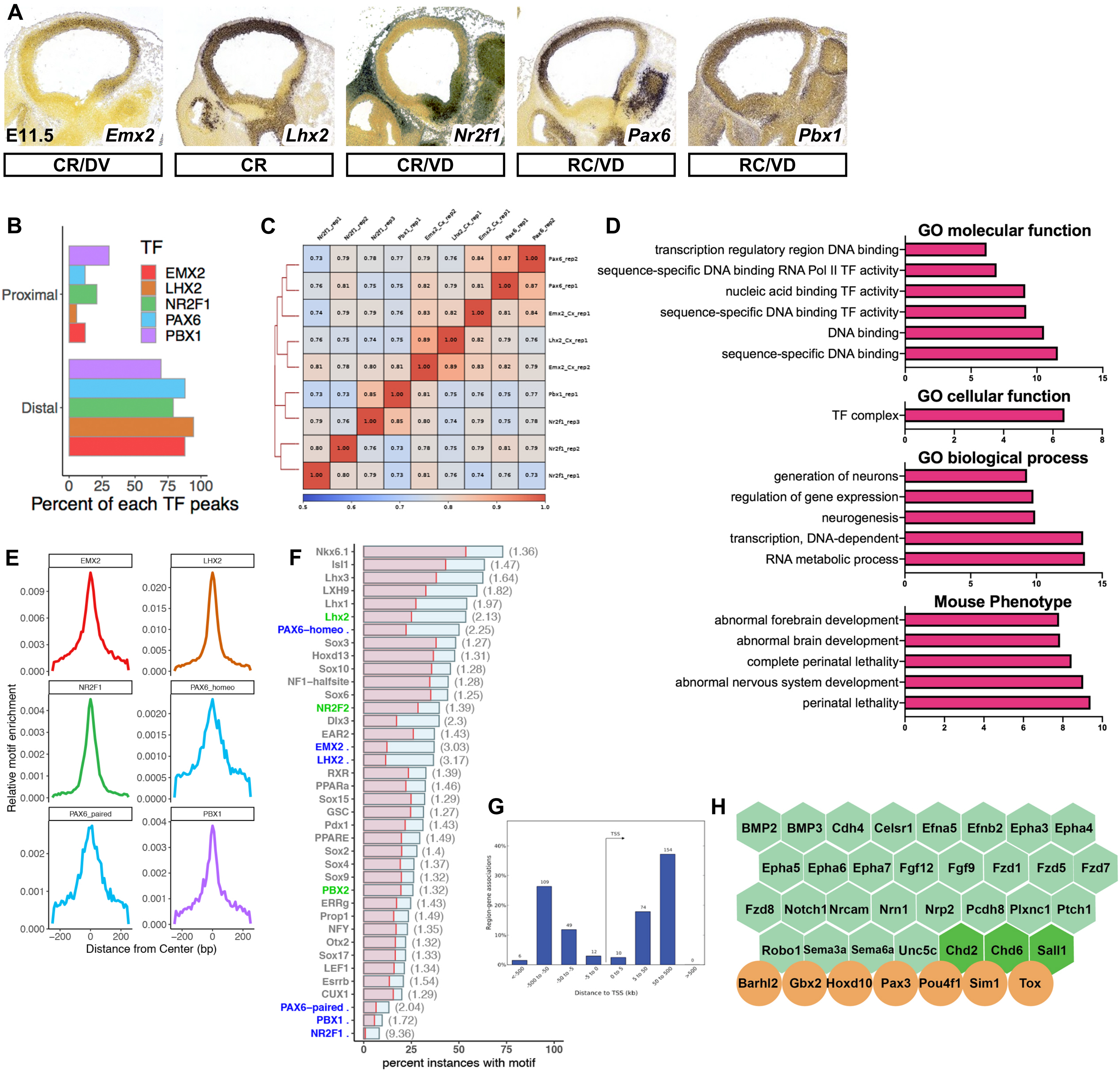


I

| | | | | | | | | | | | | | | | | | | | | | |
|---|---|---|---|---|---|---|---|---|---|---|---|---|---|---|---|---|---|---|---|---|---|
| | PBX1 | | | | | PBX1 | PBX1 | | | | | | PBX1 | PBX1 | | | | PBX1 | PBX1 | PBX1 | | PBX1 |
| | | LHX2 | | | | | | LHX2 | LHX2 | | | | LHX2 | | LHX2 | LHX2 | | LHX2 | LHX2 | | LHX2 | LHX2 |
| | | | PAX6 | | | | | | | PAX6 | PAX6 | | | | PAX6 | | PAX6 | PAX6 | | PAX6 | PAX6 | PAX6 |
| | | | | EMX2 | | EMX2 | | EMX2 | | EMX2 | | EMX2 | EMX2 | EMX2 | EMX2 | EMX2 | EMX2 | EMX2 | EMX2 | EMX2 | EMX2 | EMX2 |
| | | | | | NR2F1 | | NR2F1 | | NR2F1 | | NR2F1 | NR2F1 | | NR2F1 | | NR2F1 | NR2F1 | | NR2F1 | NR2F1 | NR2F1 | NR2F1 |
| Motif percentage | | | | | | | | | | | | | | | | | | | | | | |
| Pbx1 motif | 46.62 | 3.18 | 3.622 | 5.647 | 3.626 | 33.62 | 47.1 | 3.251 | 6.4 | 2.577 | 5.537 | 5.271 | 22.15 | 32.77 | 3.071 | 4.678 | 4.575 | 16.67 | 19.7 | 27.08 | 8.211 | 21.87 |
| Hbox_Emx2_Lhx2 | 11.24 | 97.48 | 26.2 | 70.5 | 16.13 | 55.6 | 13.41 | 96.2 | 84 | 67.53 | 27.36 | 62.11 | 93.9 | 50.21 | 92.13 | 95.22 | 62.09 | 93.43 | 93.79 | 56.25 | 96.19 | 95.67 |
| Paired_Pax6 | 11.57 | 6.469 | 60.39 | 9.17 | 8.221 | 11.94 | 16.3 | 8.678 | 12 | 38.92 | 40.72 | 11.82 | 13.96 | 14.47 | 34.17 | 14.55 | 35.95 | 27.78 | 13.92 | 43.75 | 36.07 | 35.54 |
| Hbox_Pax6 | 2.274 | 3.838 | 45.43 | 5.669 | 3.746 | 5.39 | 5.072 | 5.234 | 3.2 | 25.26 | 27.36 | 4.274 | 7.063 | 13.62 | 22.46 | 7.692 | 20.92 | 19.7 | 6.424 | 30.21 | 21.99 | 21.87 |
| Nuclear_ NR2F1 | 0.134 | 0.329 | 0.768 | 0.131 | 41.05 | 0.146 | 7.609 | 0.11 | 4.8 | 0 | 10.42 | 6.125 | 0.321 | 2.553 | 0.192 | 3.846 | 3.922 | 0 | 3.64 | 2.083 | 3.812 | 2.506 |

| | | | |
|---|---|---|---|
| <10 | <33 | <66 | <100 |
