## Supplementary figures and images for "Transcriptional Network Orchestrating Regional Patterning of Cortical Progenitors"

### Supplemental Figure 2

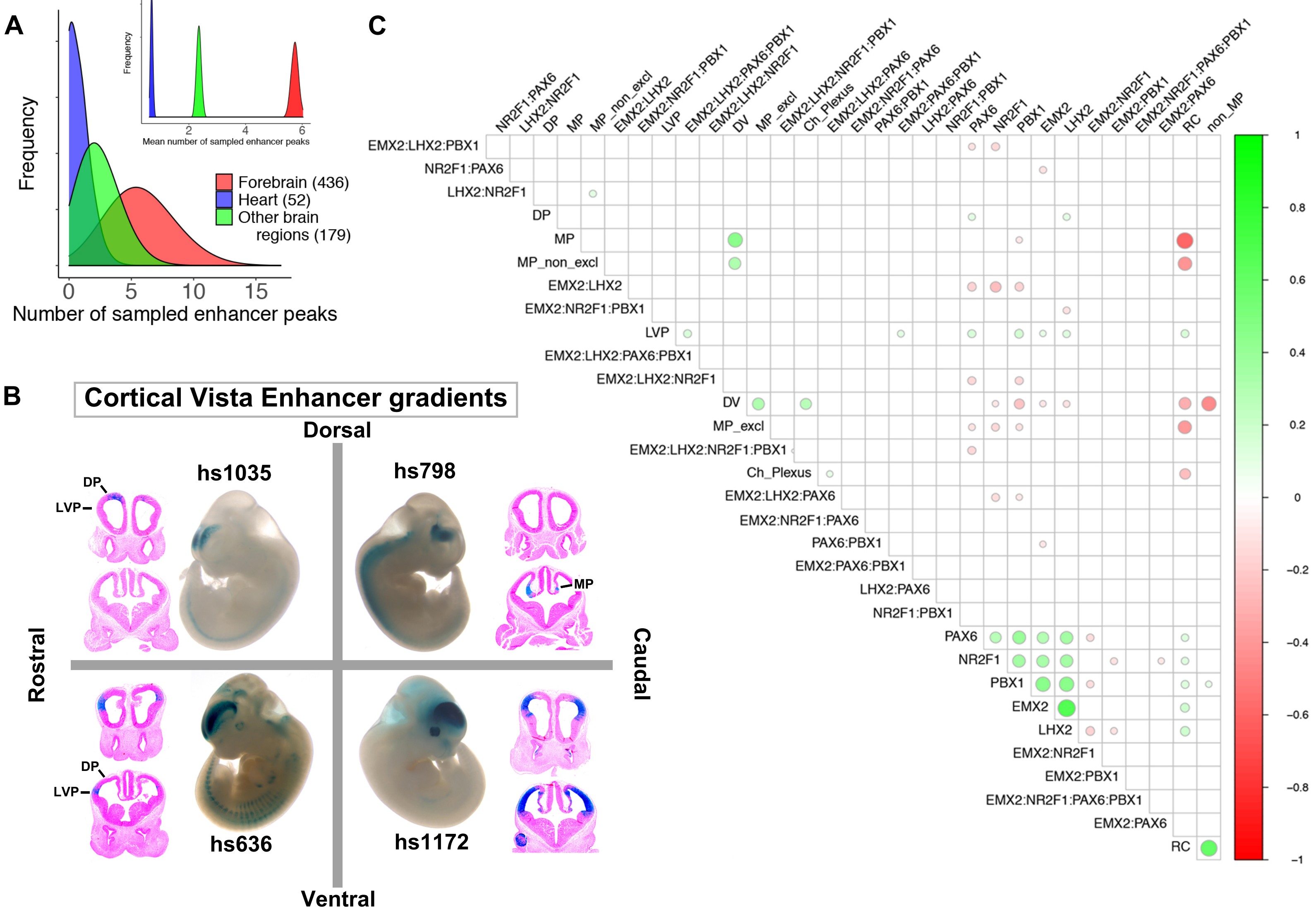

### Supplemental Figure 3

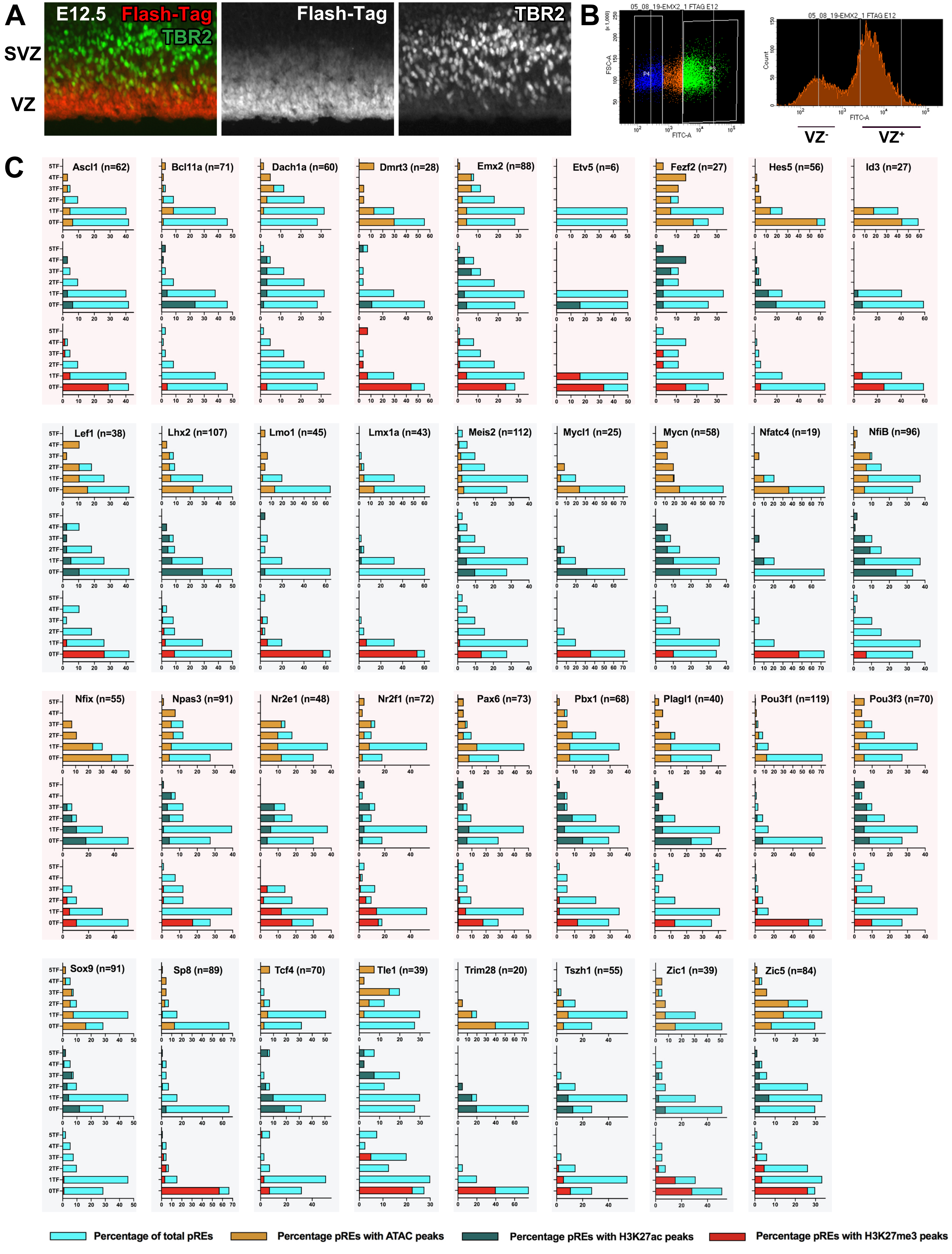

### Supplemental Figure 6

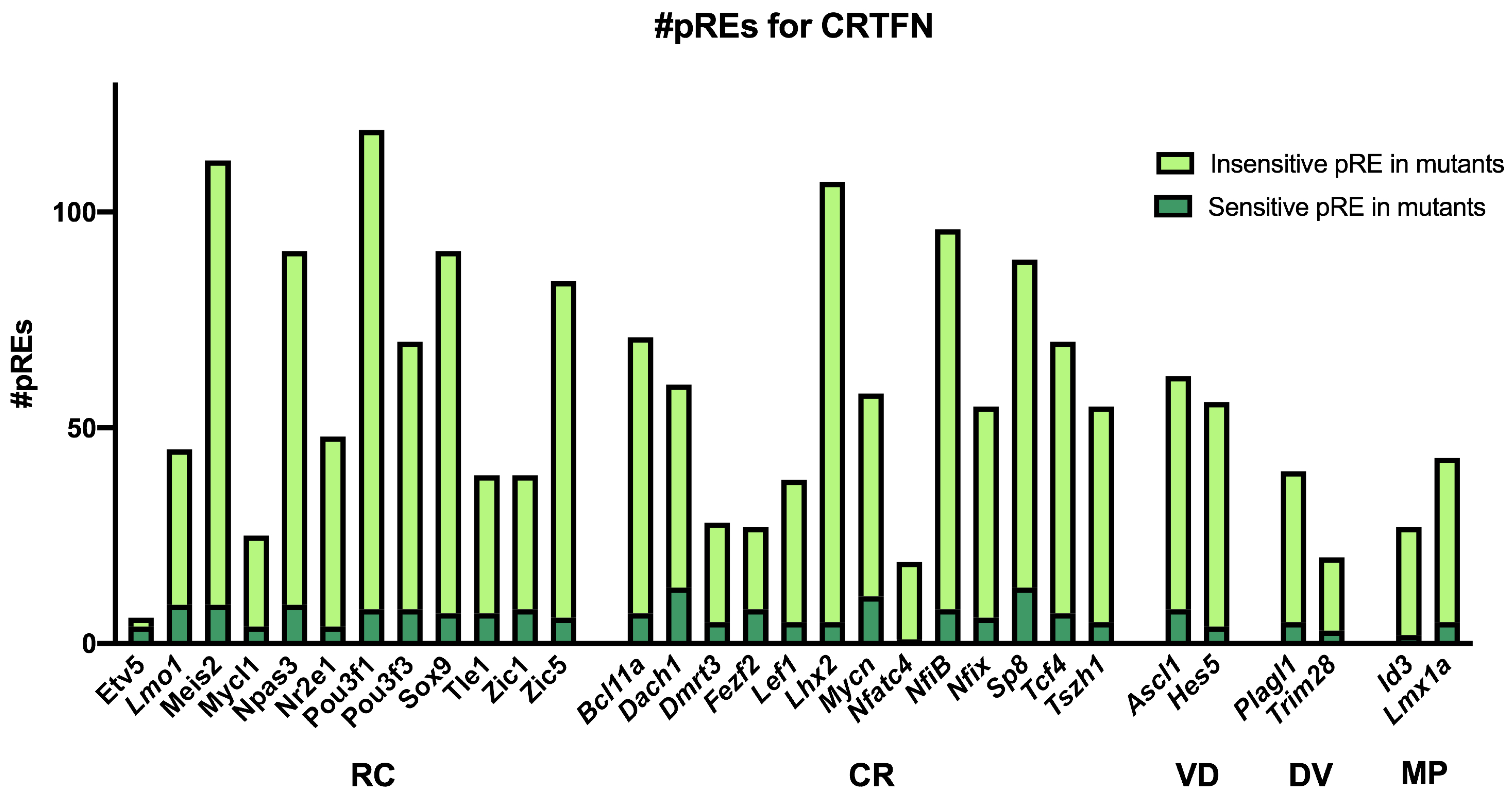
