## Supplemental Figure 4 for "Transcriptional Network Orchestrating Regional Patterning of Cortical Progenitors"

**A**

| | | # pREs | TF ChIP-Seq | | | | | Ratio H3K27ac / H3K27me3 | VZ Epigenomic Marks | | |
|---|---|---|---|---|---|---|---|---|---|---|---|
| | | | Emx2 | Lhx2 | Nr2f1 | Pax6 | Pbx1 | | H3K27ac | H3K27me3 | ATAC |
| **TFs with Rostral to Caudal Gradient** | | | | | | | | | | | |
| *Lmo1* | ★ | 45 | 20.0 | 17.8 | 13.3 | 15.6 | 4.4 | 0.1 | 8.9 | 68.9 | 31.1 |
| *Pou3f1* | | 119 | 14.3 | 10.9 | 13.4 | 5.9 | 7.6 | 0.2 | 10.9 | 63.9 | 23.5 |
| *Zic1* | O | 39 | 23.1 | 20.5 | 15.4 | 20.5 | 2.6 | 0.3 | 12.8 | 46.2 | 38.5 |
| *Etv5* | | 6 | 16.7 | 16.7 | 0.0 | 16.7 | 0.0 | 0.3 | 16.7 | 50.0 | 0.0 |
| *Tle1* | O | 39 | 42.5 | 42.5 | 27.5 | 35.0 | 15.0 | 0.5 | 12.5 | 27.5 | 32.5 |
| *Zic5* | O | 84 | 46.4 | 21.4 | 26.2 | 17.9 | 11.9 | 0.5 | 17.9 | 35.7 | 48.8 |
| *Nr2e1* | O | 48 | 34.0 | 16.0 | 44.0 | 20.0 | 2.0 | 0.7 | 26.0 | 36.0 | 44.0 |
| *Npas3* | | 91 | 39.6 | 24.2 | 35.2 | 19.8 | 17.6 | 1.0 | 19.8 | 19.8 | 30.8 |
| *Pax6* | ★ | 73 | 35.6 | 23.3 | 26.0 | 28.8 | 9.6 | 1.1 | 26.0 | 24.7 | 39.7 |
| *Mycl1* | ★ | 25 | 8.0 | 0.0 | 24.0 | 0.0 | 4.0 | 1.1 | 40.0 | 36.0 | 36.0 |
| *Meis2* | | 112 | 37.5 | 26.8 | 33.0 | 23.2 | 13.4 | 1.7 | 27.7 | 16.1 | 22.3 |
| *Pbx1* | | 68 | 44.1 | 30.9 | 26.5 | 13.2 | 13.2 | 2.6 | 38.2 | 14.7 | 35.3 |
| *Pou3f3* | | 70 | 37.1 | 25.7 | 41.4 | 27.1 | 14.3 | 2.9 | 37.1 | 12.9 | 31.4 |
| *Sox9* | | 91 | 30.8 | 22.0 | 37.4 | 18.7 | 13.2 | 13.5 | 29.7 | 2.2 | 40.7 |
| **TFs with Caudal to Rostral Gradient** | | | | | | | | | | | |
| *Sp8* | ✚ | 89 | 22.6 | 14.3 | 14.3 | 10.7 | 7.1 | 0.1 | 7.1 | 69.0 | 28.6 |
| *Dmrt3* | O | 28 | 18.5 | 14.8 | 22.2 | 18.5 | 11.1 | 0.2 | 14.8 | 63.0 | 51.9 |
| *Nfatc4* | O | 19 | 5.3 | 5.3 | 26.3 | 0.0 | 0.0 | 0.3 | 15.8 | 47.4 | 52.6 |
| *Emx2* | O | 88 | 47.7 | 36.4 | 25.0 | 12.5 | 9.1 | 0.6 | 18.2 | 30.7 | 26.1 |
| *Nr2f1* | O | 72 | 36.1 | 26.4 | 43.1 | 22.2 | 13.9 | 0.6 | 22.2 | 37.5 | 31.9 |
| *Lef1* | ✚ | 38 | 28.9 | 26.3 | 23.7 | 21.1 | 13.2 | 0.8 | 23.7 | 28.9 | 50.0 |
| *Tszh1* | | 55 | 25.5 | 20.0 | 25.5 | 12.7 | 10.9 | 1.3 | 23.6 | 18.2 | 21.8 |
| *Fezf2* | ✚ | 27 | 48.1 | 44.4 | 33.3 | 22.2 | 18.5 | 1.7 | 37.0 | 22.2 | 63.0 |
| *Nfix* | ✚ | 55 | 18.2 | 7.3 | 40.0 | 7.3 | 1.8 | 2.0 | 40.0 | 20.0 | 80.0 |
| *Lhx2* | | 107 | 21.5 | 19.6 | 25.2 | 10.3 | 11.2 | 3.2 | 50.5 | 15.9 | 43.9 |
| *Tcf4* | ✚ | 70 | 34.8 | 20.3 | 26.1 | 20.3 | 8.7 | 3.4 | 39.1 | 11.6 | 18.8 |
| *Mycn* | | 58 | 32.8 | 25.9 | 39.7 | 10.3 | 8.6 | 4.2 | 43.1 | 10.3 | 48.3 |
| *Dach1* | | 60 | 43.3 | 45.0 | 18.3 | 20.0 | 11.7 | 4.5 | 15.0 | 3.3 | 18.3 |
| *NfiB* | ✚ | 96 | 44.8 | 25.0 | 19.8 | 16.7 | 8.3 | 6.7 | 49.0 | 7.3 | 34.4 |
| *Bcl11a* | ✚ | 71 | 25.4 | 14.1 | 22.5 | 15.5 | 5.6 | 7.7 | 32.4 | 4.2 | 16.9 |
| **TFs with Ventral-Dorsal Gradient** | | | | | | | | | | | |
| *Ascl1* | ★ | 62 | 29.0 | 11.3 | 24.2 | 21.0 | 1.6 | 0.3 | 12.9 | 37.1 | 19.4 |
| *Hes5* | ★ | 56 | 16.1 | 7.1 | 23.2 | 5.4 | 1.8 | 7.3 | 39.3 | 5.4 | 82.1 |
| **TFs with Dorsal-Ventral Gradient** | | | | | | | | | | | |
| *Trim28* | ✚ | 20 | 5.0 | 0.0 | 10.0 | 5.0 | 10.0 | 1.0 | 40.0 | 40.0 | 60.0 |
| *Plagl1* | ✚ | 40 | 28.2 | 28.2 | 23.1 | 20.5 | 7.7 | 3.4 | 43.6 | 12.8 | 41.0 |
| **TFs with Expression in MP** | | | | | | | | | | | |
| *Lmx1a* | | 43 | 18.6 | 14.0 | 9.3 | 2.3 | 4.7 | 0.1 | 4.7 | 60.5 | 20.9 |
| *Id3* | | 27 | 0.0 | 3.7 | 37.0 | 0.0 | 0.0 | 0.3 | 11.1 | 33.3 | 63.0 |

★ Ventral-Dorsal
✚ Dorsal-Ventral
O Medial Pallium

**B**

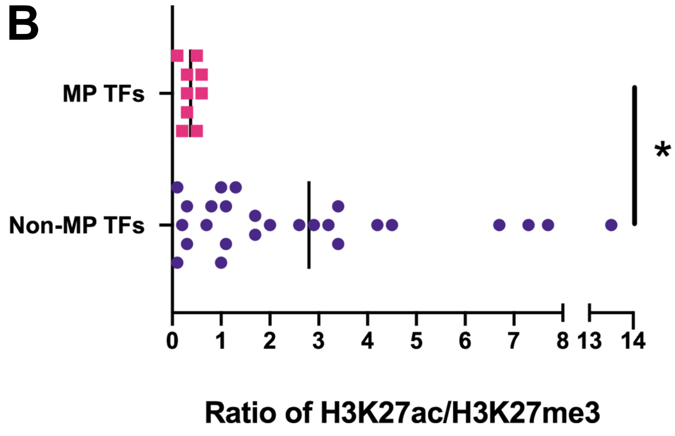


**C**

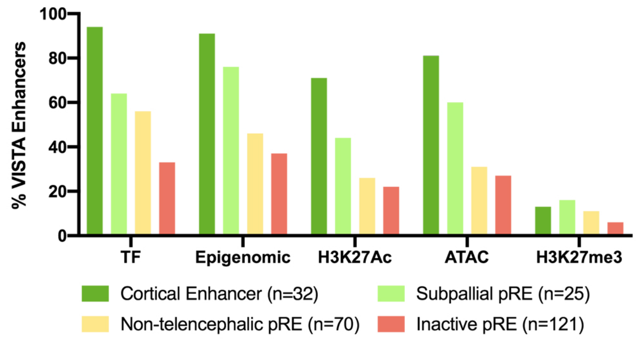
