## Supplemental Figure 5 for "Transcriptional Network Orchestrating Regional Patterning of Cortical Progenitors"

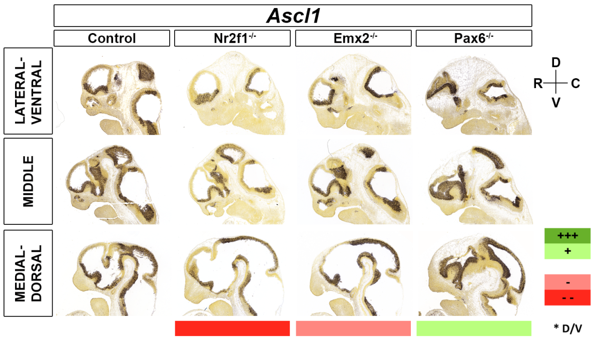


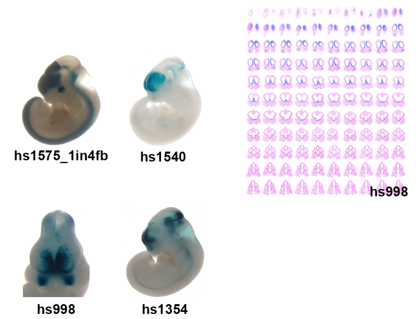


| | | Motifs | | | | | TF ChIP Seq | | | | | Differential epigenomic marks in WT vs. Mutant | | | | | | | | | Plac | Computation | Genomic features |
|---|---|---|---|---|---|---|---|---|---|---|---|---|---|---|---|---|---|---|---|---|---|---|---|
| | | | | | | | | | | | | Pax6 mutant | | | Emx2 mutant | | | Nr2f1 mutant | | | | | |
| | | Emx2 HB | Nr2f1 | Pax6 Hb | Pax6 P | Pbx1 | Emx2 | Lhx2 | Nr2f1 | Pax6 | Pbx1 | 27Ac | 27me3 | ATAC | 27Ac | 27me3 | ATAC | 27Ac | 27me3 | ATAC | | | |
| Ascl1 | chr10:86400000-87800000 | | | | | | | | | | | | | | | | | | | | | | |
| 1 | chr10:86434673-86435586 | | | | | | | | x | | | | | | | | | | | Small Gain | x | | |
| 2 | chr10:86487025-86488510 | | | | | | x | | | | | | | | | | Gain | | | Small Gain | x | | |
| 3 | chr10:86563808-86565878 | x | | x | x | | x | x | x | x | | Same | | Same | Loss | | Gain | Same | | Same | x | | |
| 4 | chr10:86691462-86694122 | | | | | | | | | | | | | | | Gain | | | | | x | | 5"UTR 1700113h08Rik |
| 5 | chr10:86906349-86912592 | x | | | | | x | | | | | Same | | Loss | Loss | | Same | Same | | Same | | x | hs1540 |
| 6 | chr10:87387234-87391021 | | | x | x | | x | | x | x | | | Small Loss | Same | | Small Loss | Same | | Same | Same | | | Igf1 Gene Body |
| 7 | chr10:87605288-87611277 | x | | | | | | | | x | | Loss | | Loss | Same | | Same | Same | | Same | x | | TSS Nup37 |
| 8 | chr10:87660733-87665800 | | | | | | | | | | | Loss | | Loss | Same | | Same | | | Same | x | | TSS Cdc53 |

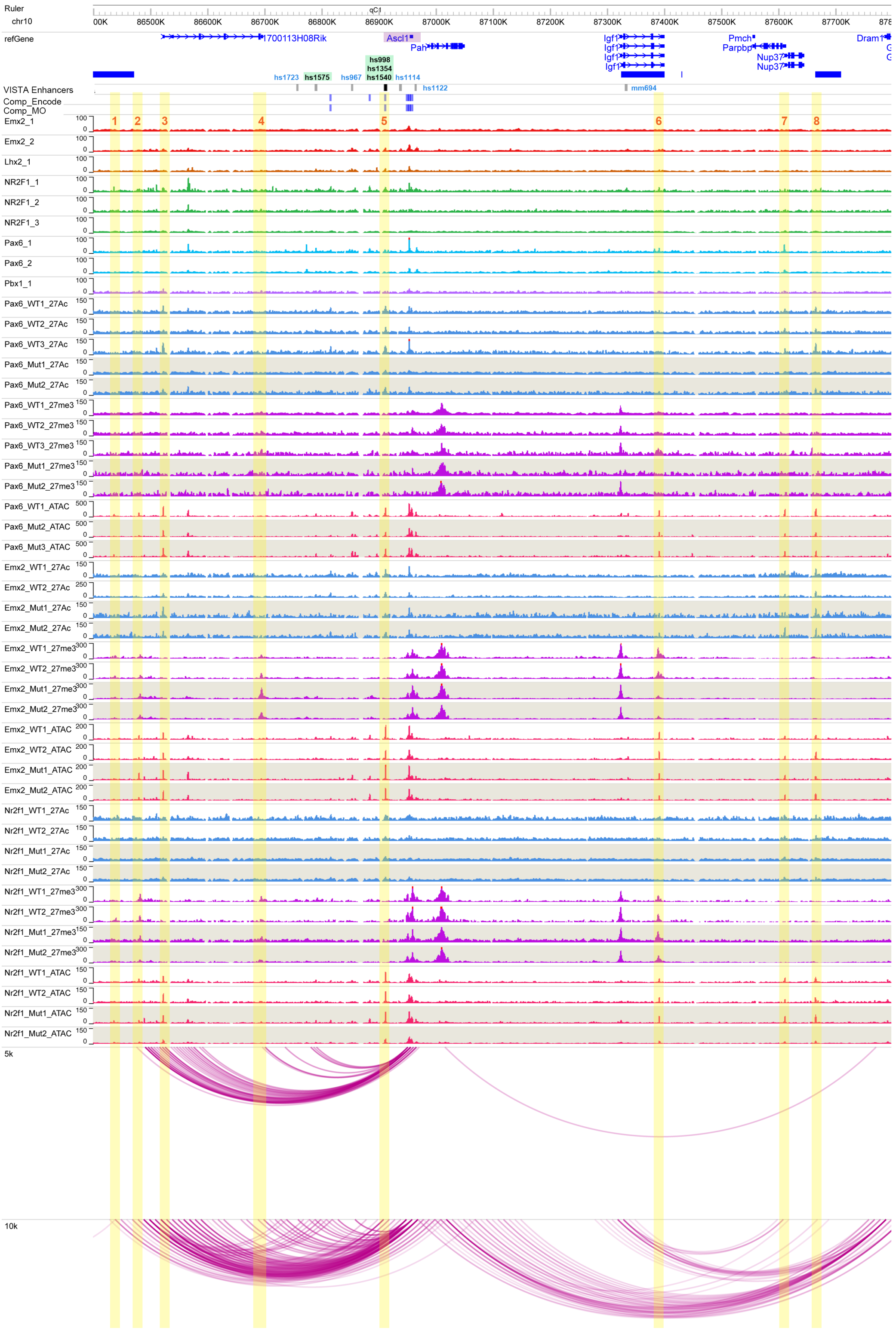

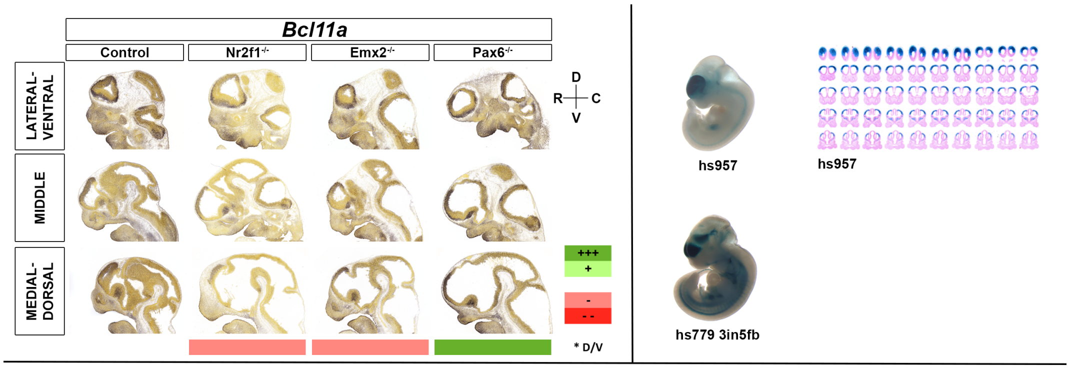


| | | Motifs | | | | | TF ChIP Seq | | | | | Differential epigenomic marks in WT vs. Mutant | | | | | | | | | Plac | Compu tation | Genomic features |
|---|---|---|---|---|---|---|---|---|---|---|---|---|---|---|---|---|---|---|---|---|---|---|---|
| | | | | | | | | | | | | Pax6 mutant | | | Emx2 mutant | | | Nr2f1 mutant | | | | | |
| | | Emx2 HB | Nr2f1 | Pax6 Hb | Pax6 P | Pbx1 | Emx2 | Lhx2 | Nr2f1 | Pax6 | Pbx1 | 27Ac | 27me3 | ATAC | 27Ac | 27me3 | ATAC | 27Ac | 27me3 | ATAC | | | |
| Bcl11a | chr11:23875000-25000000 | | | | | | | | | | | | | | | | | | | | | | |
| 1 | chr11:23949196-23950435 | | | | | | | | | | | Loss | | Same | Loss | | Same | Loss | | Same | | x | |
| 2 | chr11:23962200-23963523 | | | | | | | | | | | | Same | | | Gain | | | Same | | | | hs1176 |
| 3 | chr11:23973144-23974804 | | x | | | | | | x | | | | | | | Gain | | | | | | x | |
| 4 | chr11:23976082-23978715 | | | | | | | | x | | | Loss | | Same | Same | | Same | Loss | | Same | | x | TSS Bcl11a |
| 5 | chr11 23995402 23996537 | | | | | | x | x | x | x | x | Loss | | Same | Same | | Loss | Same | | Same | | x | hs957 |
| 6 | chr11:24270356-24271960 | | | | | | x | x | x | x | x | Loss | | Same | Loss | Gain | Same | Loss | | Same | x | | hs399 |
| 7 | chr11:24327290-24328274 | | | | | | x | | x | | | | | Same | | | Gain | | | Small Gain | x | | hs779 |

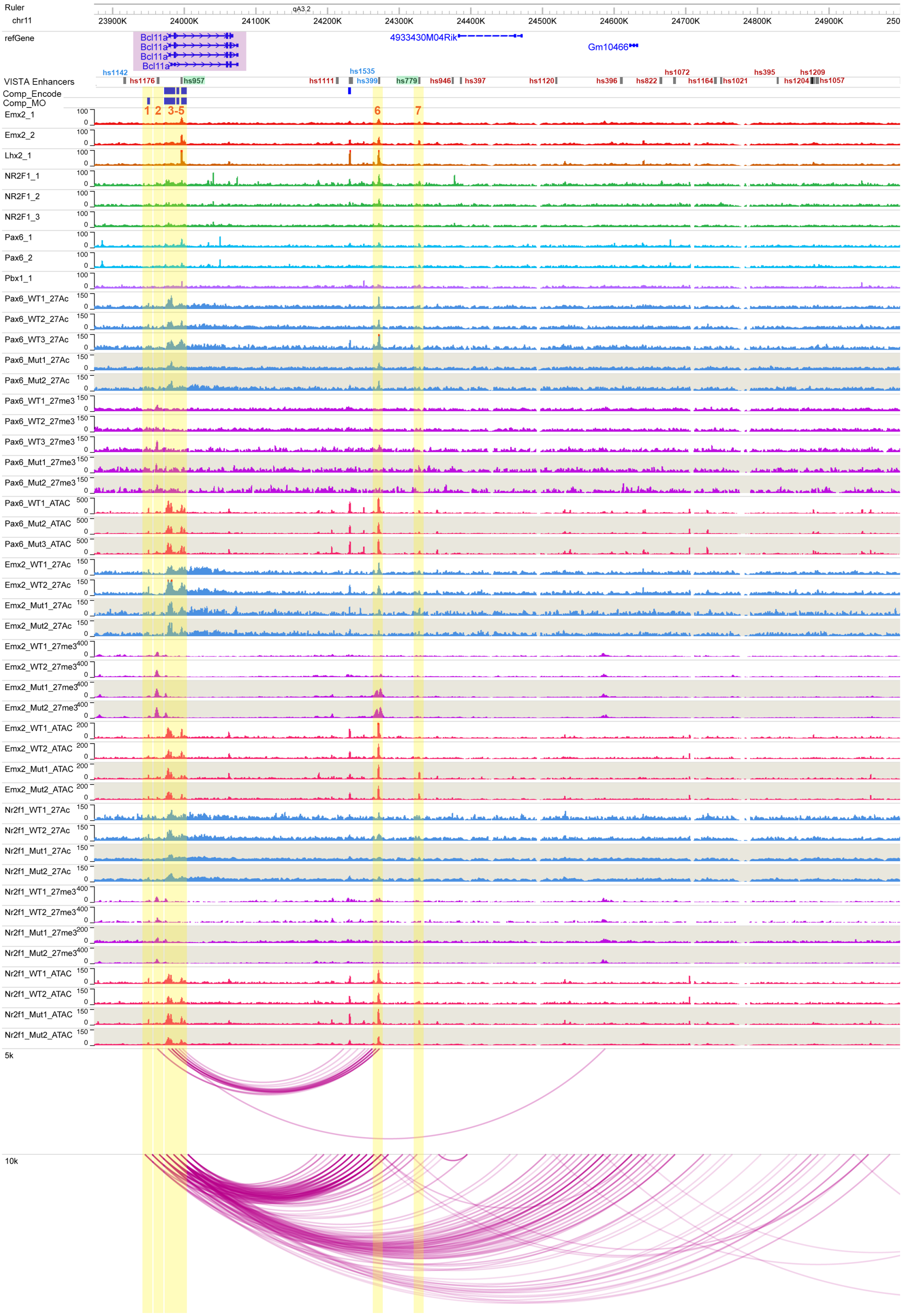

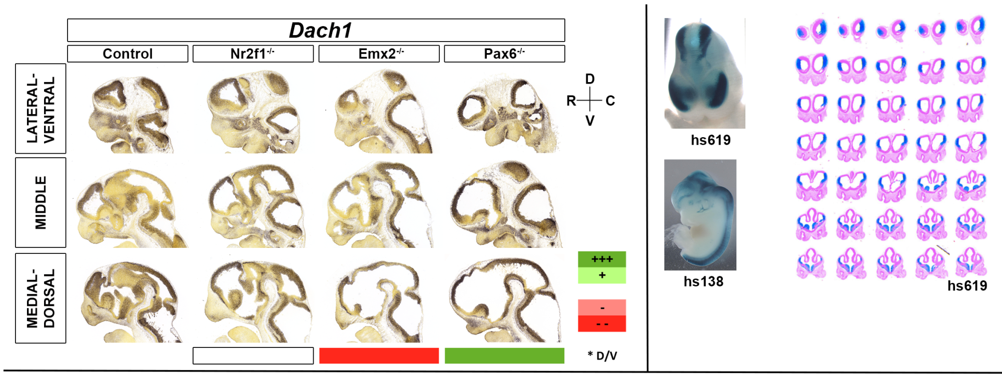


| Dach1a | chr14:97700000-99400000 | Motifs: Emx2 HB | Motifs: Nr2f1 | Motifs: Pax6 Hb | Motifs: Pax6 P | Motifs: Pbx1 | TF ChIP Seq: Emx2 | TF ChIP Seq: Lhx2 | TF ChIP Seq: Nr2f1 | TF ChIP Seq: Pax6 | TF ChIP Seq: Pbx1 | Differential epigenomic marks in WT vs. Mutant — Pax6 mutant: 27Ac | Pax6 mutant: 27me3 | Pax6 mutant: ATAC | Emx2 mutant: 27Ac | Emx2 mutant: 27me3 | Emx2 mutant: ATAC | Nr2f1 mutant: 27Ac | Nr2f1 mutant: 27me3 | Nr2f1 mutant: ATAC | Plac | Computation | Genomic features |
|---|---|---|---|---|---|---|---|---|---|---|---|---|---|---|---|---|---|---|---|---|---|---|---|
| 1 | chr14:97794887-97795910 | | | | | | x | x | x | | | | | Small Gain | | | Small Gain | | | Small Gain | | | |
| 2 | chr14:97936954-97937320 | x | | | | | x | x | | | | | | Small Gain | | | | | | | x | | hs129 |
| 3 | chr14:97984506-97988219 | | | | | | x | x | x | | | | | Same | | Gain | Same | | | Same | x | | |
| 4 | chr14:98213227-98214488 | x | | x | x | | x | x | | x | | Same? | | Same | Loss | | Loss | Same? | | Same | x | | |
| 5 | chr14:98471567-98473364 | x | | | | x | x | x | x | | x | Small Loss | | Same | Loss | Gain | Same | Same | | Same | x | | |
| 6 | chr14:98507543-98508878 | x | | | | | x | x | | x | x | Same | | Same | Loss | | Same? | Same | | Same | x | | |
| 7 | chr14:98522021-98522688 | x | | | | x | x | x | x | x | x | | | Loss | | | Loss | | | Same | | | |
| 8 | chr14:98533233-98535459 | x | | x | x | | x | x | x | x | | | | Loss | | | Loss | | | Same | | | |
| 9 | chr14:98564922-98570142 | | | | | | | | x | | | Loss | | Same | Loss | Gain | Same | Same | | Same | | | Dach1a TSS |
| 10 | chr14:98571770-98573377 | x | | | | | x | x | | | x | | | Same | | Gain | Loss | | | Same | | x | |
| 11 | chr14:98787230-98787662 | x | | | | | x | x | | | x | | | Same | | | Loss | | | Same | | | |
| 12 | chr14:98935571-98938292 | | | | | | x | | x | | | | | Gain | | | | | | | | | hs131 |
| 13 | chr14:99256151-99264870 | | | | | | | | | | | | | | | Small Gain | | | | | x | | |

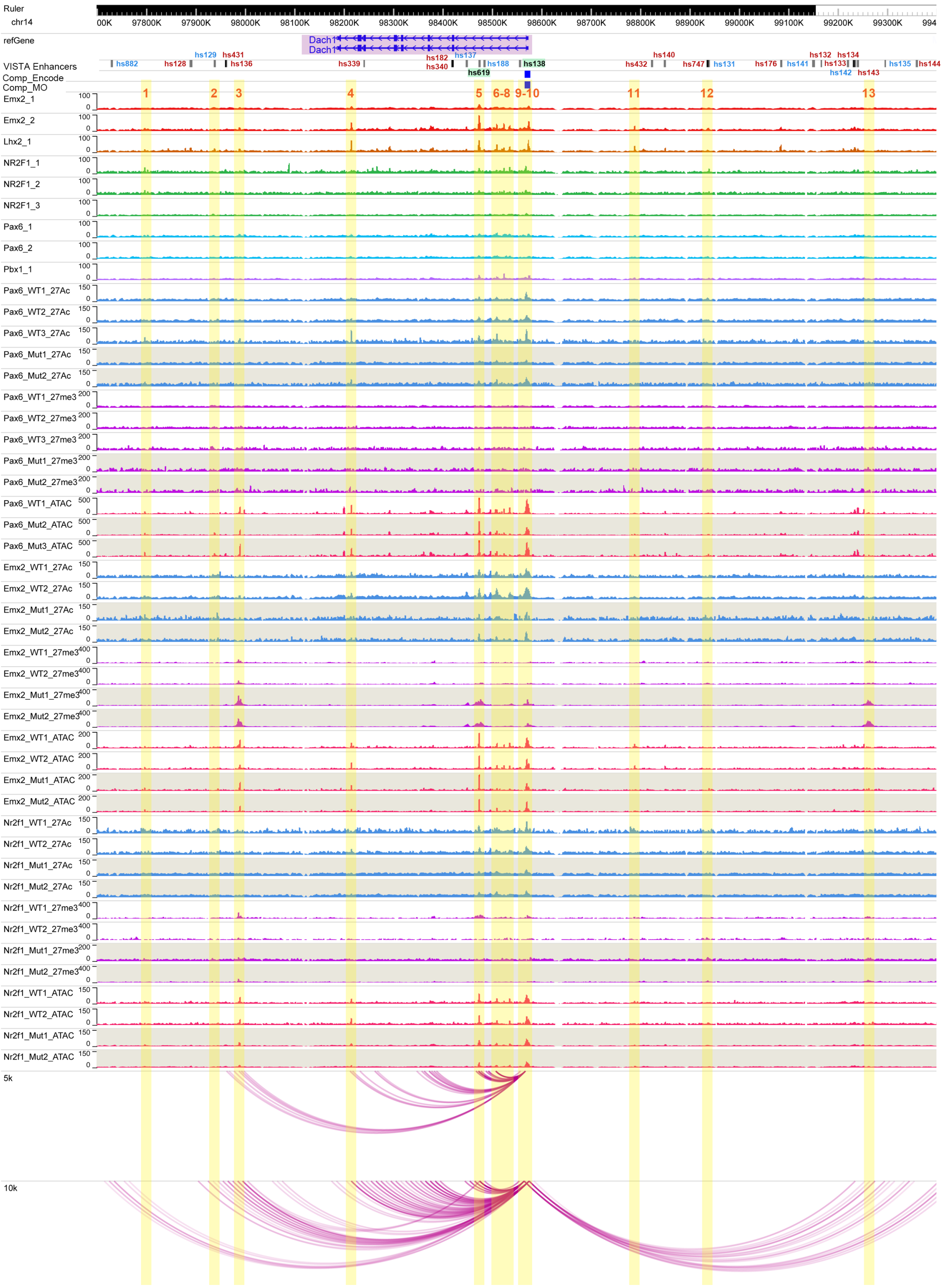

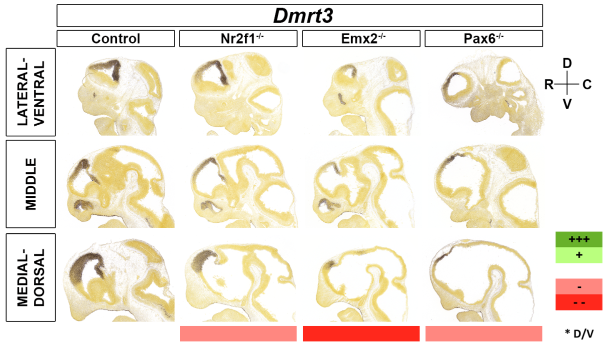


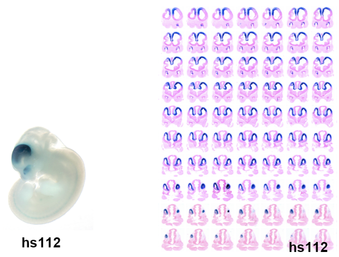


| | | Motifs | | | | | TF ChIP Seq | | | | | Differential epigenomic marks in WT vs. Mutant | | | | | | | | | Plac | Computation | Genomic features |
|---|---|---|---|---|---|---|---|---|---|---|---|---|---|---|---|---|---|---|---|---|---|---|---|
| | | | | | | | | | | | | Pax6 mutant | | | Emx2 mutant | | | Nr2f1 mutant | | | | | |
| | | Emx2 HB | Nr2f1 | Pax6 Hb | Pax6 P | Pbx1 | Emx2 | Lhx2 | Nr2f1 | Pax6 | Pbx1 | 27Ac | 27me3 | ATAC | 27Ac | 27me3 | ATAC | 27Ac | 27me3 | ATAC | | | |
| Dmrt3 | chr19:25014000-25800000 | | | | | | | | | | | | | | | | | | | | | | |
| Ctrl1 | chr19:25579959-25581177 | | | | | | | | | | | | Same | Same | | Same | Same | | Same | Same | | | Dmrt1 TSS |
| 1 | chr19:25673010-25673662 | x | | | | | x | x | | | x | | | Loss | | Gain | Loss | | | Same | | | |
| 2 | chr19:25676337-25677056 | x | | | | | x | x | x | x | x | | Gain | Loss | | Gain | Loss | | Same | Same | | | |
| 3 | chr19:25682256-25683612 | x | | x | x | x | x | x | x | x | x | Loss | Gain | Loss | Loss | Gain | Same | Loss | Same | Same | | x | hs112 |
| 4 | chr19:25684275-25686162 | | | | | | | | | | | Loss | Gain | Loss | Loss | Gain | Same | Loss | Same | Same | | x | Dmrt3 TSS |
| 5 | chr19:25698077-25700494 | | | | | | | | | | | | Gain | | | Gain | | | Same | | | | Dmrt3 5'UTR |
| Ctrl2 | chr19:25746293-25747747 | | x | | | | | | | | | | Same | Same | | Same | Same | | Loss | Same | | | Dmrt2 TSS |

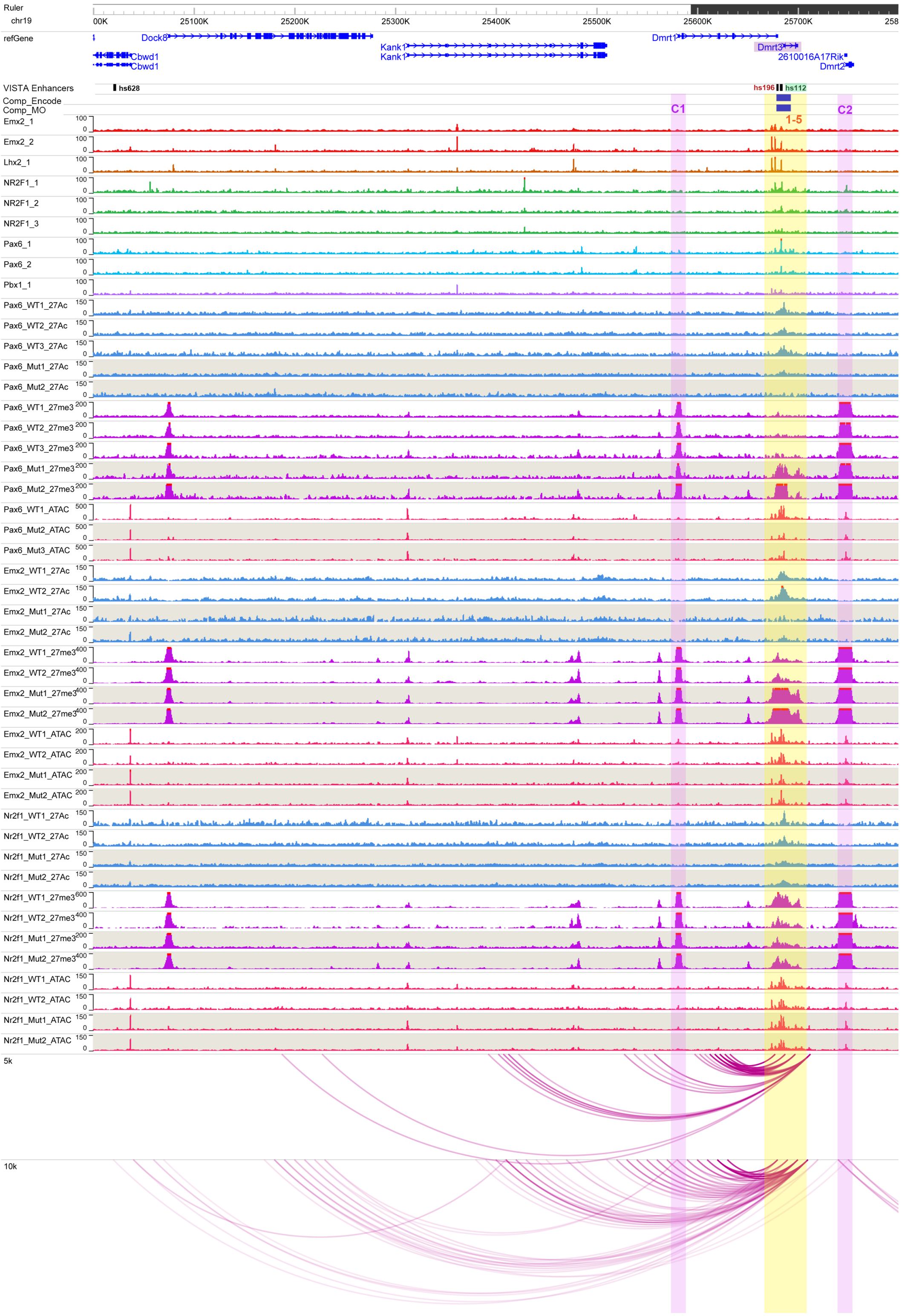

### *Dmrt5*

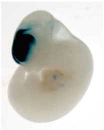

hs200

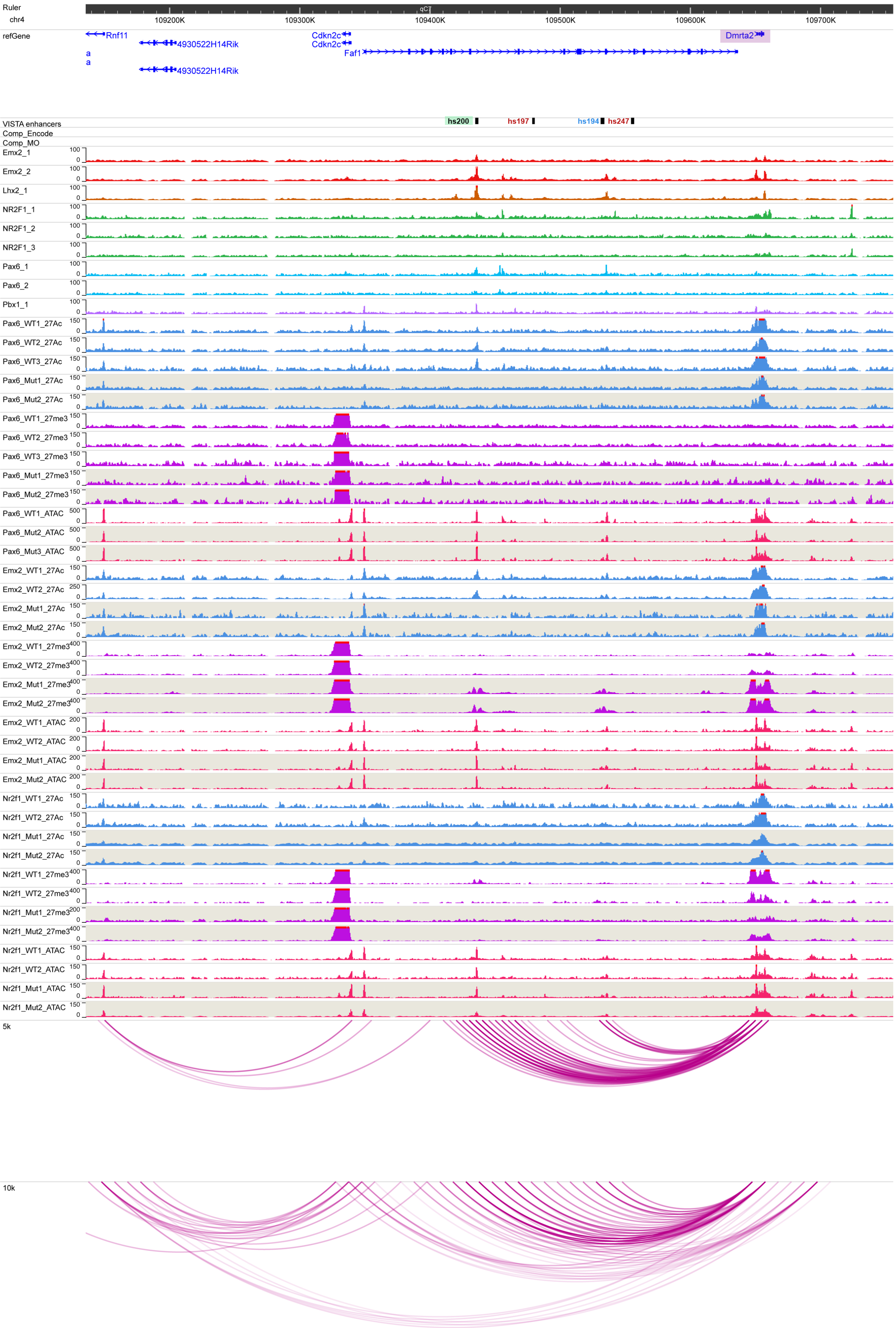

Ruler
chr4
109200K
109300K
109400K
109500K
109600K
109700K
refGene
Rnf11
4930522H14Rik
Cdkn2c
Faf1
Dmrta2
VISTA enhancers
hs200
hs197
hs194
hs247
Comp_Encode
Comp_MO
Emx2_1
Emx2_2
Lhx2_1
NR2F1_1
NR2F1_2
NR2F1_3
Pax6_1
Pax6_2
Pbx1_1
Pax6_WT1_27Ac
Pax6_WT2_27Ac
Pax6_WT3_27Ac
Pax6_Mut1_27Ac
Pax6_Mut2_27Ac
Pax6_WT1_27me3
Pax6_WT2_27me3
Pax6_WT3_27me3
Pax6_Mut1_27me3
Pax6_Mut2_27me3
Pax6_WT1_ATAC
Pax6_Mut2_ATAC
Pax6_Mut3_ATAC
Emx2_WT1_27Ac
Emx2_WT2_27Ac
Emx2_Mut1_27Ac
Emx2_Mut2_27Ac
Emx2_WT1_27me3
Emx2_WT2_27me3
Emx2_Mut1_27me3
Emx2_Mut2_27me3
Emx2_WT1_ATAC
Emx2_WT2_ATAC
Emx2_Mut1_ATAC
Emx2_Mut2_ATAC
Nr2f1_WT1_27Ac
Nr2f1_WT2_27Ac
Nr2f1_Mut1_27Ac
Nr2f1_Mut2_27Ac
Nr2f1_WT1_27me3
Nr2f1_WT2_27me3
Nr2f1_Mut1_27me3
Nr2f1_Mut2_27me3
Nr2f1_WT1_ATAC
Nr2f1_WT2_ATAC
Nr2f1_Mut1_ATAC
Nr2f1_Mut2_ATAC
5k
10k

#### *Emx2*

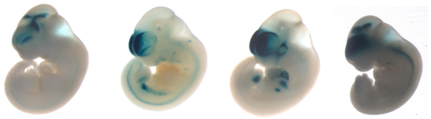

hs1032 hs1236 mm449 hs672

| | | TF ChIP Seq | | | | | Differential epigenomic marks in WT vs. Mutant | | | | | | | | | Plac-Seq | Computation | Genomic features |
|---|---|---|---|---|---|---|---|---|---|---|---|---|---|---|---|---|---|---|
| | | | | | | | Pax6 mutant | | | Emx2 mutant | | | Nr2f1 mutant | | | | | |
| | | Emx2 | Lhx2 | Nr2f1 | Pax6 | Pbx1 | 27Ac | 27me3 | ATAC | 27Ac | 27me3 | ATAC | 27Ac | 27me3 | ATAC | | | |
| **Emx2** | **chr19:59400000-60526000** | | | | | | | | | | | | | | | | | |
| **1** | chr19:59525206-59526676 | | | | | | Loss | | Same | Loss | | Same | Same | | Same | | x | |
| **2** | chr19:59531268-59535857 | x | x | x | | x | Loss | | Same | Loss | | Same | Same | | Same | | x | Emx2 TSS |
| **3** | chr19:59536094-59538976 | x | | x | | | Loss | | Same | Loss | | Loss | Same | | Same | | x | Emx2 Gene Body |
| **4** | chr19:59539919-59541154 | x | x | x | x | | Loss | | Same | Loss | | Same | Same | | Same | | x | hs1032 |
| **5** | chr19:59541331-59542154 | x | x | x | | | Loss | | Same | Loss | | Same | Same | | Same | | x | hs935 |
| **6** | chr19:59543685-59546274 | x | x | x | | | Loss | | Same | Loss | Gain | Same | Same | | Same | | x | hs1221 |
| **7** | chr19:59773144-59774850 | x | x | x | x | x | | | Loss | | | Same | | | Same | x | | |
| **8** | chr19:59963467-59965820 | x | x | | | | | | Same | | | Loss | | | Same | x | | |
| **9** | chr19:59986713-59988654 | x | x | x | | x | Loss | | Same | Loss | | Loss | Same | | Same | x | | Rab11fip2 |
| **10** | chr19:60091231-60093643 | x | x | x | | x | Loss | | Same | Loss | | Same | Same | | Same | x | | mm449 |
| **11** | chr19:60275728-60287674 | x | x | | | | | Same | Same | | Loss | Same | | Same | Same | x | | |
| **12** | chr19:60301681-60303387 | | | | | | Same | | Same | Same | | Same | Loss | | Same | x | | D19ertd737e TSS |

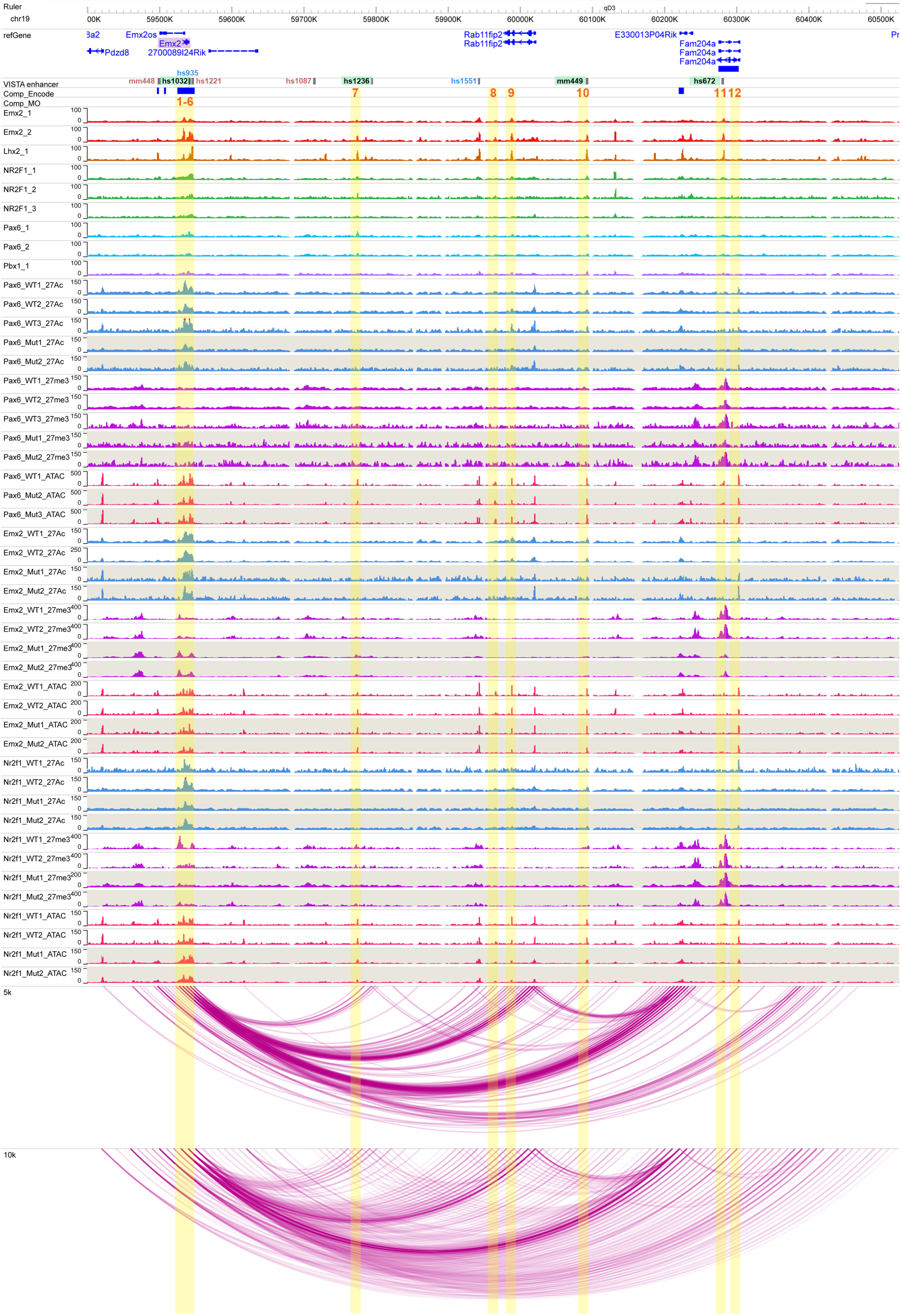

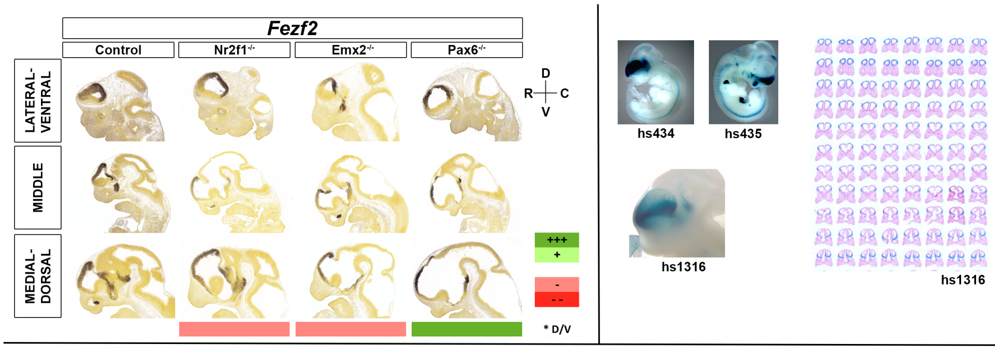


| Fezf2 | chr14:13000000-13370000 | Motifs | | | | | TF ChIP Seq | | | | | Differential epigenomic marks in WT vs. Mutant | | | | | | | | | Plac | Compu tation | Genomic features |
|---|---|---|---|---|---|---|---|---|---|---|---|---|---|---|---|---|---|---|---|---|---|---|---|
| | | | | | | | | | | | | Pax6 mutant | | | Emx2 mutant | | | Nr2f1 mutant | | | | | |
| | | Emx2 HB | Nr2f1 | Pax6 Hb | Pax6 P | Pbx1 | Emx2 | Lhx2 | Nr2f1 | Pax6 | Pbx1 | 27Ac | 27me3 | ATAC | 27Ac | 27me3 | ATAC | 27Ac | 27me3 | ATAC | | | |
| 1 | chr14:13021753-13022951 | x | | x | x | | x | x | | | x | Loss | | Same | Same | | Same | Same | | Same | x | | |
| 2 | chr14:13169800-13171368 | x | | | | | x | x | x | x | x | Loss | | Same | Loss | Gain | Same | Loss | | Same | | | hs434 |
| 3 | chr14:13173323-13174387 | | | | | | x | x | | x | | Same | | Same | Loss | Gain | Loss | Same | | Same | | | |
| 4 | chr14:13177200-13178264 | | | | | | x | x | | x | | Loss | | Loss | Loss | Gain | Same | | | Same | | | Fezf2 TSS |
| 5 | chr14:13178561-13179655 | | | | | | x | x | | | | Same | | Same | Loss | Gain | Same | | | Same | | | hs435 |
| 6 | chr14:13180544-13181786 | x | | x | x | x | x | x | x | x | | | | Loss | | Gain | Loss | | | Same | | | |
| 7 | chr14:13214985-13216615 | | | | | | | | | | | | Same | | | Gain | | | Same | | <10k | | |
| 8 | chr14:13221010-13222735 | | | | | | x | x | x | | | Loss | | Loss | Loss | | Same | Same? | | Same | x | | |

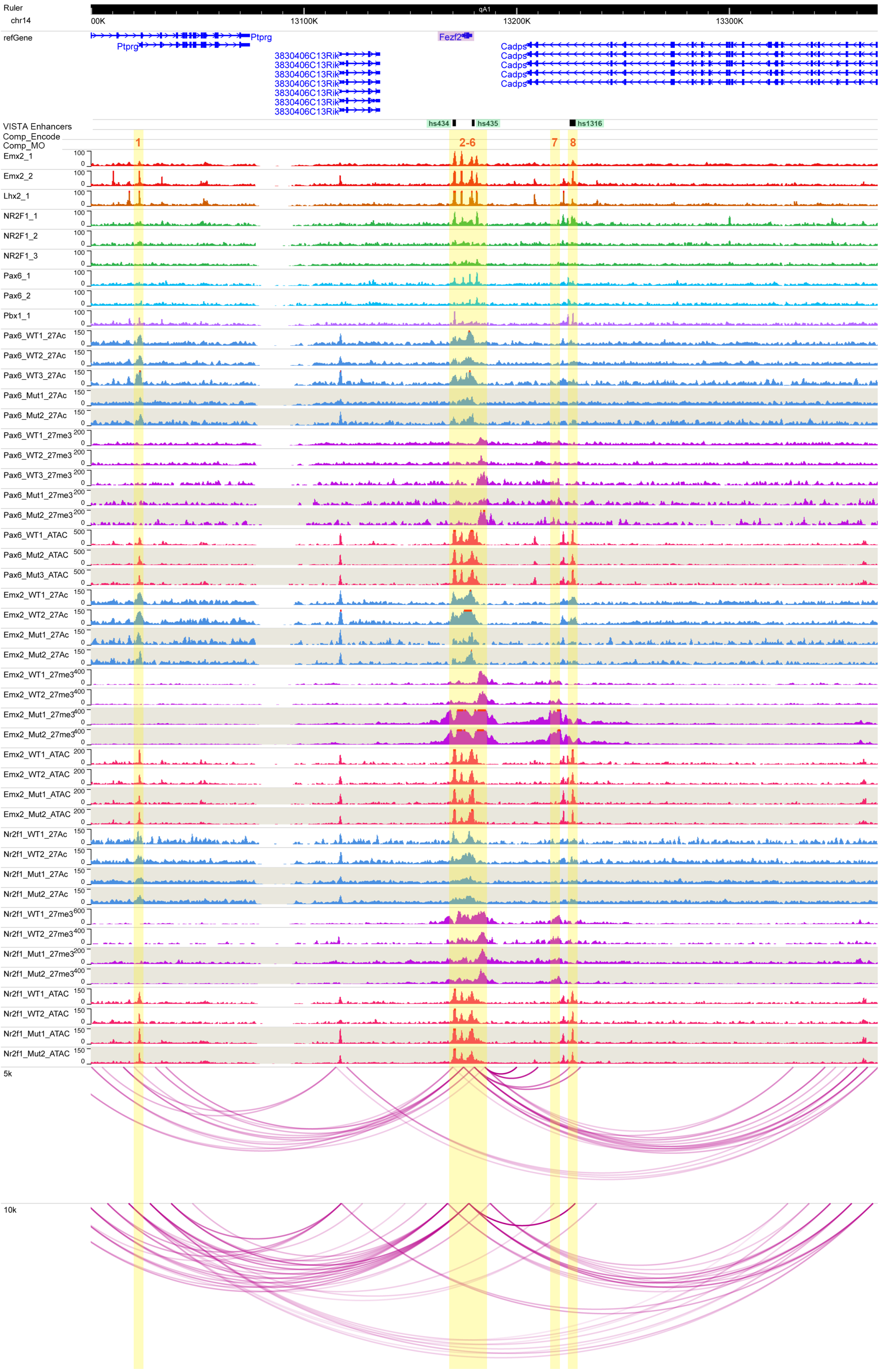

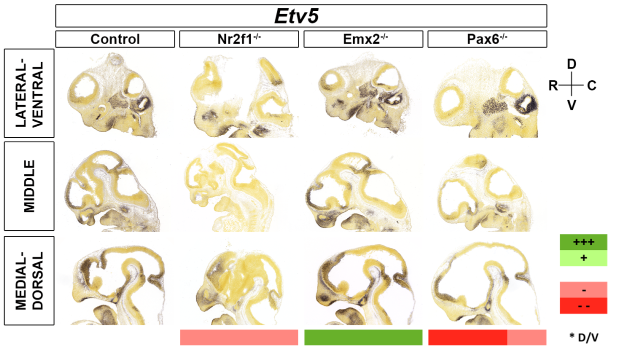


| | | Motifs | | | | | TF ChIP Seq | | | | | Differential epigenomic marks in WT vs. Mutant | | | | | | | | | Plac | Computation | Genomic features |
|---|---|---|---|---|---|---|---|---|---|---|---|---|---|---|---|---|---|---|---|---|---|---|---|
| | | | | | | | | | | | | Pax6 mutant | | | Emx2 mutant | | | Nr2f1 mutant | | | | | |
| | | Emx2 HB | Nr2f1 | Pax6 Hb | Pax6 P | Pbx1 | Emx2 | Lhx2 | Nr2f1 | Pax6 | Pbx1 | 27Ac | 27me3 | ATAC | 27Ac | 27me3 | ATAC | 27Ac | 27me3 | ATAC | | | |
| Etv5 | chr16:22275000-22830000 | | | | | | | | | | | | | | | | | | | | | | |
| 1 | chr16:22427340-22431723 | | | | | x | x | x | | | x | | Same | Loss | | Small Loss | | | Same | | | x | |
| 2 | chr16:22433515-22438593 | | | | | x | x | | | | x | | Same | Same | | Small Loss | | | Same | | | | Etv5 TSS |
| 3 | chr16:22628211-22636846 | | | | | | | | | | | | | | | Small Gain | | | | | | | |
| 4 | chr16:22796861-22801628 | | | | | | | | | x | | | | Same? | | Gain | Gain | | | Same | x | | |

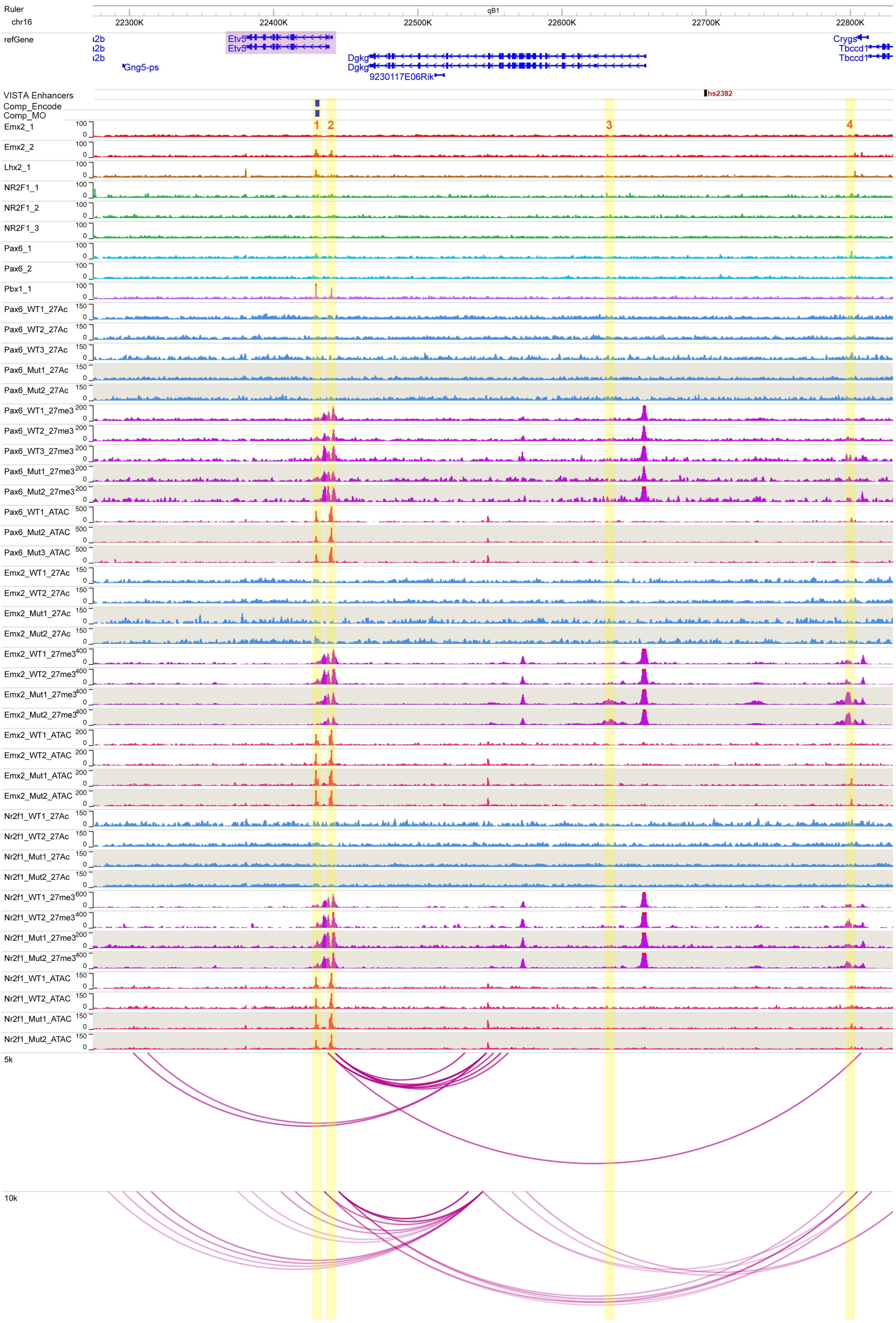

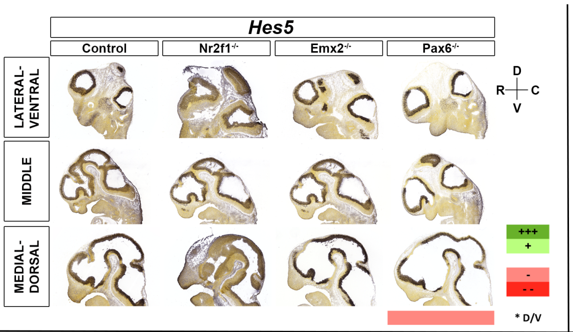


| Hes5 | chr4:154230000-154650000 | Motifs: Emx2 HB | Motifs: Nr2f1 | Motifs: Pax6 Hb | Motifs: Pax6 P | Motifs: Pbx1 | TF ChIP Seq: Emx2 | TF ChIP Seq: Lhx2 | TF ChIP Seq: Nr2f1 | TF ChIP Seq: Pax6 | TF ChIP Seq: Pbx1 | Differential epigenomic marks in WT vs. Mutant — Pax6 mutant: 27Ac | Pax6 mutant: 27me3 | Pax6 mutant: ATAC | Emx2 mutant: 27Ac | Emx2 mutant: 27me3 | Emx2 mutant: ATAC | Nr2f1 mutant: 27Ac | Nr2f1 mutant: 27me3 | Nr2f1 mutant: ATAC | Plac | Computation | Genomic features |
|---|---|---|---|---|---|---|---|---|---|---|---|---|---|---|---|---|---|---|---|---|---|---|---|
| 1 | chr4:154322202-154323976 | | | | | | | | x | | | Loss | | Same | | | Same | Loss | | Same | | x | |
| 2 | chr4:154326550-154327836 | | | | | | x | | x | x | | Loss | | Loss | Same | | Same | Same | | Same | | | |
| 3 | chr4:154334677-154335804 | x | | | | | x | x | | | | Same | | Same | Small Gain? | | Same | | | Same | | x | Hes5 Gene Body |
| 4 | chr4:154607823-154610352 | | | | | | | | x | | | Same | | Loss | Same | | Same | Same | | Same | x | | mm318 |

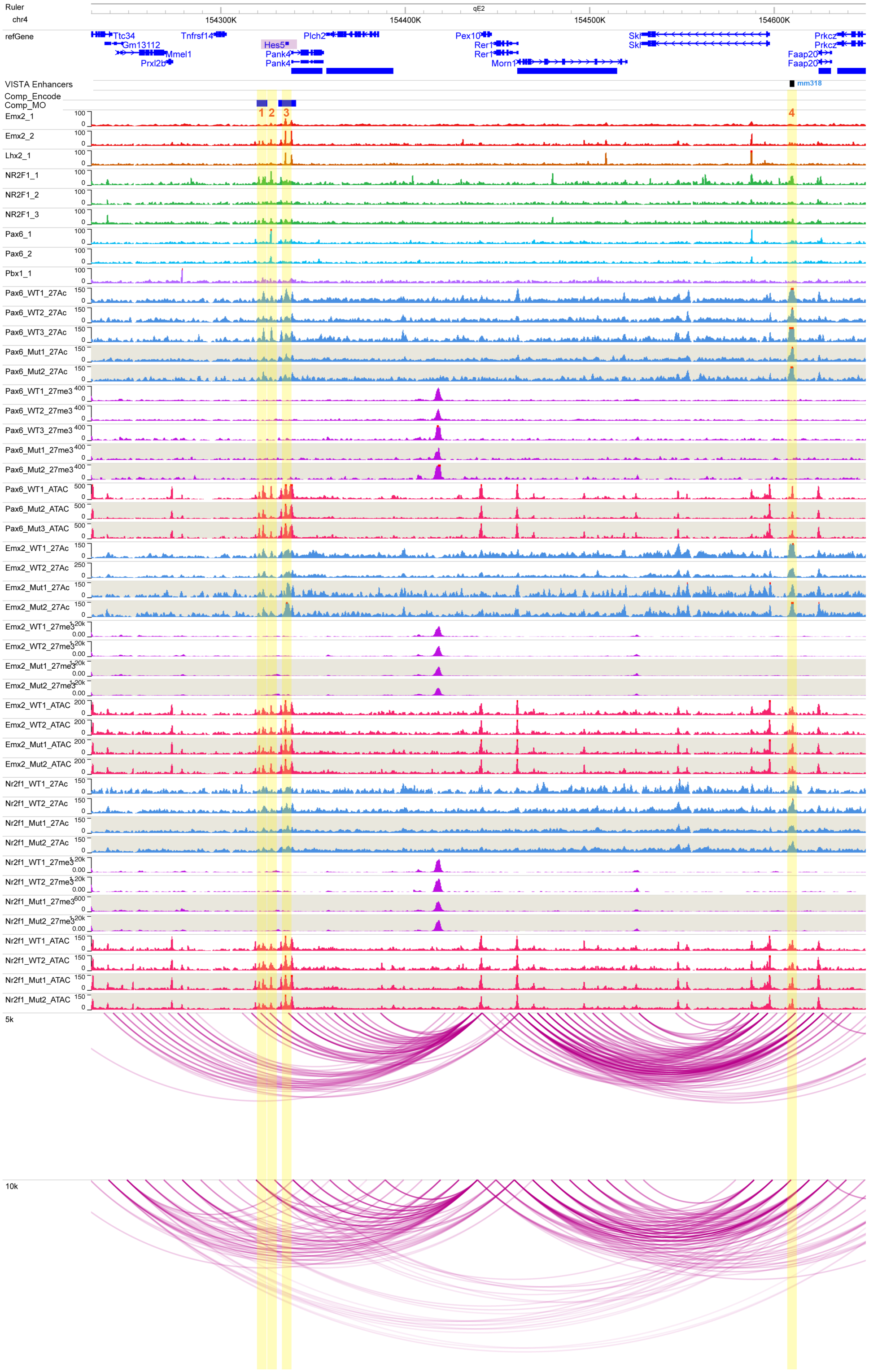

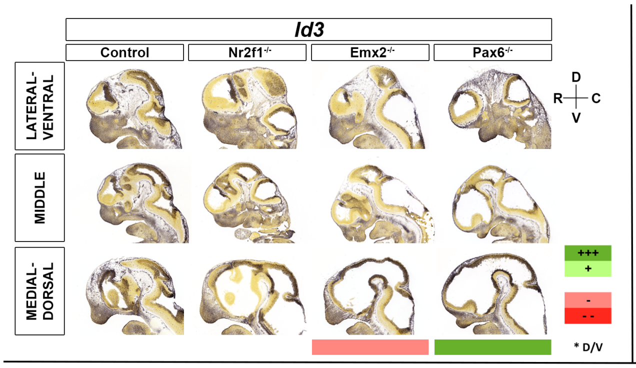


| | | Motifs | | | | | TF ChIP Seq | | | | | Differential epigenomic marks in WT vs. Mutant | | | | | | | | | Plac | Computation | Genomic features |
|---|---|---|---|---|---|---|---|---|---|---|---|---|---|---|---|---|---|---|---|---|---|---|---|
| | | | | | | | | | | | | Pax6 mutant | | | Emx2 mutant | | | Nr2f1 mutant | | | | | |
| | | Emx2 HB | Nr2f1 | Pax6 Hb | Pax6 P | Pbx1 | Emx2 | Lhx2 | Nr2f1 | Pax6 | Pbx1 | 27Ac | 27me3 | ATAC | 27Ac | 27me3 | ATAC | 27Ac | 27me3 | ATAC | | | |
| Id3 | chr4:135590000-135810000 | | | | | | | | | | | | | | | | | | | | | | |
| 1 | chr4:135608849-135609824 | | | | | | | | x | | | Same | | Same | Same | | Same | Loss | | Same | x | | Rpl11 TSS |
| 2 | chr4:135793403-135795370 | | | | | | | | | | | | Loss | | | Same | | | Same | | | x | |

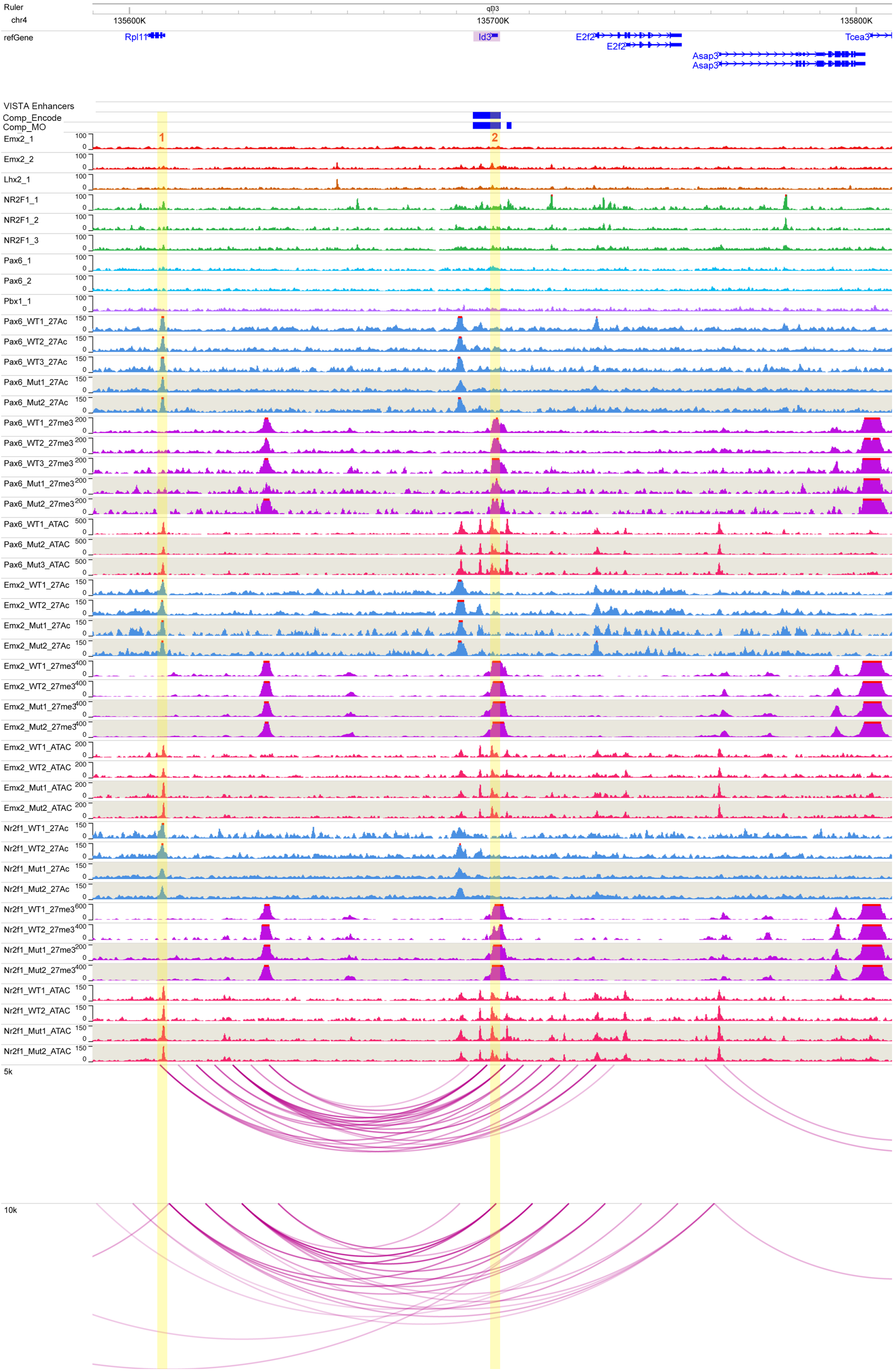

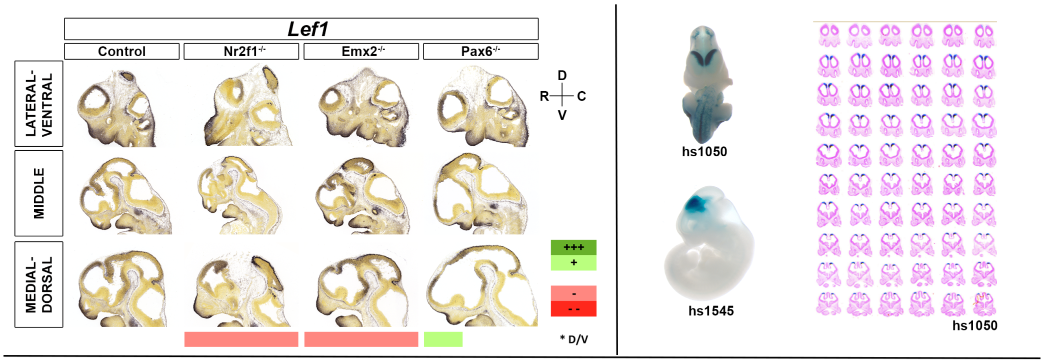


| Lef1 | chr3:130280000-131120000 | Motifs | | | | | TF ChIP Seq | | | | | Differential epigenomic marks in WT vs. Mutant | | | | | | | | | Plac | Compu tation | Genomic features |
|---|---|---|---|---|---|---|---|---|---|---|---|---|---|---|---|---|---|---|---|---|---|---|---|
| | | | | | | | | | | | | Pax6 mutant | | | Emx2 mutant | | | Nr2f1 mutant | | | | | |
| | | Emx2 HB | Nr2f1 | Pax6 Hb | Pax6 P | Pbx1 | Emx2 | Lhx2 | Nr2f1 | Pax6 | Pbx1 | 27Ac | 27me3 | ATAC | 27Ac | 27me3 | ATAC | 27Ac | 27me3 | ATAC | | | |
| 1 | chr3:130664999-130667081 | x | | x | x | x | x | | x | x | | Loss | | Loss | Same | | Same | Same | | Same | x | | hs1545 |
| 2 | chr3:130774287-130776523 | | | | | | x | x | x | x | | | | Same | | | Loss | | | Same | | | |
| 3 | chr3:130786473-130789557 | | | | | | | | | | | | Loss | Same | | Same | Same | | Same | Same | | | |
| 4 | chr3:130809043-130812870 | | | | | | | | | | | | Same | Same | | Gain | Same | | Same | Same | | | Lef1 Promoter |
| 5 | chr3:131035570-131040367 | x | | | | | x | x | | x | x | | Same | Same | | Loss | Loss | | Same | Same | | | Cyp2u1 |

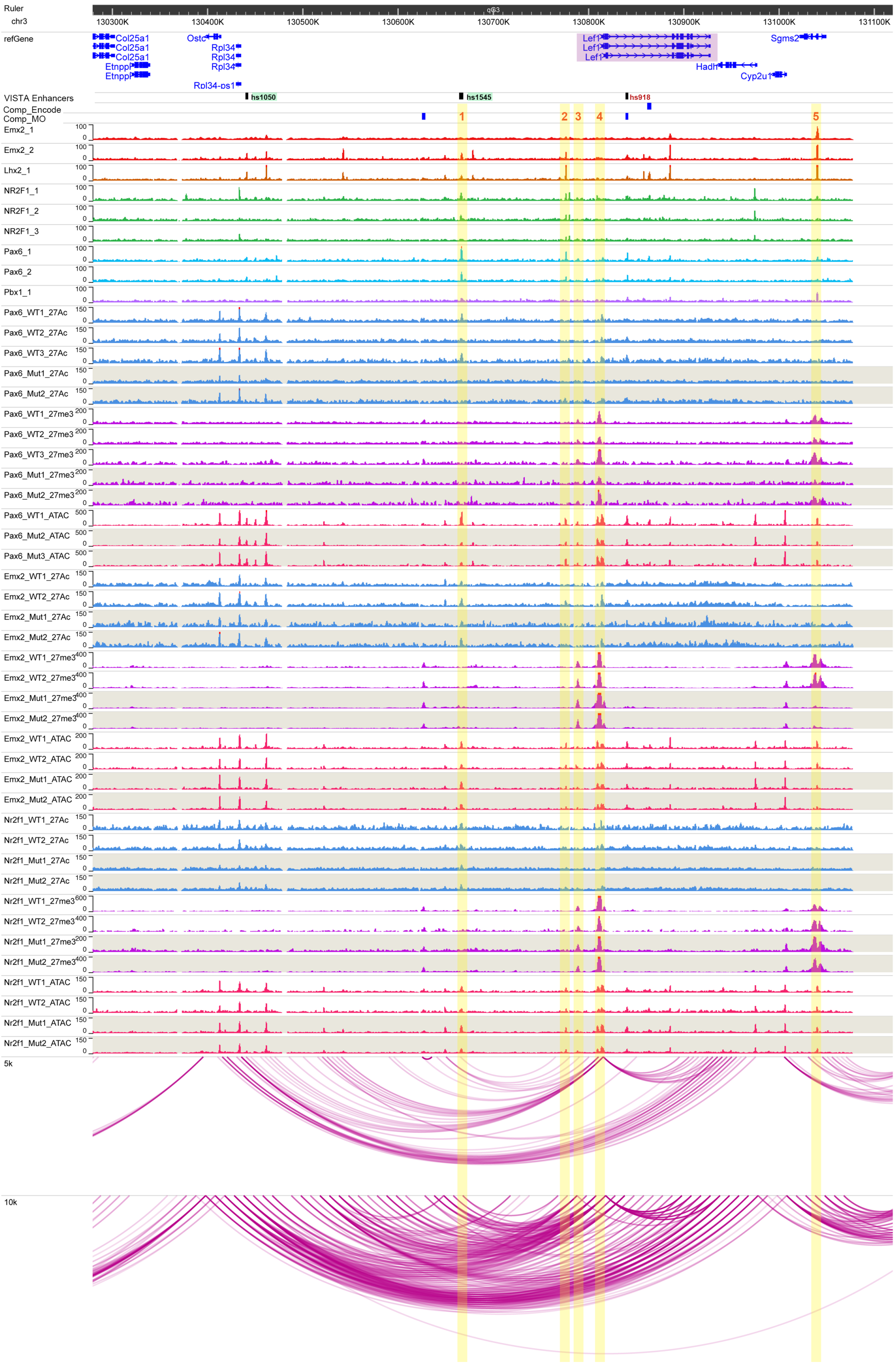

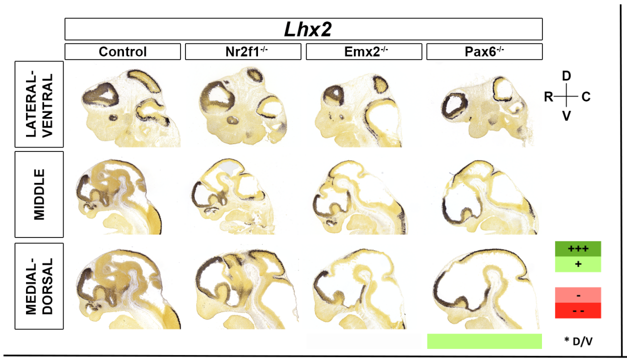


| | | Motifs | | | | | TF ChIP Seq | | | | | Differential epigenomic marks in WT vs. Mutant | | | | | | | | | Plac | Computation | Genomic features |
|---|---|---|---|---|---|---|---|---|---|---|---|---|---|---|---|---|---|---|---|---|---|---|---|
| | | | | | | | | | | | | Pax6 mutant | | | Emx2 mutant | | | Nr2f1 mutant | | | | | |
| | | Emx2 HB | Nr2f1 | Pax6 Hb | Pax6 P | Pbx1 | Emx2 | Lhx2 | Nr2f1 | Pax6 | Pbx1 | 27Ac | 27me3 | ATAC | 27Ac | 27me3 | ATAC | 27Ac | 27me3 | ATAC | | | |
| Lhx2 | chr2:37560000-38600000 | | | | | | | | | | | | | | | | | | | | | | |
| 1 | chr2:37997193-38001351 | x | | | | x | x | | x | | x | Loss | | Same | Loss | | Same | Same | | Same | | | hs314 |
| 2 | chr2:38048398-38056606 | | | | | | x | x | x | | | | | Same | | Gain | Same | | | Same | | | |
| 3 | chr2:38194392-38197269 | x | | | | | x | x | x | | x | Loss | | Same | Loss | Gain | Same | Loss | | Same | | | Lhx2 TSS |
| 4 | chr2:38202560-38207368 | x | | | | x | x | x | x | | x | Loss | | Same | Loss | Gain | Same | Loss | | Same | | | Lhx2 alternative TSS |
| 5 | chr2:38370512-38372007 | | | | | | | | x | | x | Loss | | Loss | Loss | | Same | | | Same | | | Nek6 TSS |

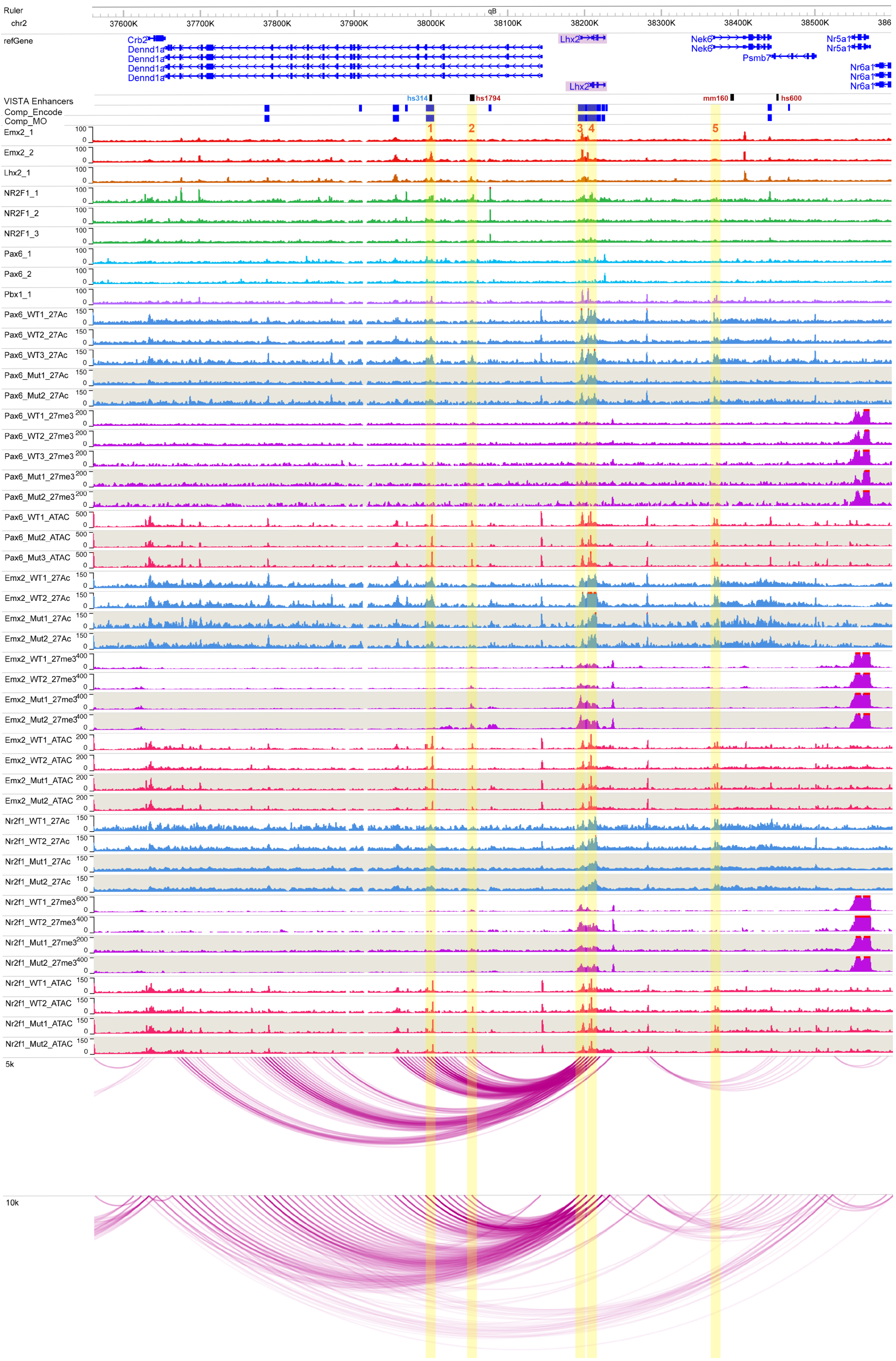

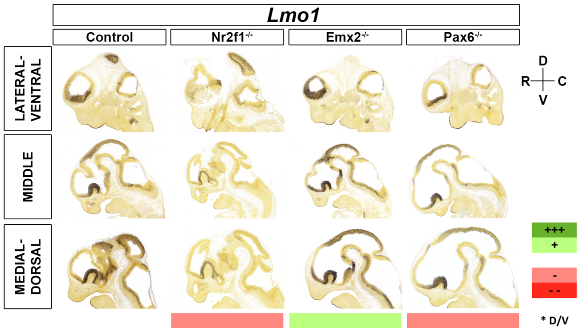


| | | Motifs | | | | | TF ChIP Seq | | | | | Differential epigenomic marks in WT vs. Mutant | | | | | | | | | Plac | Computation | Genomic features |
|---|---|---|---|---|---|---|---|---|---|---|---|---|---|---|---|---|---|---|---|---|---|---|---|
| | | | | | | | | | | | | Pax6 mutant | | | Emx2 mutant | | | Nr2f1 mutant | | | | | |
| | | Emx2 HB | Nr2f1 | Pax6 Hb | Pax6 P | Pbx1 | Emx2 | Lhx2 | Nr2f1 | Pax6 | Pbx1 | 27Ac | 27me3 | ATAC | 27Ac | 27me3 | ATAC | 27Ac | 27me3 | ATAC | | | |
| Lmo1 | chr7:116182079-116670000 | | | | | | | | | | | | | | | | | | | | | | |
| 1 | chr7:116254369-116260805 | | | | | | x | x | x | x | | | Same | | | Loss | | | Same | | | | hs1859 |
| 2 | chr7:116269340-116274412 | x | | | | | x | x | | | | | Same | | | Loss | | | Same | | | | |
| 3 | chr7:116285437-116289362 | x | | | | | x | x | x | | | | Same | Loss | | Loss | Loss | | Same | | | | Lmo1 Gene Body |
| 4 | chr7:116307961-116310857 | x | | | | | x | x | | x | | | Same | Loss | | Loss | Same | | Same | Same | | | Lmo1 Gene Body |
| 5 | chr7:116310160-116318282 | | | | | | x | | | | | | Same | | | Loss | | | Same | | | | Lmo1 TSS |
| 6 | chr7:116318810-116326219 | x | | x | x | | x | x | x | x | x | Loss | Same | Same | Gain | Loss | Same | Loss | Same | Same | | x | |
| 7 | chr7:116333909-116336398 | x | | | | | x | x | x | x | x | Loss | Same | Same | Same | Loss | Same | Loss | Same | Same | | | |
| 8 | chr7:116344968-116348740 | | | | | | | | | | | | Same | | | Loss | | | Same | | | | |
| 9 | chr7:116581666-116584183 | | | | | | | | | | | | Same | | Gain | Loss | | | Same | | x | | |

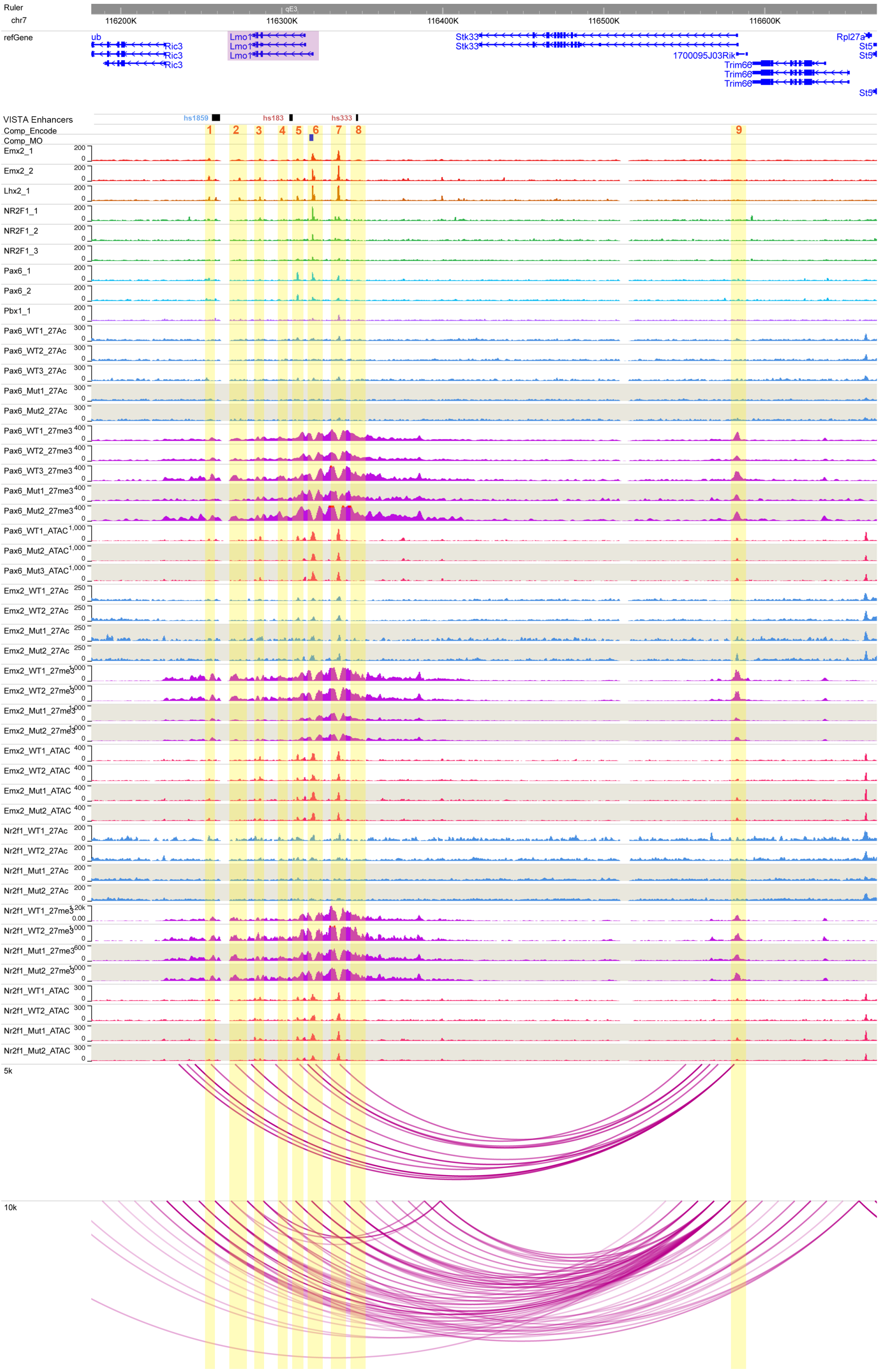

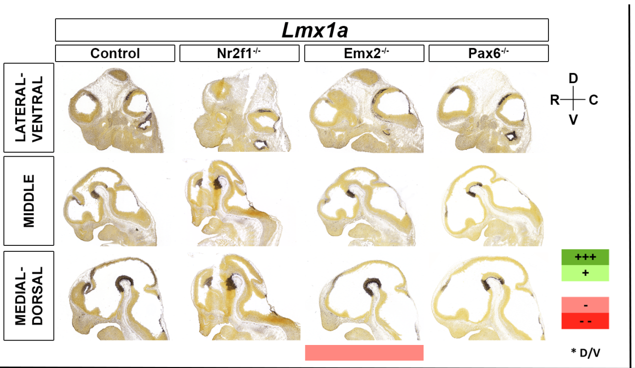


| | | Motifs | | | | | TF ChIP Seq | | | | | Differential epigenomic marks in WT vs. Mutant | | | | | | | | | Plac | Computation | Genomic features |
|---|---|---|---|---|---|---|---|---|---|---|---|---|---|---|---|---|---|---|---|---|---|---|---|
| | | | | | | | | | | | | Pax6 mutant | | | Emx2 mutant | | | Nr2f1 mutant | | | | | |
| | | Emx2 HB | Nr2f1 | Pax6 Hb | Pax6 P | Pbx1 | Emx2 | Lhx2 | Nr2f1 | Pax6 | Pbx1 | 27Ac | 27me3 | ATAC | 27Ac | 27me3 | ATAC | 27Ac | 27me3 | ATAC | | | |
| Lmx1a | chr1:169280000-169880000 | | | | | | | | | | | | | | | | | | | | | | |
| 1 | chr1:169527243-169530622 | | | | | | | | | | | | Same | | | Loss | | | Same | | | | |
| 2 | chr1:169589344-169592457 | | | | | | | | | | | | Same | | | Gain | | | Loss | | | | |
| 3 | chr1:169601564-169604678 | | | | | | | | | | | | Same | | | Same | | | Loss | | | | |
| 4 | chr1:169617808-169623158 | | | | | | | | x | | | | Same | Small Loss | | Same | Small loss | | Same | Same | | | |
| 5 | chr1:169731979-169733692 | | | x | x | | x | x | | | | Loss | | Same | Same | | Same | Same | | Same | | | |

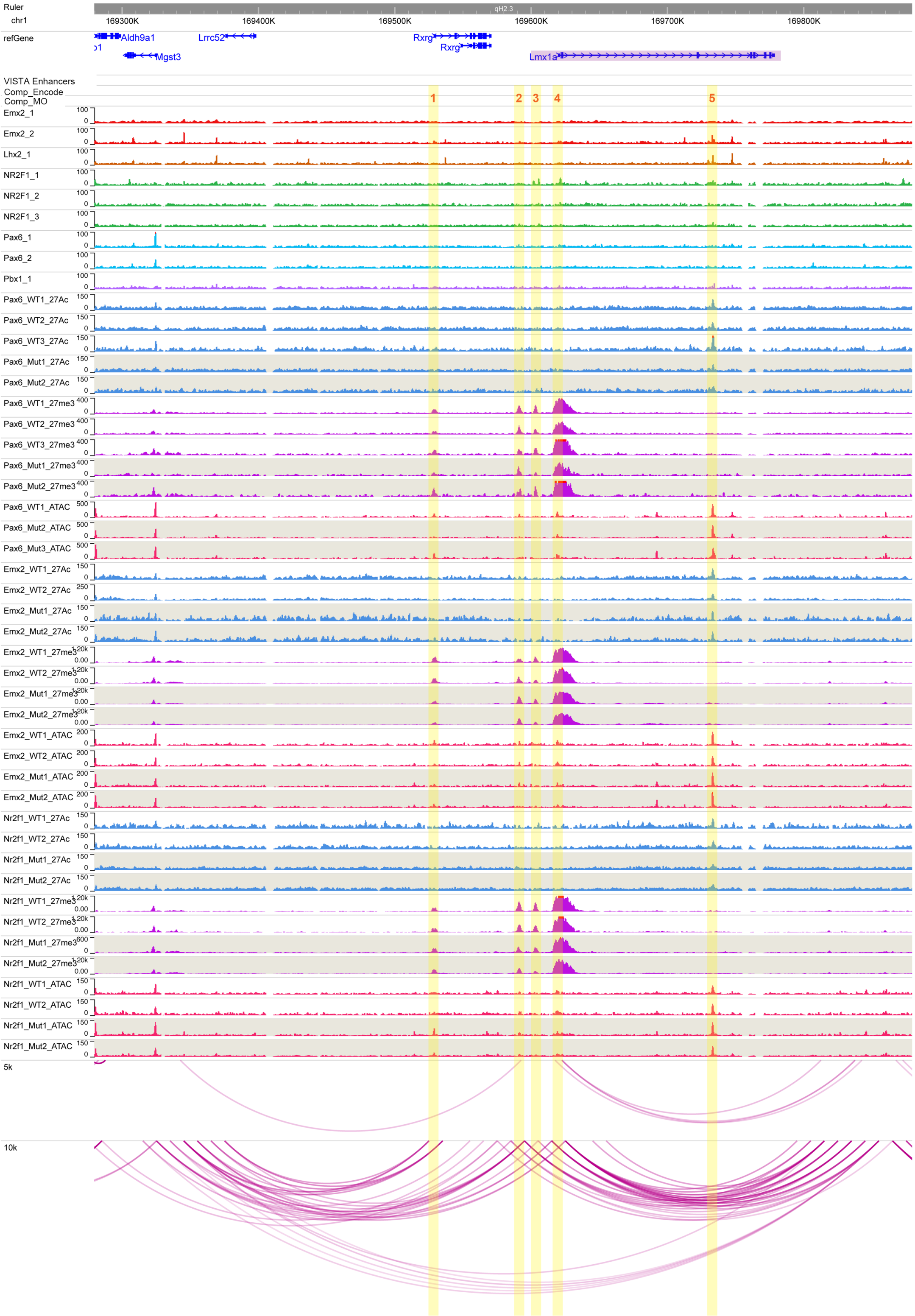

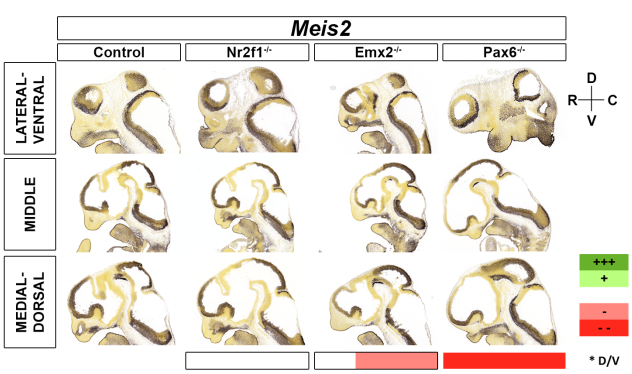


| | | Motifs | | | | | TF ChIP Seq | | | | | Differential epigenomic marks in WT vs. Mutant | | | | | | | | | Plac-Seq | Computation | Genomic features |
|---|---|---|---|---|---|---|---|---|---|---|---|---|---|---|---|---|---|---|---|---|---|---|---|
| | | | | | | | | | | | | Pax6 mutant | | | Emx2 mutant | | | Nr2f1 mutant | | | | | |
| | | Emx2 HB | Nr2f1 | Pax6 Hb | Pax6 P | Pbx1 | Emx2 | Lhx2 | Nr2f1 | Pax6 | Pbx1 | 27Ac | 27me3 | ATAC | 27Ac | 27me3 | ATAC | 27Ac | 27me3 | ATAC | | | |
| Meis2 | chr2:114600000-116700000 | | | | | | | | | | | | | | | | | | | | | | |
| 1 | chr2:114957467-114958808 | x | | x | x | x | x | x | x | x | x | Loss | | Same | Loss | | Same | Loss | | Same | x | | |
| 2 | chr2:115144553-115152886 | | | | | | | | | | | | Small gain | | | Gain | | | Same | | | | |
| 3 | chr2:115281249-115282302 | x | | | | | x | x | x | x | x | Loss | | Same | Loss | | Loss | Same | | Same | x | | |
| 4 | chr2:115293609-115294718 | x | | x | x | | x | x | | x | | Same | | Loss | Loss | | Loss | Same | | Same | x | | |
| 5 | chr2:115482362-115483493 | x | | | | | x | x | x | | x | Same | | Same | Loss | | Loss | Same | | Same | x | | hs812 |
| 6 | chr2:115485397-115487093 | x | | | | | x | x | x | x | x | Loss | | Same | Loss | | Loss | Loss | | Same | x | x | |
| 7 | chr2:115550455-115552059 | x | | x | x | | x | x | x | | x | Loss | | Same | Loss | | Loss | Same | | Same | x | x | |
| 8 | chr2:115677352-115681367 | | | | | | | | | | | | Gain | | | Gain | | | Same | | x | x | Meis2 5' UTR |
| 9 | chr2:115897214-115902068 | x | | | x | | x | x | x | | | | Gain | Same | | Gain | Same | | Same | Same | | x | Near Meis2 TSS |

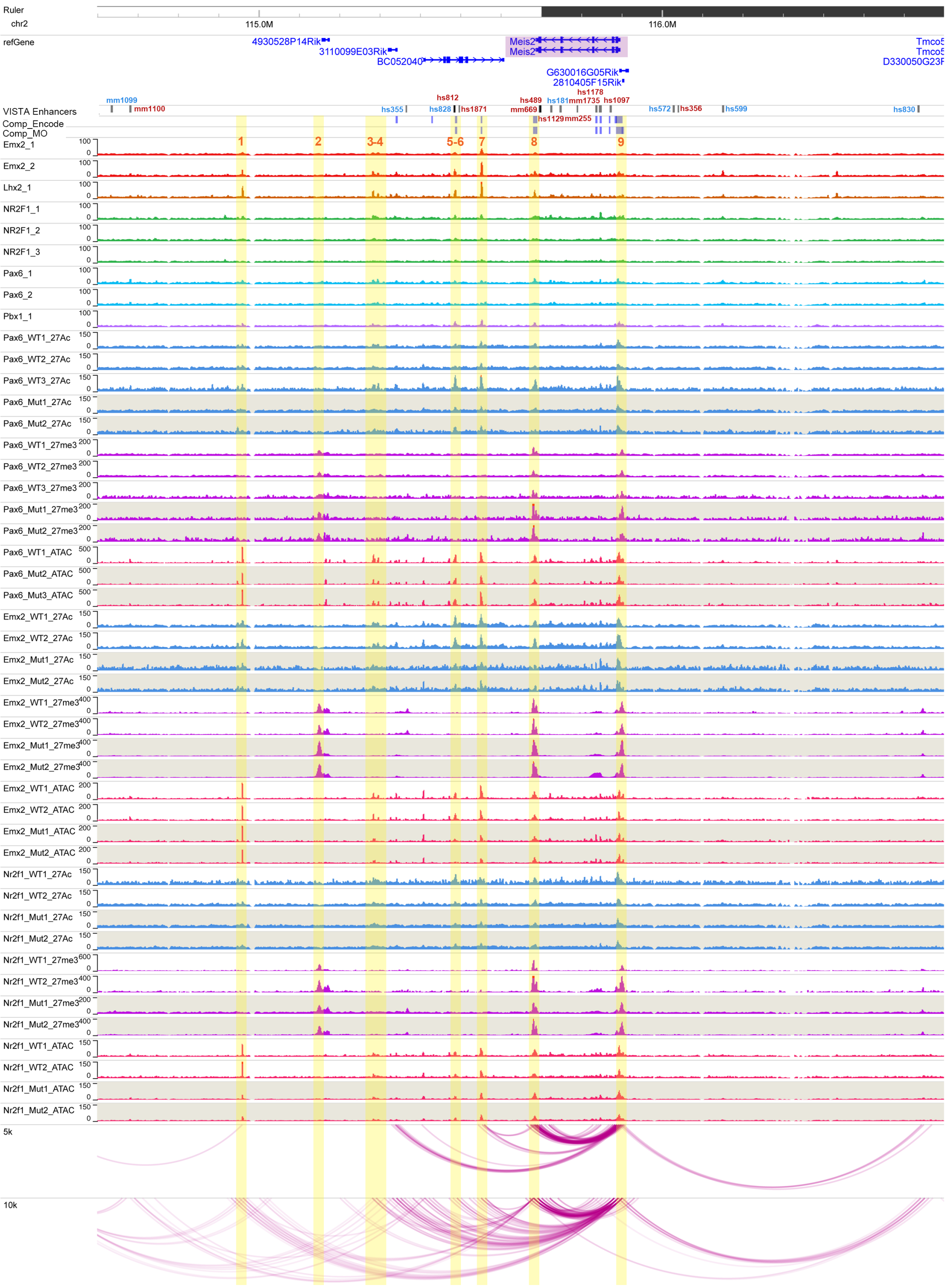

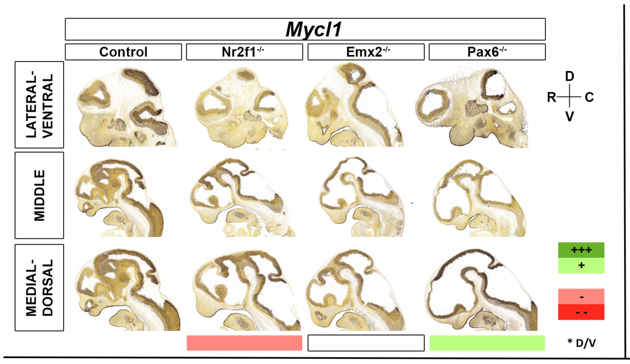


| | | Motifs | | | | | TF ChIP Seq | | | | | Differential epigenomic marks in WT vs. Mutant | | | | | | | | | Plac-Seq | Computation | Genomic features |
|---|---|---|---|---|---|---|---|---|---|---|---|---|---|---|---|---|---|---|---|---|---|---|---|
| | | | | | | | | | | | | Pax6 mutant | | | Emx2 mutant | | | Nr2f1 mutant | | | | | |
| | | Emx2 HB | Nr2f1 | Pax6 Hb | Pax6 P | Pbx1 | Emx2 | Lhx2 | Nr2f1 | Pax6 | Pbx1 | 27Ac | 27me3 | ATAC | 27Ac | 27me3 | ATAC | 27Ac | 27me3 | ATAC | | | |
| Mycl1 | chr4:122560000-122800000 | | | | | | | | | | | | | | | | | | | | | | |
| 1 | chr4:122672621-122674388 | | | | | | | | | | | Loss | | Same | Same | | Same | Loss | | Same | | x | Mycl1 TSS |
| 2 | chr4:122690008-122692021 | | | | | | | | | | | Loss | | Same | Same | | Same | Same | | Same | | x | |
| 3 | chr4:122692924-122694514 | | | | | | | | | | | Loss | | Same | Same | | Same | Same | | Same | | x | Trit1 TSS |
| 4 | chr4:122727680-122729103 | | | | | | | | x | | | Same | | Same | Same | | Same | Loss | | Same | | x | Trit1 Gene body |

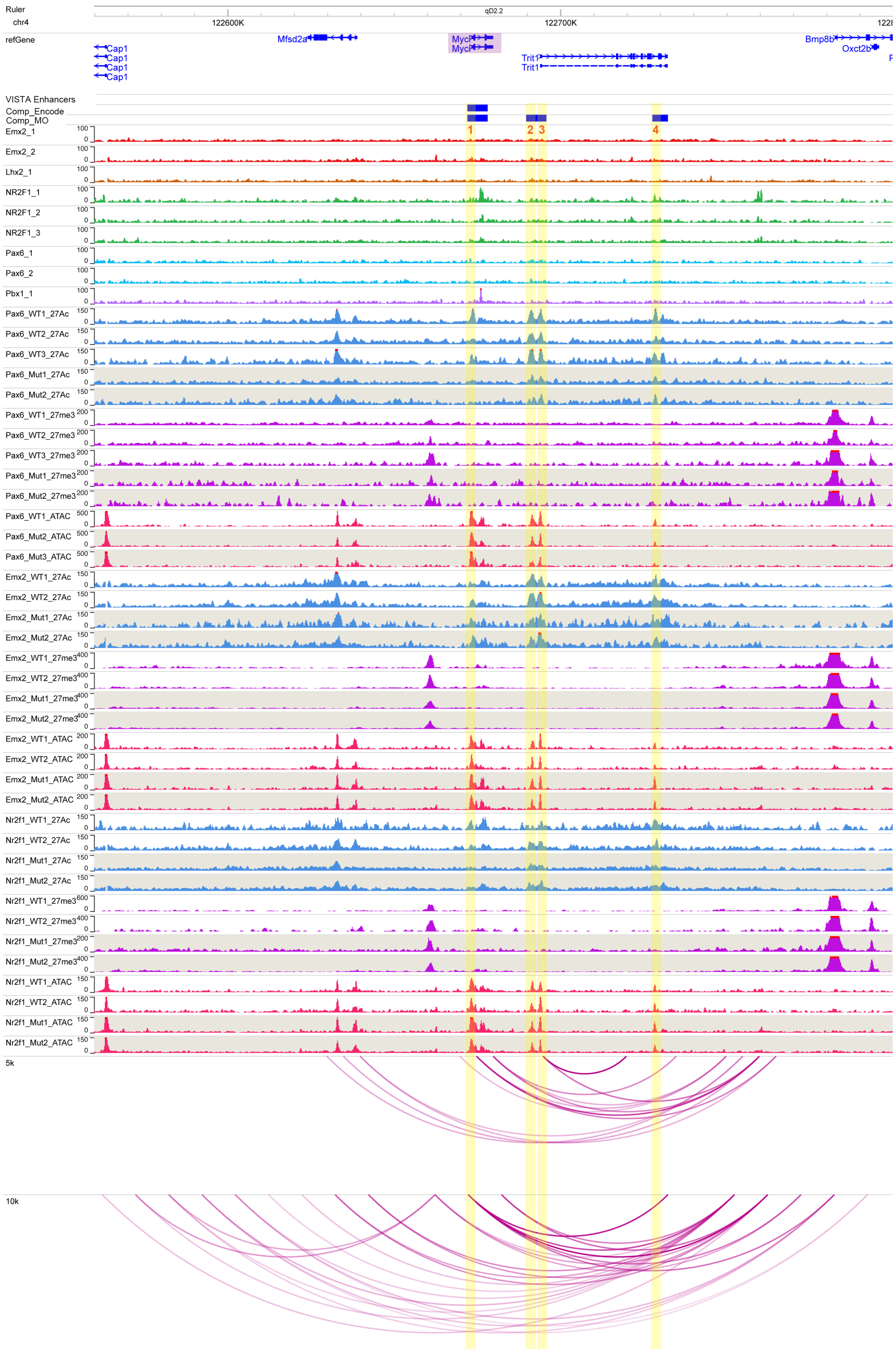

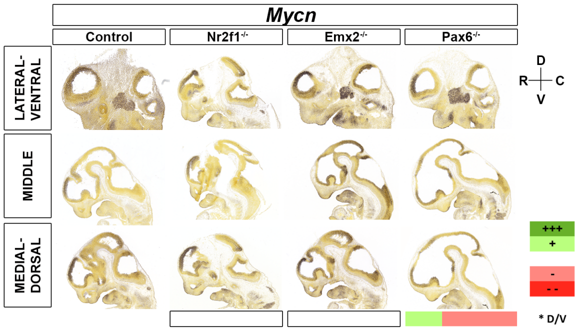


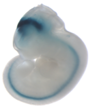


| | | Motifs | | | | | TF ChIP Seq | | | | | Differential epigenomic marks in WT vs. Mutant | | | | | | | | | Plac-Seq | Compu tation | Genomic features |
|---|---|---|---|---|---|---|---|---|---|---|---|---|---|---|---|---|---|---|---|---|---|---|---|
| | | | | | | | | | | | | Pax6 mutant | | | Emx2 mutant | | | Nr2f1 mutant | | | | | |
| | | Emx2 HB | Nr2f1 | Pax6 Hb | Pax6 P | Pbx1 | Emx2 | Lhx2 | Nr2f1 | Pax6 | Pbx1 | 27Ac | 27me3 | ATAC | 27Ac | 27me3 | ATAC | 27Ac | 27me3 | ATAC | | | |
| **Mycn** | chr12:12285000-13290000 | | | | | | | | | | | | | | | | | | | | | | |
| **1** | chr12:12514739-12516576 | x | | x | x | | x | x | x | | | Loss | | Loss | Loss | | Loss | Loss | | Same | x | | |
| **2** | chr12:12532797-12535809 | x | | | | | x | x | x | | x | Same | | Same | Loss | | Loss | Same | | Same | x | x | |
| **3** | chr12:12587394-12589056 | | | | | | x | | x | | x | Same? | | Same | Same | | Same | Loss | | Same | x | x | |
| **4** | chr12:12690102-12692253 | | | | | | | | | | | Loss | | Same | Loss | | Same | Loss | | Same | x | | |
| **5** | chr12:12745097-12746653 | x | | | | | x | x | x | x | | Loss | | Same | Loss | | Same | Loss | | Same | x | x | |
| **6** | chr12:12789957-12791513 | x | | | | | x | x | x | | | Same | | Same | Loss | | Same | Same | | Same | x | | |
| **7** | chr12:12910817-12912373 | | | | | | | | x | | | Loss | | Same | Same | | Same | Same | | Same | | | |
| **8** | chr12:12943665-12946290 | x | | | | | x | | x | | | Loss | | Same | Same | | Same | Loss | | Same | | x | Mycn Gene body |
| **9** | chr12:12948261-12950492 | | | | | | x | | x | | | Loss | | Same | Same | | Same | Loss | | Same | | x | Mycn Gene body |
| **10** | chr12:13017470-13019026 | x | | | | | x | x | | | x | | | Loss | | | Loss | | | Same | | | |
| **11** | chr12:13215205-13216274 | | | | | | | | x | x | | | | Loss | | | Same | | | Same | x | | |

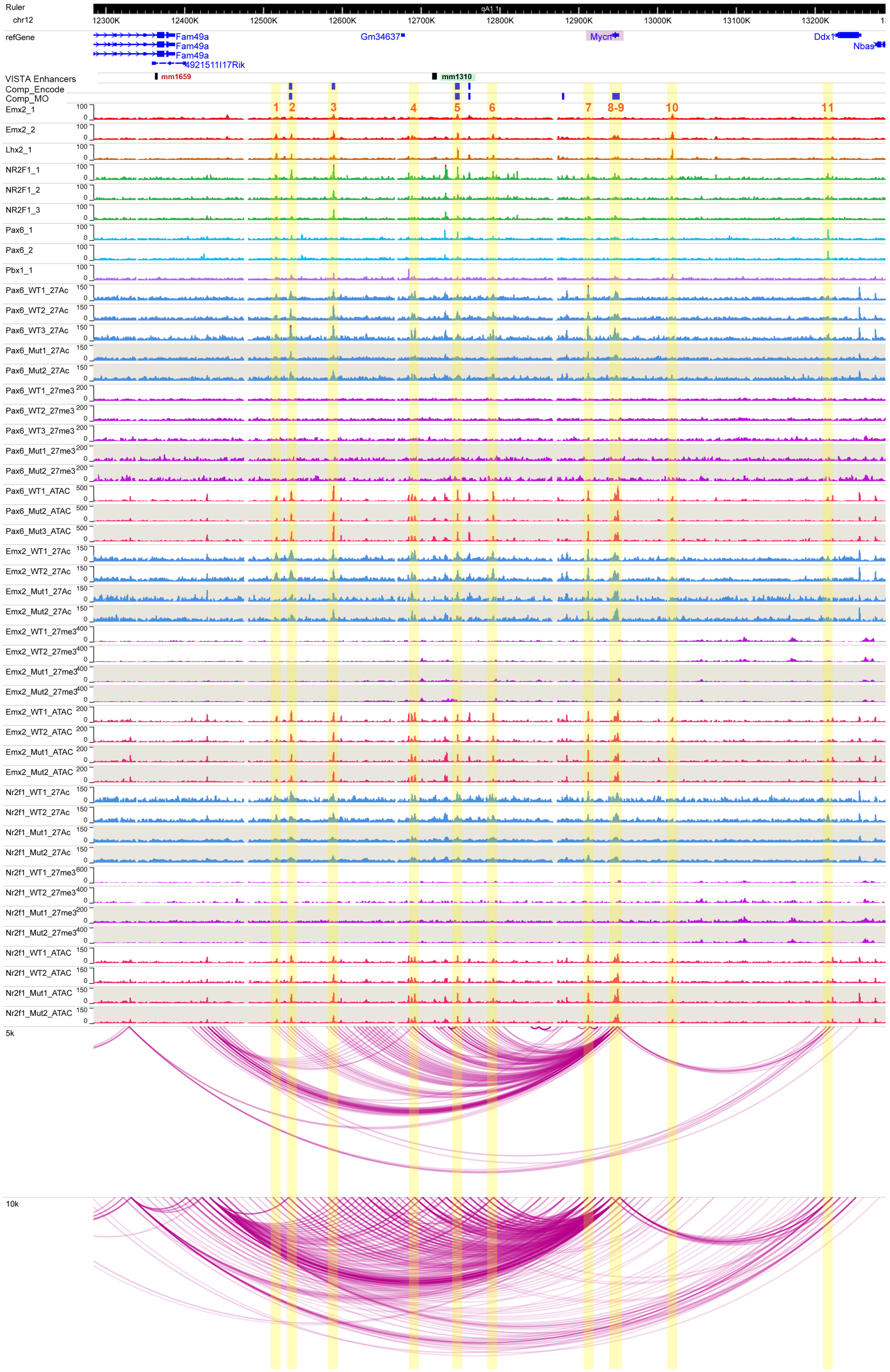

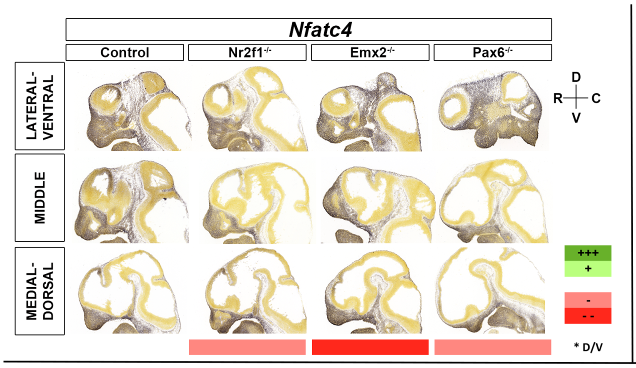


| | | Motifs | | | | | TF ChIP Seq | | | | | Differential epigenomic marks in WT vs. Mutant | | | | | | | | | Plac-Seq | Computation | Genomic features |
|---|---|---|---|---|---|---|---|---|---|---|---|---|---|---|---|---|---|---|---|---|---|---|---|
| | | | | | | | | | | | | Pax6 mutant | | | Emx2 mutant | | | Nr2f1 mutant | | | | | |
| | | Emx2 HB | Nr2f1 | Pax6 Hb | Pax6 P | Pbx1 | Emx2 | Lhx2 | Nr2f1 | Pax6 | Pbx1 | 27Ac | 27me3 | ATAC | 27Ac | 27me3 | ATAC | 27Ac | 27me3 | ATAC | | | |
| Nfatc4 | chr14:56343632-56560000 | | | | | | | | | | | | | | | | | | | | | | |
| 1 | chr14:56440792-56446602 | | | | | | | | x | | | | | | | Gain | | | | | | | Nfatc4 TSS |

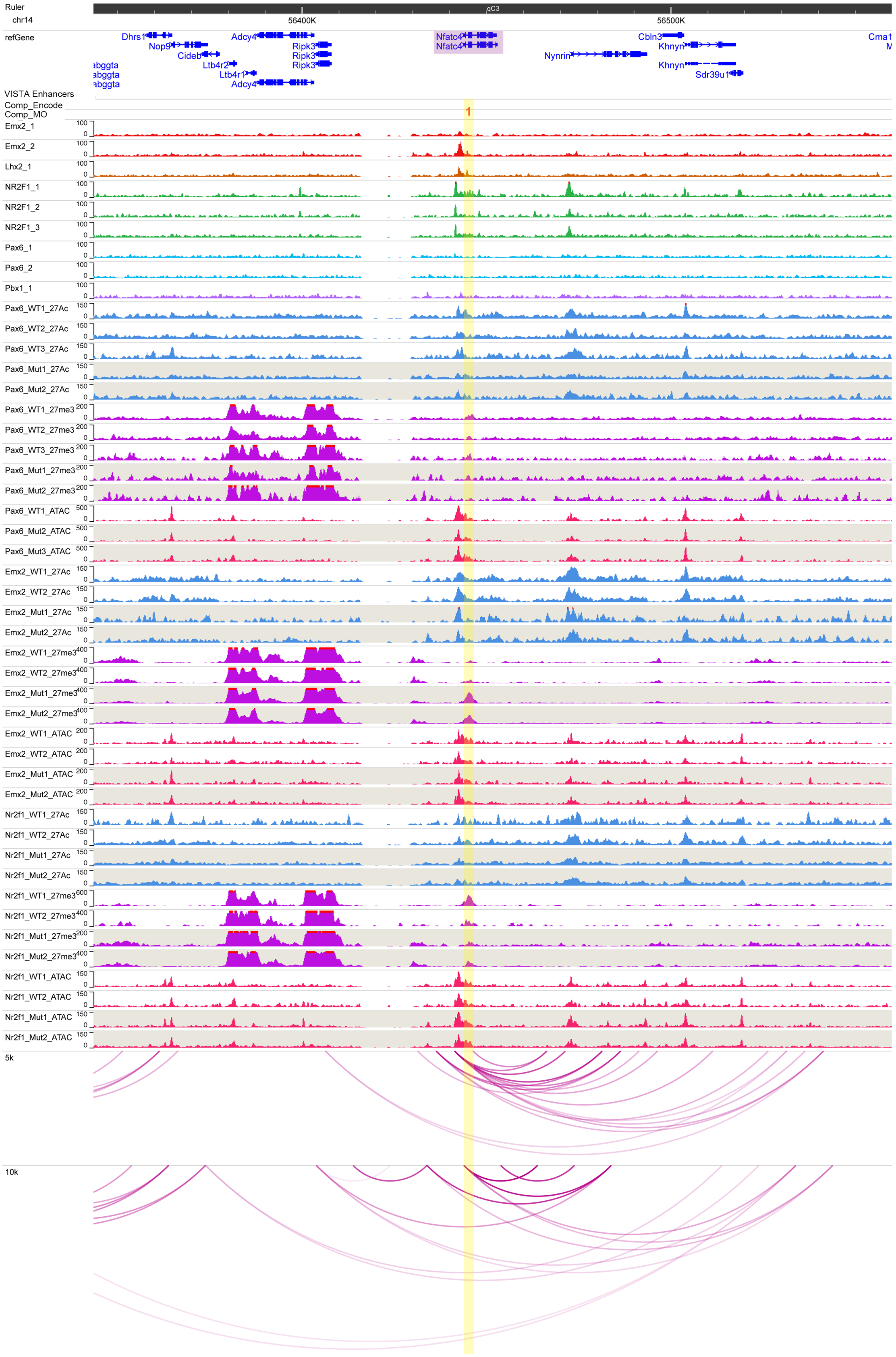

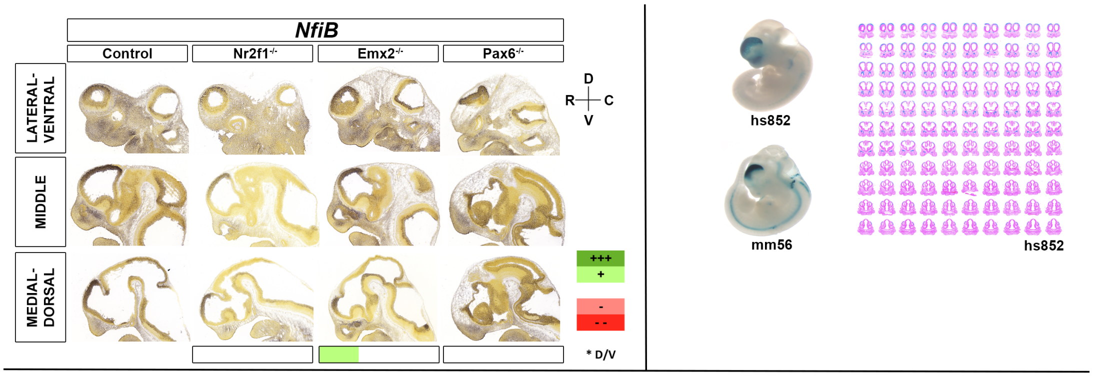


| | | Motifs | | | | | TF ChIP Seq | | | | | Differential epigenomic marks in WT vs. Mutant | | | | | | | | | Plac-Seq | Computation | Genomic features |
|---|---|---|---|---|---|---|---|---|---|---|---|---|---|---|---|---|---|---|---|---|---|---|---|
| | | | | | | | | | | | | Pax6 mutant | | | Emx2 mutant | | | Nr2f1 mutant | | | | | |
| | | Emx2 HB | Nr2f1 | Pax6 Hb | Pax6 P | Pbx1 | Emx2 | Lhx2 | Nr2f1 | Pax6 | Pbx1 | 27Ac | 27me3 | ATAC | 27Ac | 27me3 | ATAC | 27Ac | 27me3 | ATAC | | | |
| NfiB | chr4:81100000-82900000 | | | | | | | | | | | | | | | | | | | | | | |
| 1 | chr4:81270987-81272556 | | | | | | x | x | x | | | | | Same | | | Loss | | | Same | x | | |
| 2 | chr4:81673006-81674575 | x | | | | | x | x | | | x | | | Same | | | Gain | | | Same | | | |
| 3 | chr4:81949394-81952730 | | | x | | | x | | x | x | | Loss | | Same | Same | | Same | Same | | Same | x | | |
| 4 | chr4:82029483-82031642 | x | | | | | x | | | | | Same | | Same | Loss | | Loss | Same | | Same | | x | |
| 5 | chr4:82150505-82154090 | x | | x | | | x | x | x | x | x | Loss | | Same | Gain | | Same | Same | | Same | | x | Nfib TSS |
| 6 | chr4:82181026-82186913 | x | | | | | x | | x | | x | | Same | | | Gain | | | Loss | | | x | |
| 7 | chr4:82525419-82535633 | x | | | | | x | x | | | | | | | | Gain | | | | | | | Cer1 Gene Body |
| 8 | chr4:82561255-82563610 | x | | | | | x | x | | | x | Same | | Same | Loss | | Loss | Same | | Same | x | | mm56 |

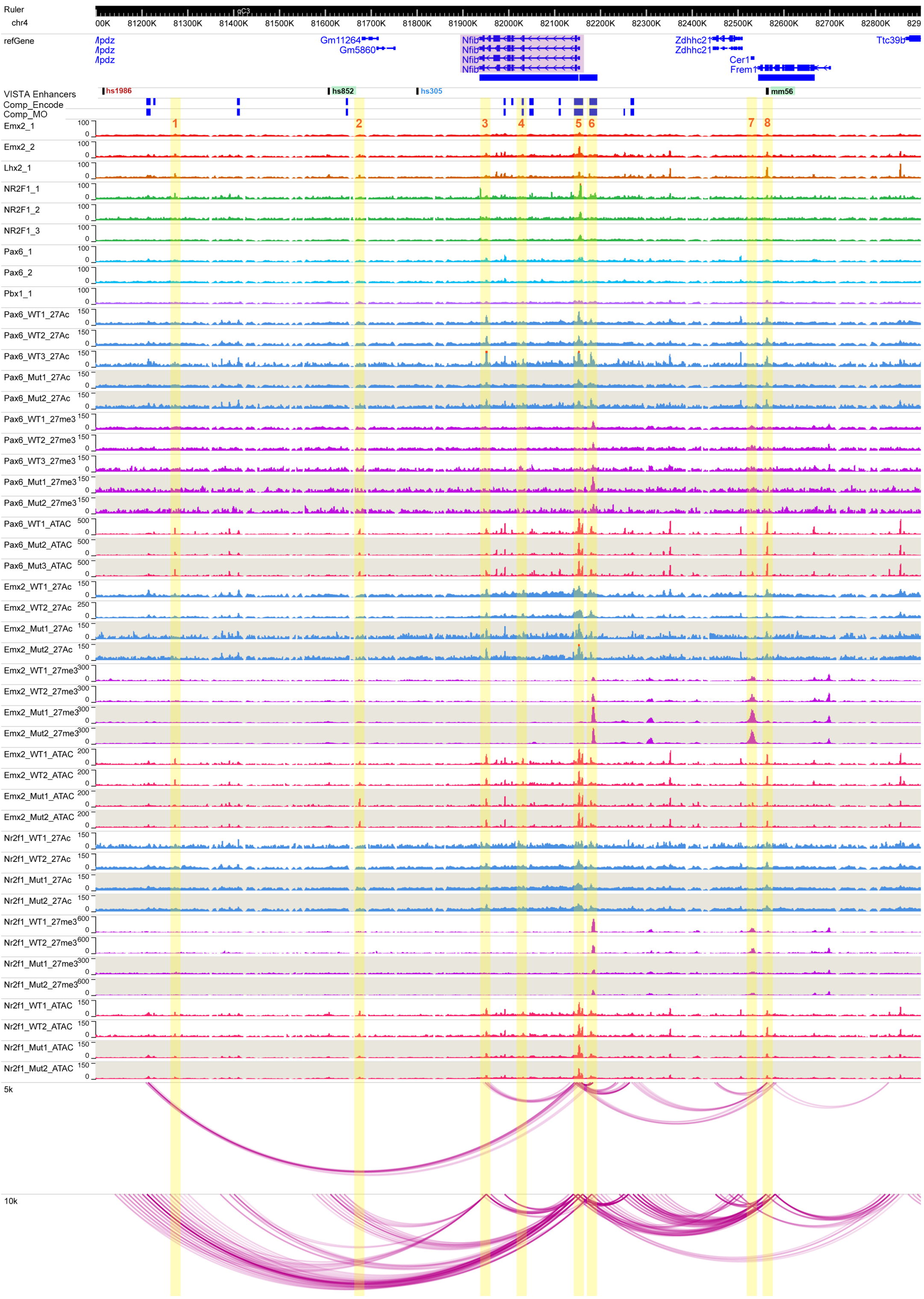

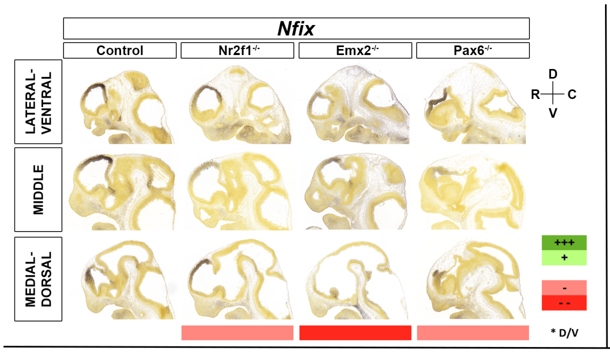


| | | Motifs | | | | | TF ChIP Seq | | | | | Differential epigenomic marks in WT vs. Mutant | | | | | | | | | Plac-Seq | Computation | Genomic features |
|---|---|---|---|---|---|---|---|---|---|---|---|---|---|---|---|---|---|---|---|---|---|---|---|
| | | | | | | | | | | | | Pax6 mutant | | | Emx2 mutant | | | Nr2f1 mutant | | | | | |
| | | Emx2 HB | Nr2f1 | Pax6 Hb | Pax6 P | Pbx1 | Emx2 | Lhx2 | Nr2f1 | Pax6 | Pbx1 | 27Ac | 27me3 | ATAC | 27Ac | 27me3 | ATAC | 27Ac | 27me3 | ATAC | | | |
| Nfix | chr8:87060000-87430000 | | | | | | | | | | | | | | | | | | | | | | |
| 1 | chr8:87183906-87187000 | x | | | | | | | x | | | Loss | | Same | Loss | | Same | Loss | | Same | x | | |
| 2 | chr8:87210441-87212579 | | | | | | | | | | | Loss | | Same | Same | | Same | Loss | | Same | x | | Nacc1 TSS |
| 3 | chr8:87225214-87232338 | | | | | | | | x | | | | Same | | | Gain | | | Same | | x | x | Nfix 5'UTR |
| 4 | chr8:87295475-87304533 | x | | | | x | | x | x | x | | | Same | | Loss | Gain | | | Loss | | | x | Nfix TSS |
| 5 | chr8:87306480-87314255 | | | | | | | | x | | | | Same | | | Gain | | | Same | | | x | |
| 6 | chr8:87319898-87327594 | | | | | | | | | | | | | Same | | Gain | Same | | | Same | | x | Nfix alternative TSS |

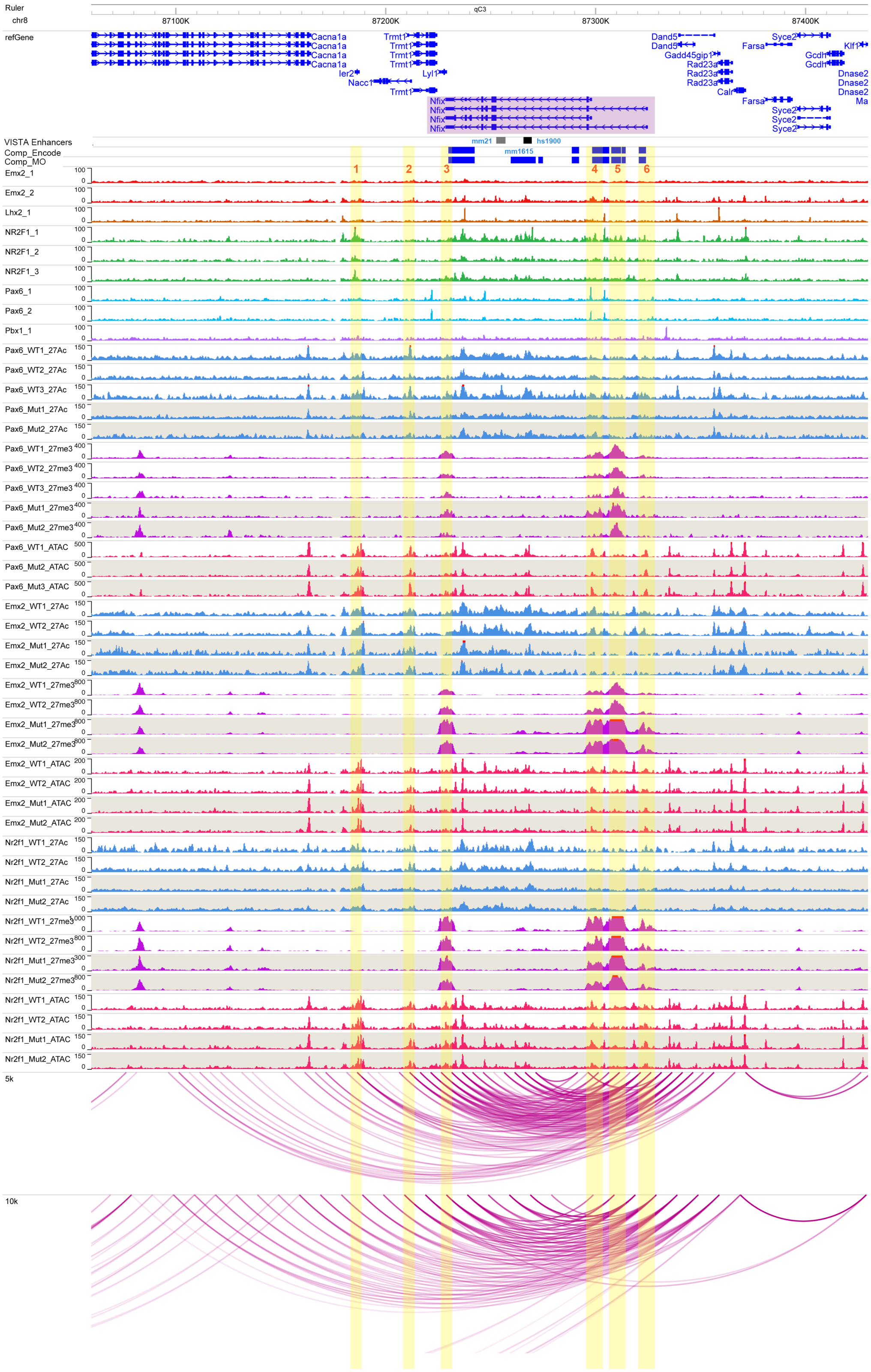

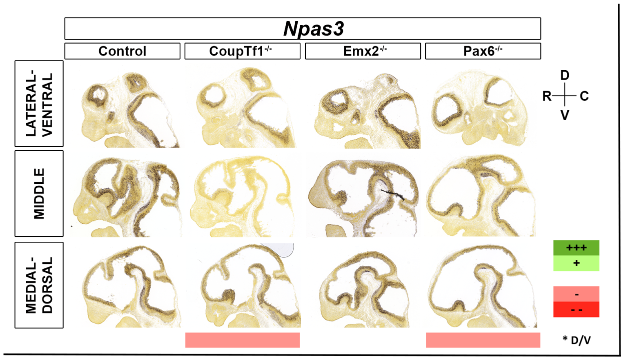


| | | Motifs | | | | | TF ChIP Seq | | | | | Differential epigenomic marks in WT vs. Mutant | | | | | | | | | Plac-Seq | Computation | Genomic features |
|---|---|---|---|---|---|---|---|---|---|---|---|---|---|---|---|---|---|---|---|---|---|---|---|
| | | | | | | | | | | | | Pax6 mutant | | | Emx2 mutant | | | Nr2f1 mutant | | | | | |
| | | Emx2 HB | Nr2f1 | Pax6 Hb | Pax6 P | Pbx1 | Emx2 | Lhx2 | Nr2f1 | Pax6 | Pbx1 | 27Ac | 27me3 | ATAC | 27Ac | 27me3 | ATAC | 27Ac | 27me3 | ATAC | | | |
| Npas3 | chr12:53900000-55700000 | | | | | | | | | | | | | | | | | | | | | | |
| 1 | chr12:54162201-54164479 | x | x | | | | x | x | | | x | Loss | | Same | Loss | | Same | Loss | | Same | x | x | mJR97 |
| 2 | chr12:54212422-54214798 | x | | | | | x | x | | | x | | | Same | | | Loss | | | Loss | x | | |
| 3 | chr12:54224810-54226545 | x | | | | | x | x | | x | x | | | Same | | | Loss | | | Loss | | | |
| 4 | chr12:54347665-54350320 | | | | | | | | | | | Loss | | Same | Same | | Same | Loss | | Same | | x | Npas3 TSS |
| 5 | chr12:54353494-54355559 | x | | | | | x | | | x | x | Loss | | Same | Same | | Same | Loss | | Same | | x | Npas3 Gene Body |
| 6 | chr12:54391045-54394848 | | | | | | x | | x | | x | Loss | | Same | Same | | Same | Loss | | | | x | Npas3 Gene Body |
| 7 | chr12:54548080-54550100 | | | | | | x | x | x | x | x | | | Loss | | Small gain | Same | | | Same | | x | mJR94 |
| 8 | chr12:54593760-54594932 | x | | | | | x | x | x | | x | | | Same | | | Loss | | | Same | x | | Npas3 Gene Body |
| 9 | chr12:54723872-54727081 | x | | x | x | x | x | x | x | x | | Loss | | Loss | Same | | Same | Same? | | Same | x | x | hs527 |

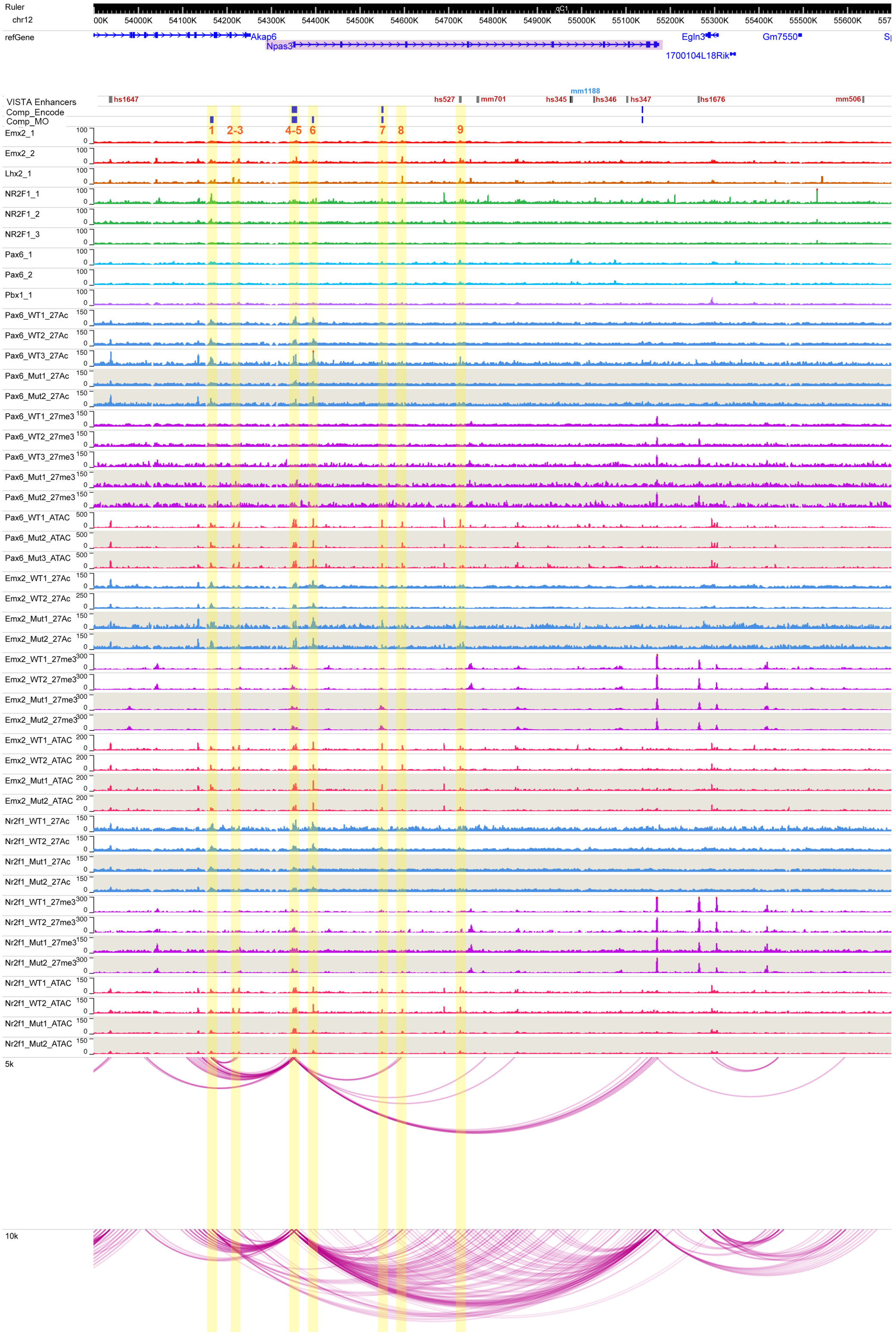

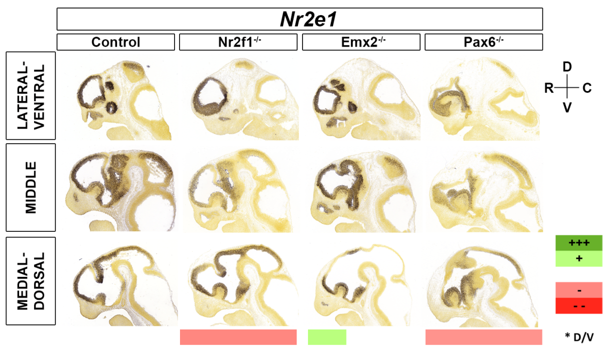


| | | Motifs | | | | | TF ChIP Seq | | | | | Differential epigenomic marks in WT vs. Mutant | | | | | | | | | Plac-Seq | Computation | Genomic features |
|---|---|---|---|---|---|---|---|---|---|---|---|---|---|---|---|---|---|---|---|---|---|---|---|
| | | | | | | | | | | | | Pax6 mutant | | | Emx2 mutant | | | Nr2f1 mutant | | | | | |
| | | Emx2 HB | Nr2f1 | Pax6 Hb | Pax6 P | Pbx1 | Emx2 | Lhx2 | Nr2f1 | Pax6 | Pbx1 | 27Ac | 27me3 | ATAC | 27Ac | 27me3 | ATAC | 27Ac | 27me3 | ATAC | | | |
| Nr2E1 | chr10:41950000-42460000 | | | | | | | | | | | | | | | | | | | | | | |
| 1 | chr10:42297819-42300842 | | | | | | x | | x | x | | Loss | | Loss | Same | | Same | Same | | Same | x | x | Nr2E1 Intron |
| 2 | chr10:42302763-42304738 | x | x | x | x | | x | x | x | x | x | Loss | | Same | Same | | Same | Loss | | Same | x | x | Nr2E1 TSS |
| 3 | chr10:42310968-42312329 | x | | | | | x | | x | x | | Loss | | Same | Same | | Same | Loss | | Same | | x | |
| 4 | chr10:42350739-42352822 | x | | x | x | | x | x | x | x | | Same | | Same | Same | | Same | Loss | | Same | | x | |

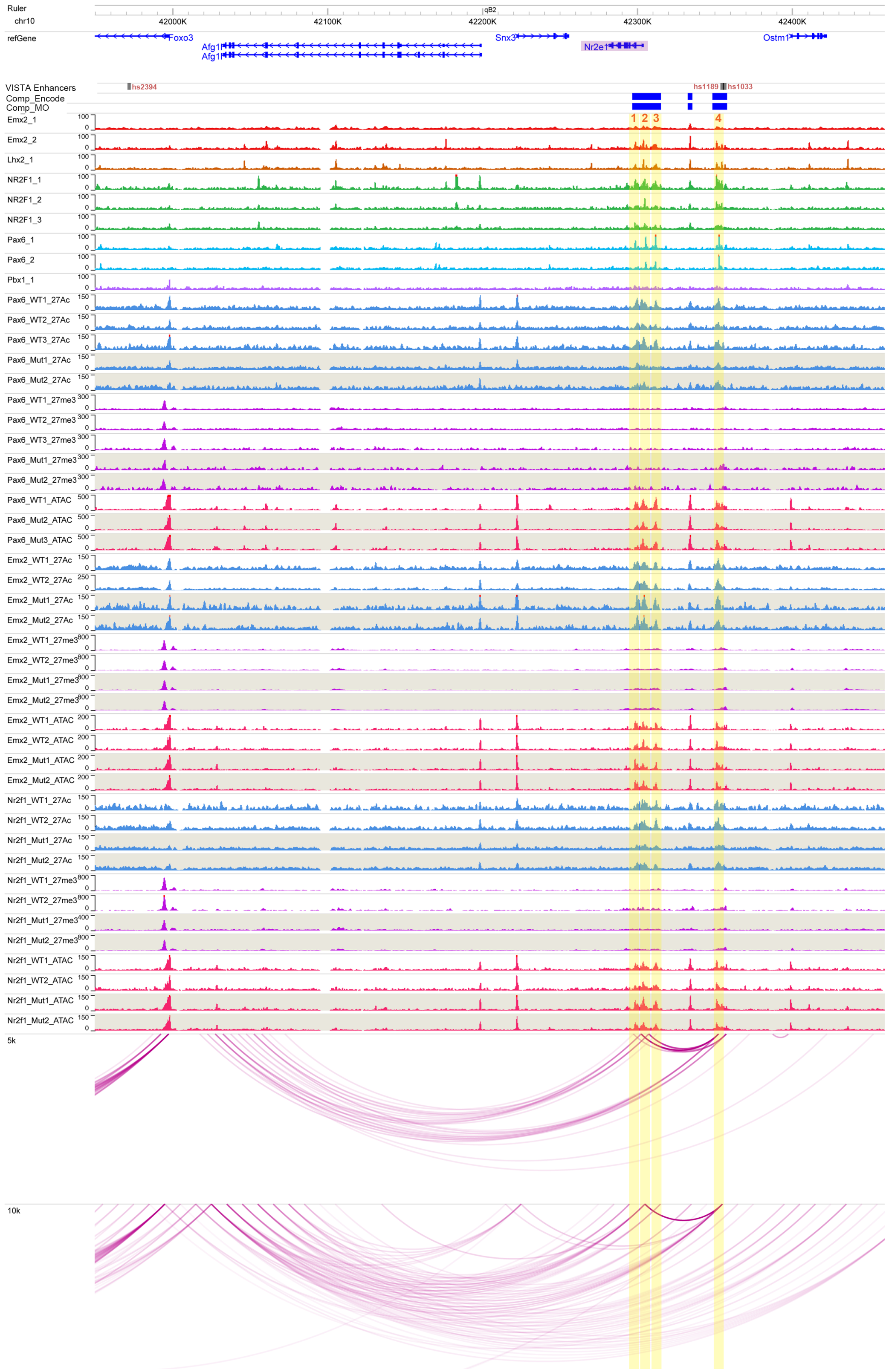

## *Nr2f1*

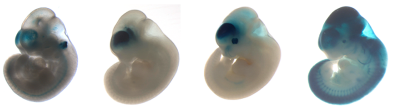


| Nr2f1 | chr13:77500000-79323000 | Motifs: Emx2 HB | Motifs: Nr2f1 | Motifs: Pax6 Hb | Motifs: Pax6 P | Motifs: Pbx1 | TF ChIP Seq: Emx2 | TF ChIP Seq: Lhx2 | TF ChIP Seq: Nr2f1 | TF ChIP Seq: Pax6 | TF ChIP Seq: Pbx1 | Differential epigenomic marks in WT vs. Mutant — Pax6 mutant: 27Ac | Pax6 mutant: 27me3 | Pax6 mutant: ATAC | Emx2 mutant: 27Ac | Emx2 mutant: 27me3 | Emx2 mutant: ATAC | Nr2f1 mutant: 27Ac | Nr2f1 mutant: 27me3 | Nr2f1 mutant: ATAC | Plac-Seq | Computation | Genomic features |
|---|---|---|---|---|---|---|---|---|---|---|---|---|---|---|---|---|---|---|---|---|---|---|---|
| 1 | chr13:77535859-77537676 | x | | | | | x | x | x | | | Same | | Same | Loss | | Loss | | | Same | x | | hs273 |
| 2 | chr13:78031585-78032706 | x | | | | | x | x | x | x | x | Loss | | Same | Loss | Gain | Same | Same | | Same | x | | hs271 |
| 3 | chr13:78033028-78033775 | x | | | | | x | x | x | x | | Loss | | Same | Loss | Gain | Loss | Loss | | Same | x | | |
| 4 | chr13:78238631-78240287 | x | | | | | x | x | | x | | Loss | | Loss | Loss | | Loss | Same | | Same | | x | |
| 5 | chr13:78301285-78303743 | x | | | | | x | | x | | | Loss | | Same | Same | Same | Same | Same | Same | Same | | x | |
| 6 | chr13:78318156-78320079 | x | | | | | x | | x | x | | Loss | Gain | Same | Loss | Gain | Same | Loss | Loss | Same | | x | |
| 7 | chr13:78320294-78321148 | x | | | | | x | | | | | Same | Gain | Same | Loss | Gain | Same | Same | Loss | Same | | x | |
| 8 | chr13:78333521-78334986 | | | | | | | | | | | Loss | Same | Same | Loss | Gain | Same | Loss | Loss | Same | | x | Nr2f1 Gene Body |
| 9 | chr13:78336209-78338805 | x | | | | | x | | | x | | Loss | Same | Same | Loss | Gain | Same | Loss | Loss | Loss | | x | Nr2f1 TSS |
| 10 | chr13:78339478-78340485 | x | | | | | x | x | x | | | | | Same | | Gain | Loss | | Loss | Same | | | |
| 11 | chr13:78341891-78344303 | | | | | | | | x | | | Same | | Same | Loss | Gain | Loss | Same | Loss | Same | | x | |
| 12 | chr13:78348092-78349649 | x | | | | | x | x | | | | Loss | Same | Loss | Loss | Gain | Same | | Loss | Same | | x | |
| 13 | chr13:78565590-78568155 | x | | | | | x | x | | x | | Loss | | Same | Loss | Gain | Same | Same | | Same | x | x | |
| 14 | chr13:78583826-78587247 | | | | | | x | | x | x | | Loss | | Loss | Loss | Gain | Same | Same | | Same | x | | hs1550 |
| 15 | chr13:78719884-78721593 | x | | x | x | x | x | x | x | x | x | | | Same | Loss | Gain | Same | | | Same | x | | hs1172 |
| 16 | chr13:78781101-78783086 | x | | | | | x | x | | | x | Loss | | Loss | Loss | | Loss | Same | | Same | x | | |

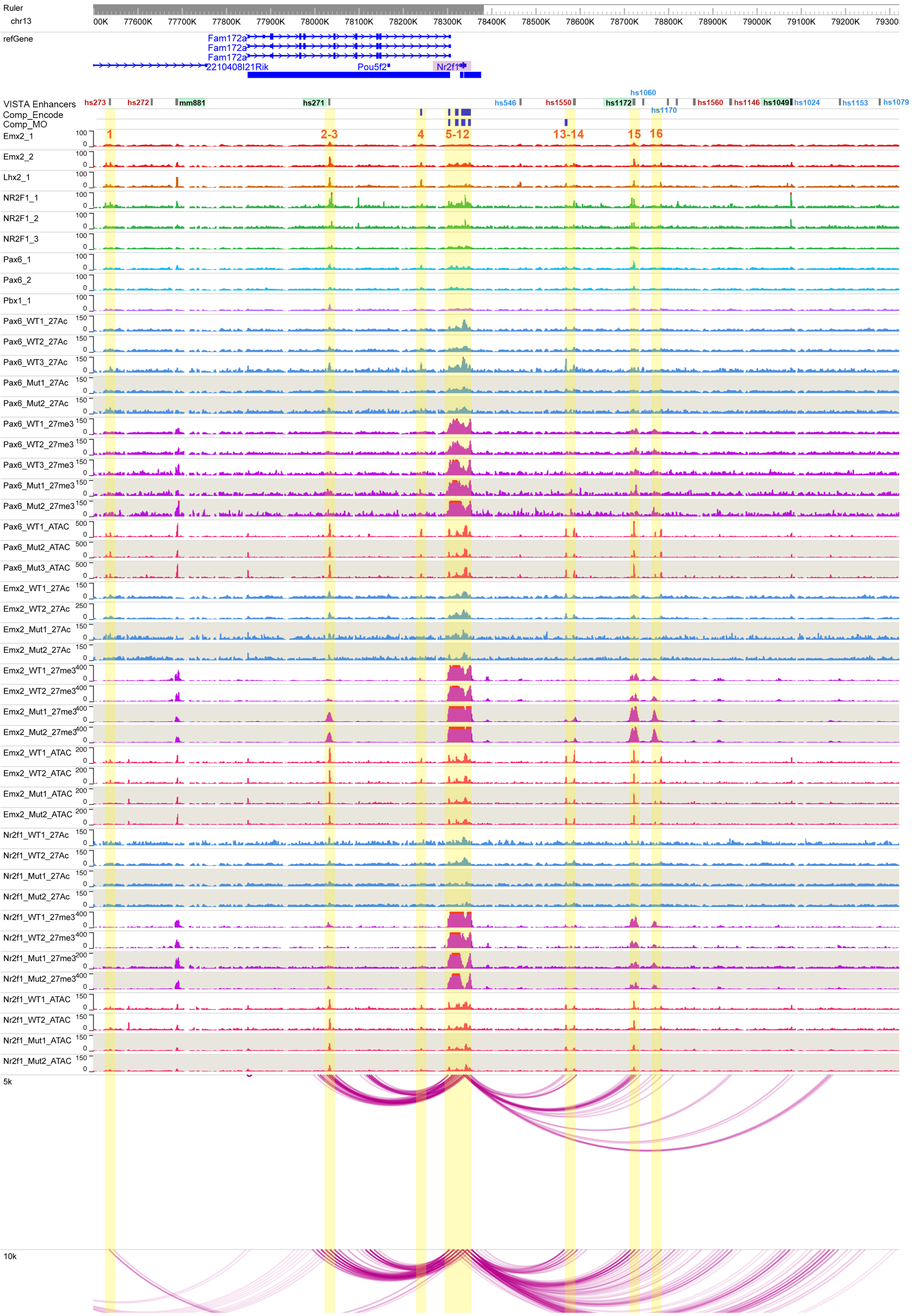

# *Nr2f2*

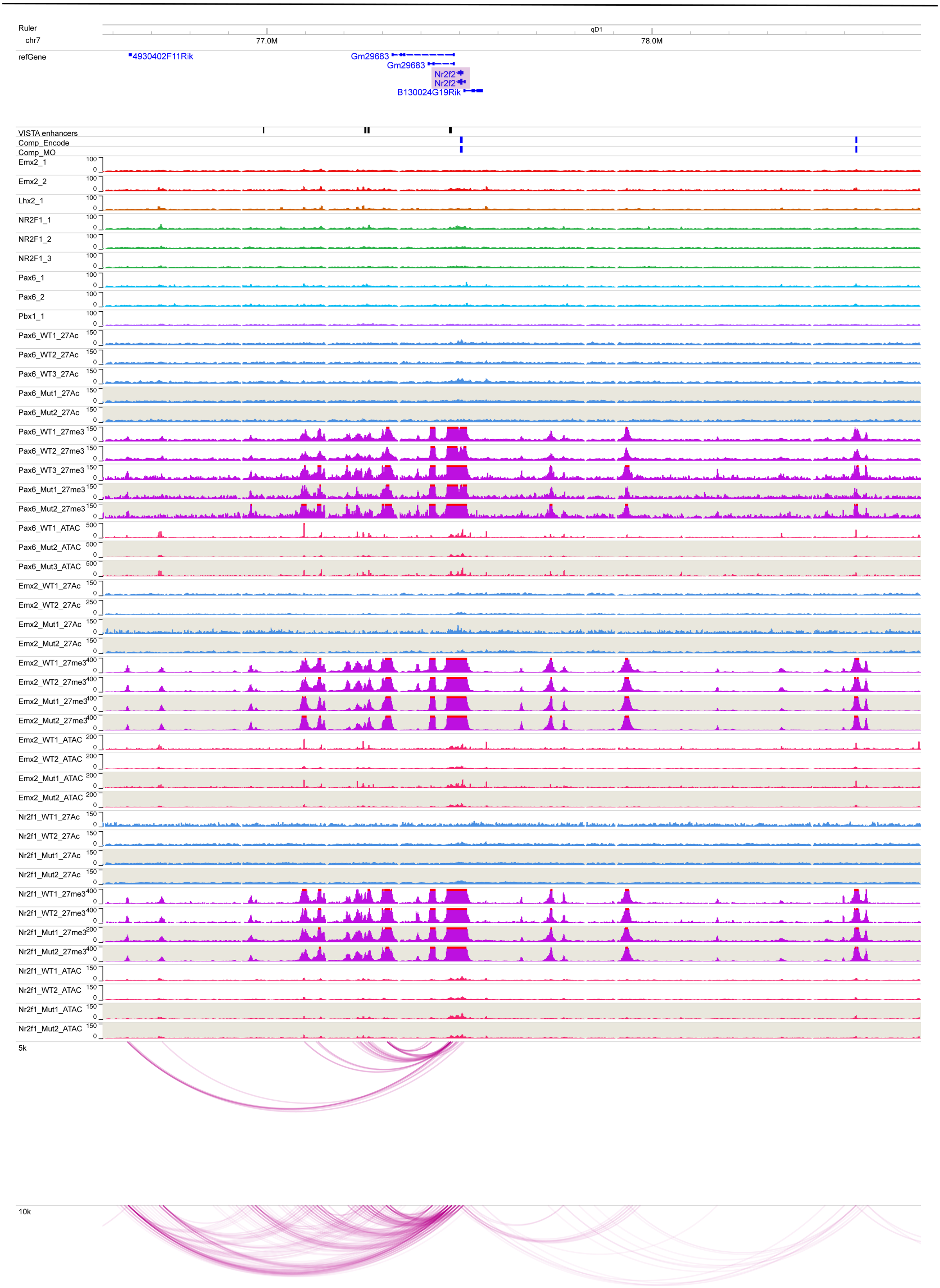
Ruler
chr7
77.0M
78.0M
qD1
refGene
4930402F11Rik
Gm29683
Gm29683
Nr2f2
Nr2f2
B130024G19Rik
VISTA enhancers
Comp_Encode
Comp_MO
Emx2_1
Emx2_2
Lhx2_1
NR2F1_1
NR2F1_2
NR2F1_3
Pax6_1
Pax6_2
Pbx1_1
Pax6_WT1_27Ac
Pax6_WT2_27Ac
Pax6_WT3_27Ac
Pax6_Mut1_27Ac
Pax6_Mut2_27Ac
Pax6_WT1_27me3
Pax6_WT2_27me3
Pax6_WT3_27me3
Pax6_Mut1_27me3
Pax6_Mut2_27me3
Pax6_WT1_ATAC
Pax6_Mut2_ATAC
Pax6_Mut3_ATAC
Emx2_WT1_27Ac
Emx2_WT2_27Ac
Emx2_Mut1_27Ac
Emx2_Mut2_27Ac
Emx2_WT1_27me3
Emx2_WT2_27me3
Emx2_Mut1_27me3
Emx2_Mut2_27me3
Emx2_WT1_ATAC
Emx2_WT2_ATAC
Emx2_Mut1_ATAC
Emx2_Mut2_ATAC
Nr2f1_WT1_27Ac
Nr2f1_WT2_27Ac
Nr2f1_Mut1_27Ac
Nr2f1_Mut2_27Ac
Nr2f1_WT1_27me3
Nr2f1_WT2_27me3
Nr2f1_Mut1_27me3
Nr2f1_Mut2_27me3
Nr2f1_WT1_ATAC
Nr2f1_WT2_ATAC
Nr2f1_Mut1_ATAC
Nr2f1_Mut2_ATAC
5k
10k

### *Pax6*

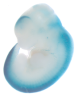
hs1531

| | | Motifs | | | | | TF ChIP Seq | | | | | Differential epigenomic marks in WT vs. Mutant | | | | | | | | | Plac-Seq | Computation | Genomic features |
|---|---|---|---|---|---|---|---|---|---|---|---|---|---|---|---|---|---|---|---|---|---|---|---|
| | | | | | | | | | | | | Pax6 mutant | | | Emx2 mutant | | | Nr2f1 mutant | | | | | |
| | | Emx2 HB | Nr2f1 | Pax6 Hb | Pax6 P | Pbx1 | Emx2 | Lhx2 | Nr2f1 | Pax6 | Pbx1 | 27Ac | 27me3 | ATAC | 27Ac | 27me3 | ATAC | 27Ac | 27me3 | ATAC | | | |
| Pax6 | chr2:105085000-106050000 | | | | | | | | | | | | | | | | | | | | | | |
| 1 | chr2:105355979-105356970 | | | | | | x | x | x | x | x | Loss | | Same | Same | | Same | Same | | Same | | | hs855 |
| 2 | chr2:105451245-105454219 | | | | | | x | x | | | | Same | | Same | Same | | Same | Loss | | Same | | | hs1531 |
| 3 | chr2:105502815-105507481 | | | | | | | | | | | | Loss | | | Same | | | Same | | | | Pax6 TSS |
| 4 | chr2:105558877-105560976 | | | | | | x | x | x | x | x | Loss | | Same | Loss | | Same | Same | | Same | | | hs198 |

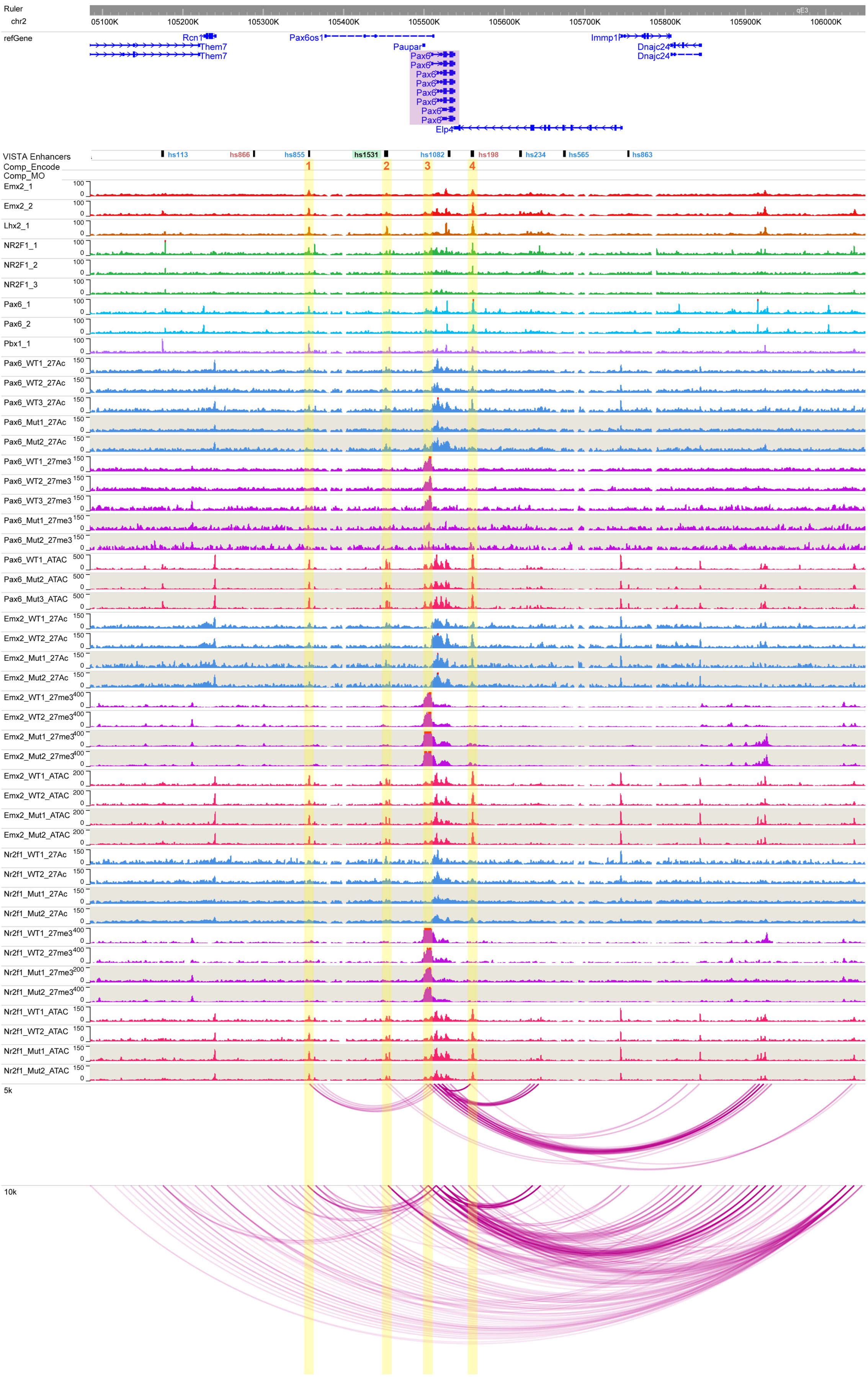

### *Pbx1*

| | | Motifs | | | | | TF ChIP Seq | | | | | Differential epigenomic marks in WT vs. Mutant | | | | | | | | | Plac-Seq | Computation | Genomic features |
|---|---|---|---|---|---|---|---|---|---|---|---|---|---|---|---|---|---|---|---|---|---|---|---|
| | | | | | | | | | | | | Pax6 mutant | | | Emx2 mutant | | | Nr2f1 mutant | | | | | |
| | | Emx2 HB | Nr2f1 | Pax6 Hb | Pax6 P | Pbx1 | Emx2 | Lhx2 | Nr2f1 | Pax6 | Pbx1 | 27Ac | 27me3 | ATAC | 27Ac | 27me3 | ATAC | 27Ac | 27me3 | ATAC | | | |
| **Pbx1** | **chr1:169860000-171370500** | | | | | | | | | | | | | | | | | | | | | | |
| **1** | chr1:169975190-169978135 | | | | | | x | x | x | x | x | Same | | Loss | Loss | | Loss | Same | | Same | x | | |
| **2** | chr1:170175012-170176937 | | | | | | x | | x | | x | Loss | | Same | Loss | | Same | Loss | | Same | x | | Pbx1 Gene Body |
| **3** | chr1:170229514-170231269 | | | | | | | | | | | Loss | | | Loss | | | Loss | | | x | | hs1144 |
| **4** | chr1:170257128-170258791 | | | | | | x | x | | | | Same | | Same | Same | | Loss | Same | | Same | x | | hs203 |
| **5** | chr1:170321692-170324410 | | | | | | x | x | x | x | | Same | | Loss | Loss | | Loss | Loss | | | x | x | Pbx1 Gene Body |
| **6** | chr1:170340225-170341639 | | | | | | x | | | | | | | Loss | | | Same | | | Same | | x | Pbx1 Gene Body |
| **7** | chr1:170360324-170363111 | | | | | | | | x | | x | Loss | | Same | Same | | Same | Same | | Same | | x | Pbx1 TSS |
| **8** | chr1:170837816-170839853 | | | | | | x | x | x | | x | Loss | | Same | Loss | | Loss | Same | | Same | x | | hs202 |

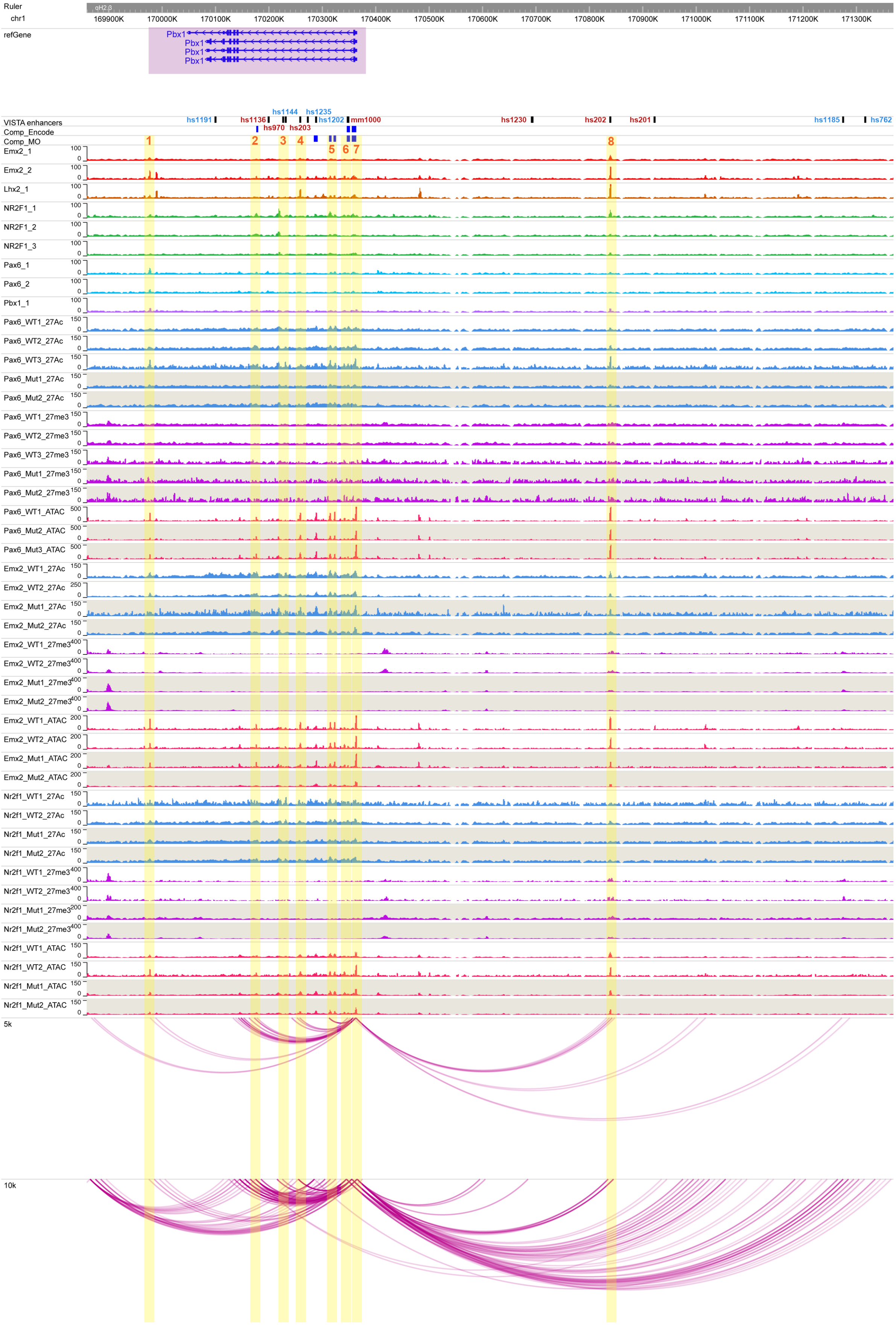

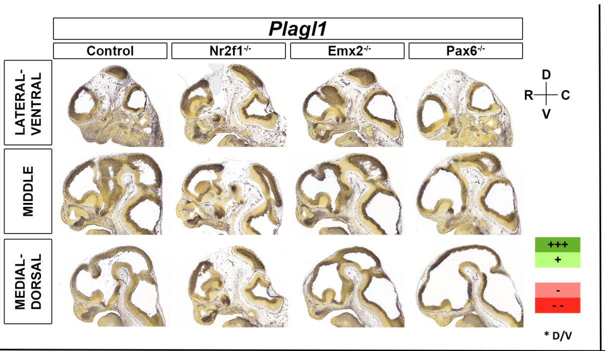


| | | Motifs | | | | | TF ChIP Seq | | | | | Differential epigenomic marks in WT vs. Mutant | | | | | | | | | Plac-Seq | Computation | Genomic features |
|---|---|---|---|---|---|---|---|---|---|---|---|---|---|---|---|---|---|---|---|---|---|---|---|
| | | | | | | | | | | | | Pax6 mutant | | | Emx2 mutant | | | Nr2f1 mutant | | | | | |
| | | Emx2 HB | Nr2f1 | Pax6 Hb | Pax6 P | Pbx1 | Emx2 | Lhx2 | Nr2f1 | Pax6 | Pbx1 | 27Ac | 27me3 | ATAC | 27Ac | 27me3 | ATAC | 27Ac | 27me3 | ATAC | | | |
| **Plagl1** | chr10:12700000-13310000 | | | | | | | | | | | | | | | | | | | | | | |
| **1** | chr10:12809880-12812996 | | | | | | | | | | | Loss | | Same | Loss | | Same | Same | | Same | | | Plagl1 TSS |
| **2** | chr10:12826983-12831335 | x | | | | | x | x | x | x | x | Loss | | Same | Loss | | Same | Same | | Same | | x | Plagl1 Gene body |
| **3** | chr10:12836410-12839285 | x | x | x | x | | x | x | x | x | | Loss | | Same | Loss | | Same | Same | | Same | | x | Plagl1 Gene body |
| **4** | chr10:13043102-13045156 | x | x | | | | | | x | | | Loss | | Same | Same | | Same | Loss | | Same | x | | Phactr2 TSS |
| **5** | chr10:13135354-13136688 | x | | | | | x | x | x | | | Loss | | Same | Loss | | Same | Same | | Same | x | | |

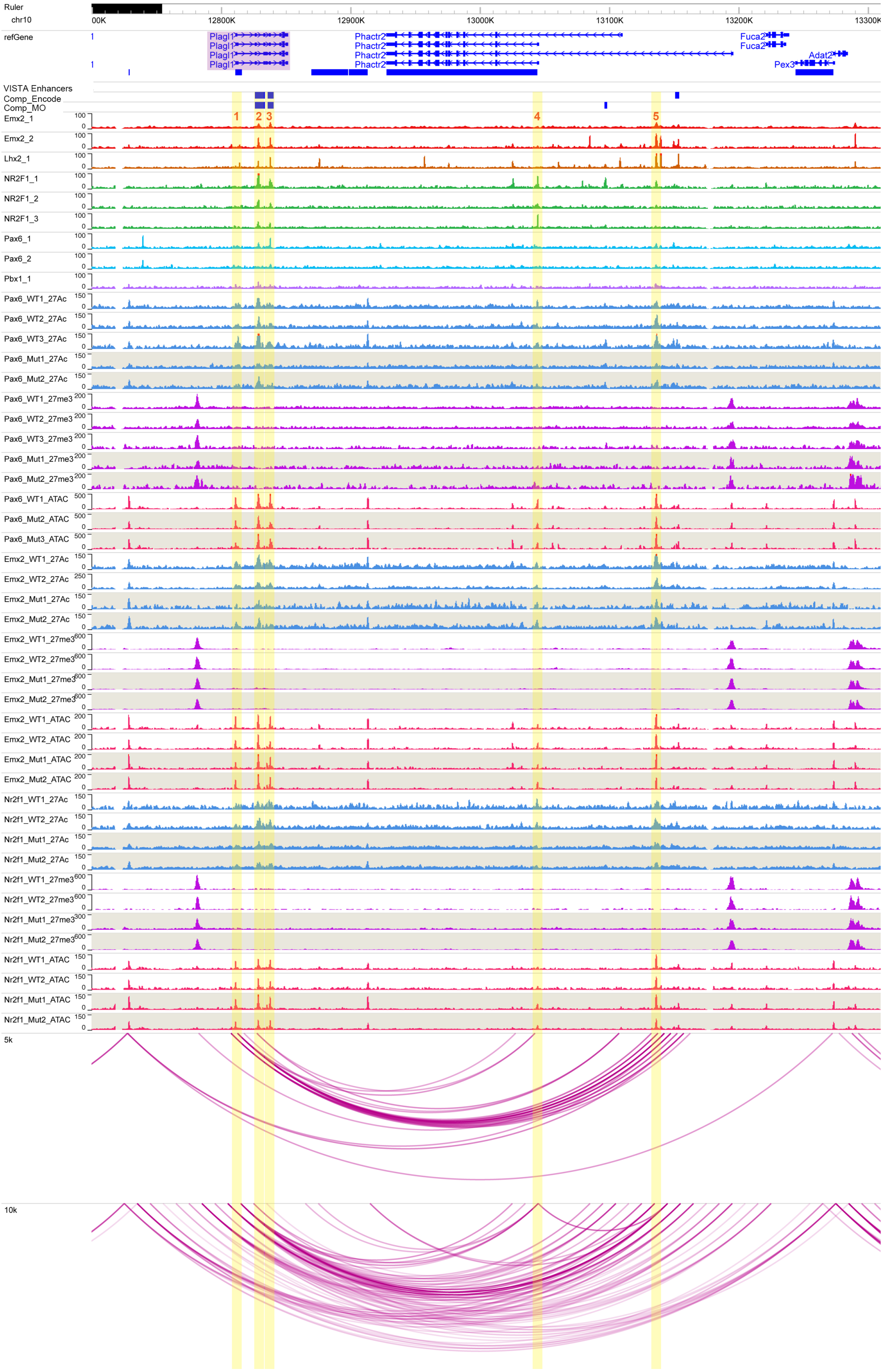

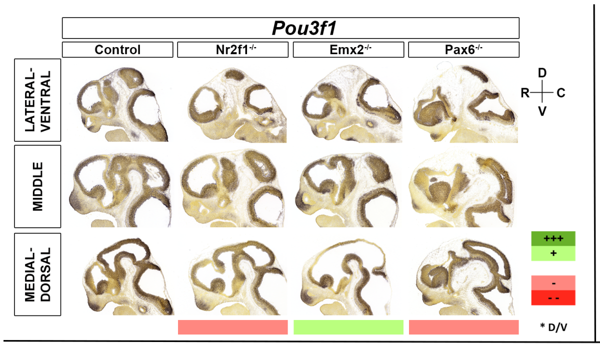


| | | Motifs | | | | | TF ChIP Seq | | | | | Differential epigenomic marks in WT vs. Mutant | | | | | | | | | Plac-Seq | Computation | Genomic features |
|---|---|---|---|---|---|---|---|---|---|---|---|---|---|---|---|---|---|---|---|---|---|---|---|
| | | | | | | | | | | | | Pax6 mutant | | | Emx2 mutant | | | Nr2f1 mutant | | | | | |
| | | Emx2 HB | Nr2f1 | Pax6 Hb | Pax6 P | Pbx1 | Emx2 | Lhx2 | Nr2f1 | Pax6 | Pbx1 | 27Ac | 27me3 | ATAC | 27Ac | 27me3 | ATAC | 27Ac | 27me3 | ATAC | | | |
| Pou3f1 | chr4:123550000-124550000 | | | | | | | | | | | | | | | | | | | | | | |
| 1 | chr4:123711996-123714726 | x | | x | x | | x | | x | | | Loss | | Same | Loss | | Same | Same | | Same | | x | hs1031 |
| 2 | chr4:124072428-124078145 | x | | x | x | x | x | x | x | | x | | Same | Same | | Loss | Loss | | Same | Same | x | | hs270 |
| 3 | chr4:124118120-124122688 | | | x | x | | | | | x | | | Same | | | Loss | | | Same | | x | | |
| 4 | chr4:124132233-124133640 | | | | | | | | | | | | Same | | | Loss | | | Same | | x | | hs1008 |
| 5 | chr4:124135319-124138674 | | | | | | | | | | | | Same | | | Loss | | | Same | | x | | hs2088 |
| 6 | chr4:124142356-124147119 | x | | | | | x | x | x | x | x | | Same | Same | | Loss | Loss | | Same | | x | | |
| 7 | chr4:124154196-124157660 | | | | | | | | x | | | | Same | | | Loss | | | Same | | x | | hs1001 |
| 8 | chr4:124335186-124337892 | | | | | | | | | | | | Same | | | Loss | | | Loss | | | | Pou3f1 gene body |

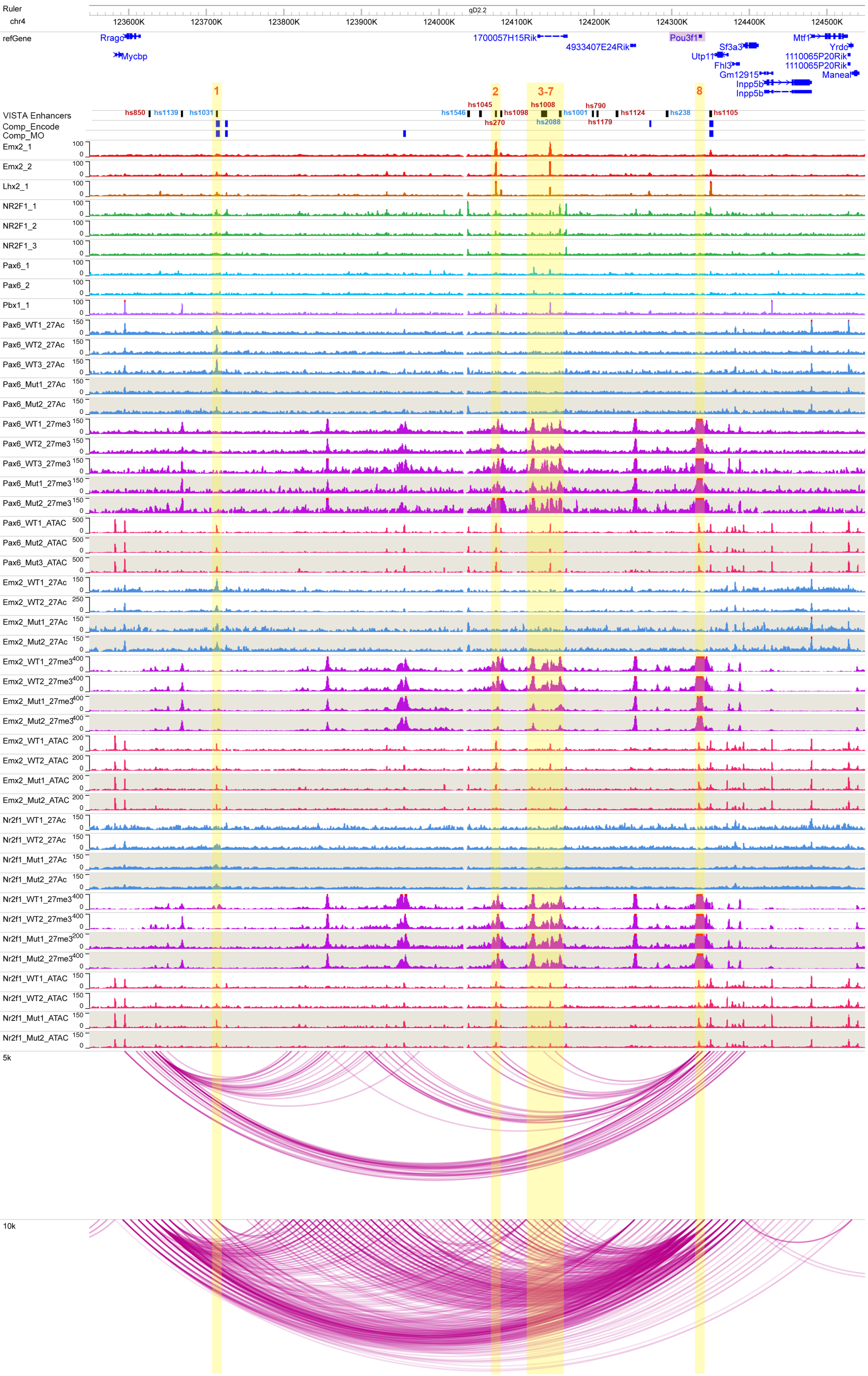

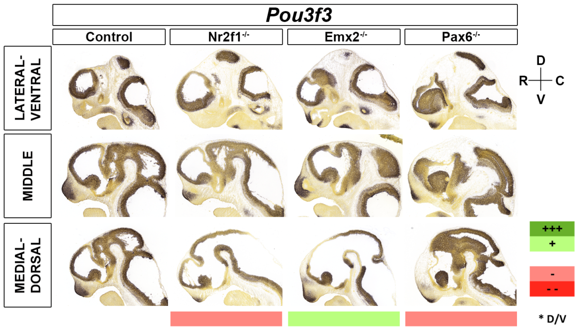


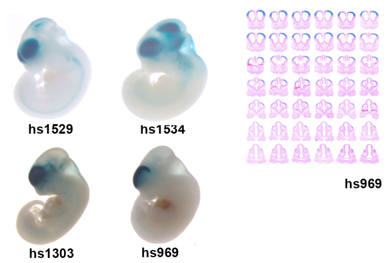


| | | Motifs | | | | | TF ChIP Seq | | | | | Differential epigenomic marks in WT vs. Mutant | | | | | | | | | Plac-Seq | Computation | Genomic features |
|---|---|---|---|---|---|---|---|---|---|---|---|---|---|---|---|---|---|---|---|---|---|---|---|
| | | | | | | | | | | | | Pax6 mutant | | | Emx2 mutant | | | Nr2f1 mutant | | | | | |
| | | Emx2 HB | Nr2f1 | Pax6 Hb | Pax6 P | Pbx1 | Emx2 | Lhx2 | Nr2f1 | Pax6 | Pbx1 | 27Ac | 27me3 | ATAC | 27Ac | 27me3 | ATAC | 27Ac | 27me3 | ATAC | | | |
| Pou3f3 | chr1:41700000-42857055 | | | | | | | | | | | | | | | | | | | | | | |
| 1 | chr1:41798138-41801834 | x | | x | x | | x | x | x | x | x | Same | | Loss | Same | | Same | Same | | Same | x | | |
| 2 | chr1:41870353-41872353 | x | | | | | x | x | x | x | x | Loss | | Same | Same | | Same | Same? | | Same | x | x | hs1529 |
| 3 | chr1:41992227-41995700 | x | | x | x | | x | x | | x | x | | Loss | Loss | | Loss | Loss | | Same | Same | x | | |
| 4 | chr1:42313865-42317133 | x | | x | | x | x | | x | x | x | Loss | | Same | Same | | Same | Loss | | Same | x | x | hs960 |
| 5 | chr1:42595469-42598795 | x | | | | | x | x | x | x | x | Loss | | Same | Same | | Same | Loss | | Same | x | x | hs969 |
| 6 | chr1:42746840-42749611 | | | | | | x | | x | x | | Loss | | Same | Gain | | Same | Loss | | Same | | | |
| 7 | chr1:42753952-42757047 | | | | | | | | x | | | Loss | | | Gain | | | Loss | | | | x | Pou3f3 gene body |
| 8 | chr1:42769552-42771799 | | x | | | | | | x | | x | Loss | | Same | Gain | | Same | Loss | | Same | | | |

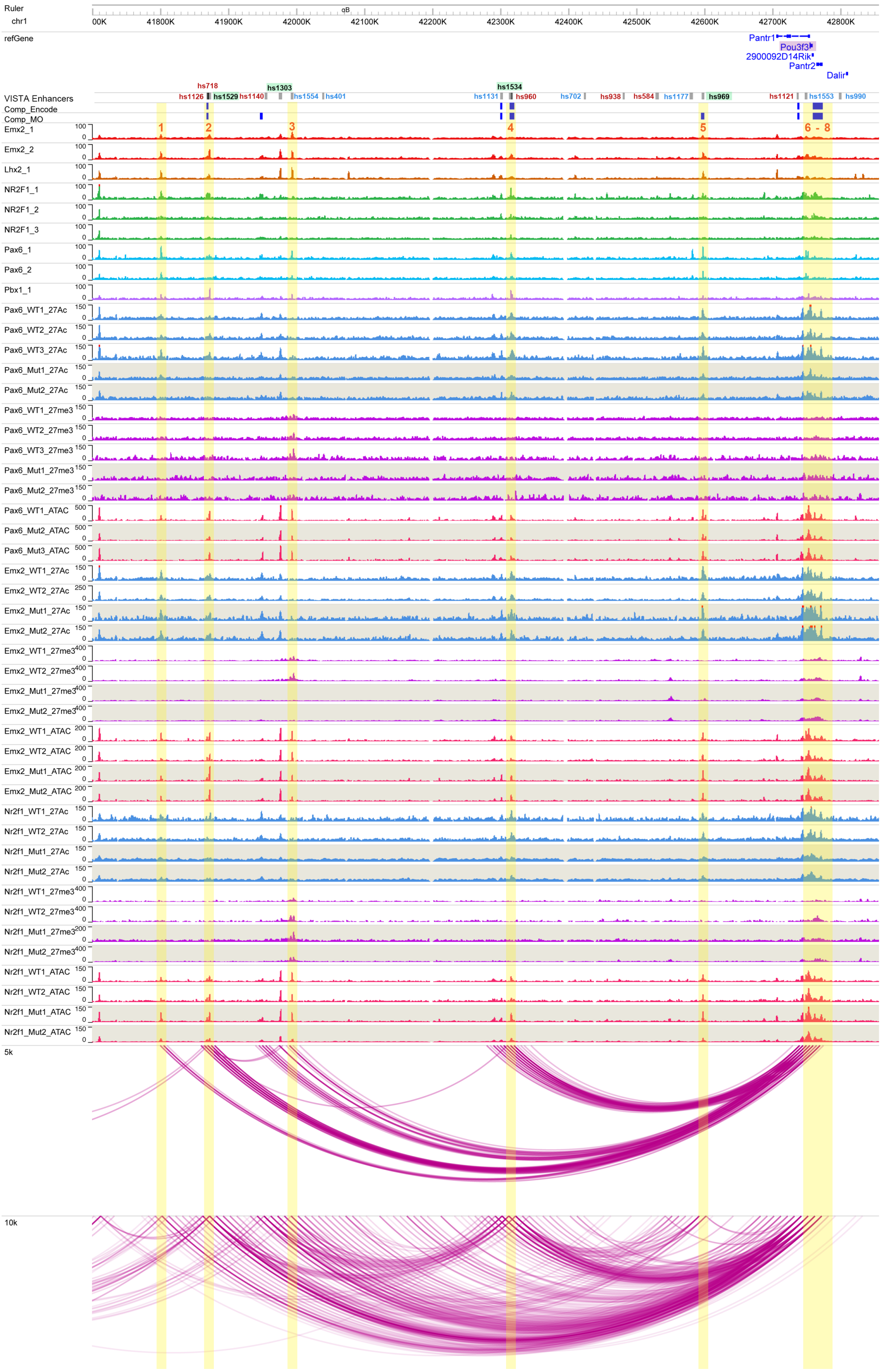

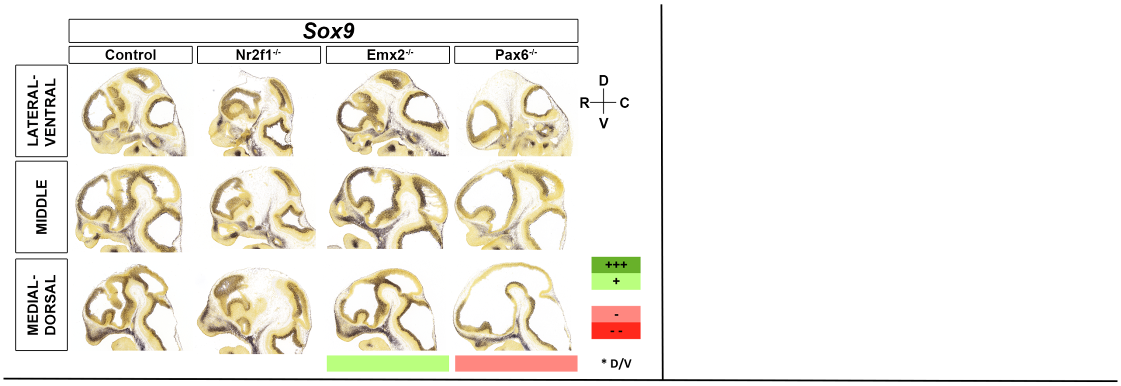


| | | Motifs | | | | | TF ChIP Seq | | | | | Differential epigenomic marks in WT vs. Mutant | | | | | | | | | Plac-Seq | Computation | Genomic features |
|---|---|---|---|---|---|---|---|---|---|---|---|---|---|---|---|---|---|---|---|---|---|---|---|
| | | | | | | | | | | | | Pax6 mutant | | | Emx2 mutant | | | Nr2f1 mutant | | | | | |
| | | Emx2 HB | Nr2f1 | Pax6 Hb | Pax6 P | Pbx1 | Emx2 | Lhx2 | Nr2f1 | Pax6 | Pbx1 | 27Ac | 27me3 | ATAC | 27Ac | 27me3 | ATAC | 27Ac | 27me3 | ATAC | | | |
| **Sox9** | chr11:111700000-113500000 | | | | | | | | | | | | | | | | | | | | | | |
| **1** | chr11:112058352-112060480 | x | | x | x | x | x | x | x | x | x | Loss | | Same | Same | | Same | Loss | | Same | | | |
| **2** | chr11:112273236-112274832 | x | | | | | x | x | x | | | Loss | | Same | Loss | | Loss | Loss | | Same | | | mm634 |
| **3** | chr11:112669970-112674132 | | | | | | x | x | x | | | Loss | | Same | Loss | | Same | Loss | | | | | |
| **4** | chr11:112838032-112839034 | x | | | | | x | x | x | | | Loss | | Same | Same | | Same | Loss | | Same | x | x | |
| **5** | chr11:112843512-112845082 | x | | x | | | x | x | x | x | | Same | | Loss | Same | | Same | Same | | Same | x | | |
| **6** | chr11:112877902-112879155 | | | | | | x | x | x | | | Same | | Same | Loss | | Same | Same | | Same | x | x | mm636 |
| **7** | chr11:113300047-113301927 | | | x | | | x | | x | | | Same | | Same | Loss | | Same | Same | | Same | | x | |

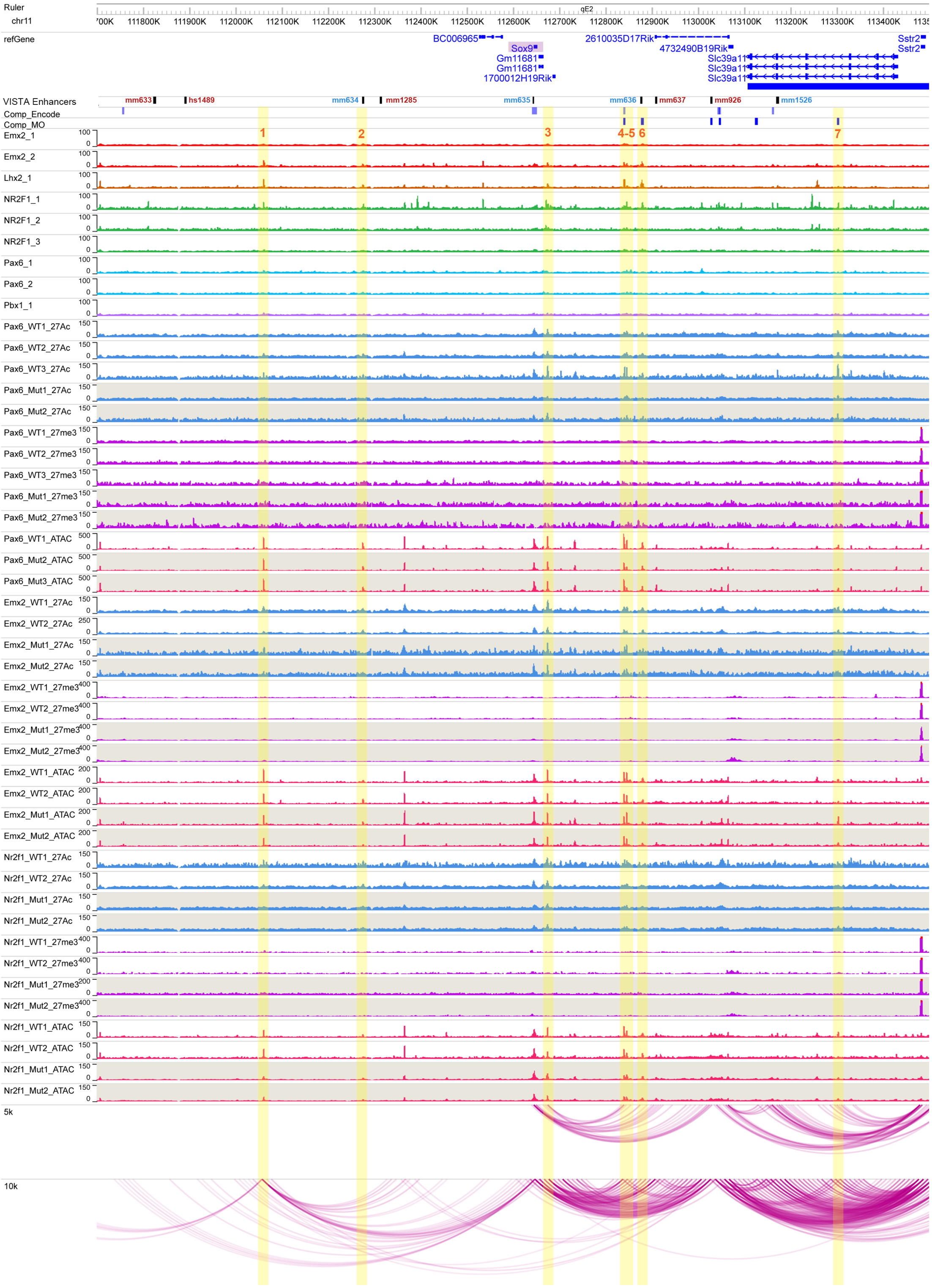

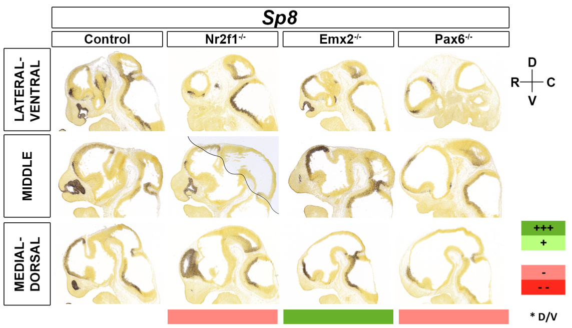


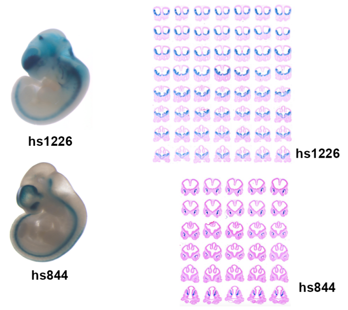


| | | Motifs | | | | | TF ChIP Seq | | | | | Differential epigenomic marks in WT vs. Mutant | | | | | | | | | Plac-Seq | Computation | Genomic features |
|---|---|---|---|---|---|---|---|---|---|---|---|---|---|---|---|---|---|---|---|---|---|---|---|
| | | | | | | | | | | | | Pax6 mutant | | | Emx2 mutant | | | Nr2f1 mutant | | | | | |
| | | Emx2 HB | Nr2f1 | Pax6 Hb | Pax6 P | Pbx1 | Emx2 | Lhx2 | Nr2f1 | Pax6 | Pbx1 | 27Ac | 27me3 | ATAC | 27Ac | 27me3 | ATAC | 27Ac | 27me3 | ATAC | | | |
| Sp8 | chr12:119000000-120191051 | | | | | | | | | | | | | | | | | | | | | | |
| 1 | chr12:119043999-119045761 | x | | x | x | | x | x | | x | x | | Same | Loss | | Same | Same | | Same | Same | x | | hs807 |
| 2 | chr12:119047456-119048848 | x | | | | | x | x | x | | x | | | Same | | | Loss | | | Same | x | | |
| 3 | chr12:119266560-119267560 | x | | | | | | x | | | | | Same | Same | | Loss | | | Same | | x | | |
| 4 | chr12:119639681-119646276 | | | | | | | | | | | | Loss | | | Loss | | | Same | | x | | |
| 5 | chr12:119767849-119769440 | x | | x | x | | x | | x | | | Loss | | Loss | Loss | | Same | Same | | Same | x | | |
| 6 | chr12:119888958-119890368 | | | x | | | x | | x | x | x | | | Loss | | | Gain | | | Same | x | x | hs1226 |
| 7 | chr12:119897620-119901100 | x | | x | | | x | x | x | x | x | | Same | Loss | | Same | Same | | Same | Same | x | | |
| 8 | chr12:119957232-119961000 | x | | | | | x | x | x | | | | Same | Same | | Loss | Same | | Same | Same | | | |
| 9 | chr12:120019929-120024242 | x | | | | | x | x | x | | | | Same | Same | | Loss | | | Same | | | | |
| 10 | chr12:120078064-120080117 | x | | | | | x | x | x | x | | Loss | Gain | Loss | Same | Same | Same | Same | Same | Same | | x | hs844 |
| 11 | chr12:120084740-120088847 | | | | | | | | | | | Loss | Gain | Same | Gain | Same | Same | Loss | Same | Same | | x | Sp8 Gene body |

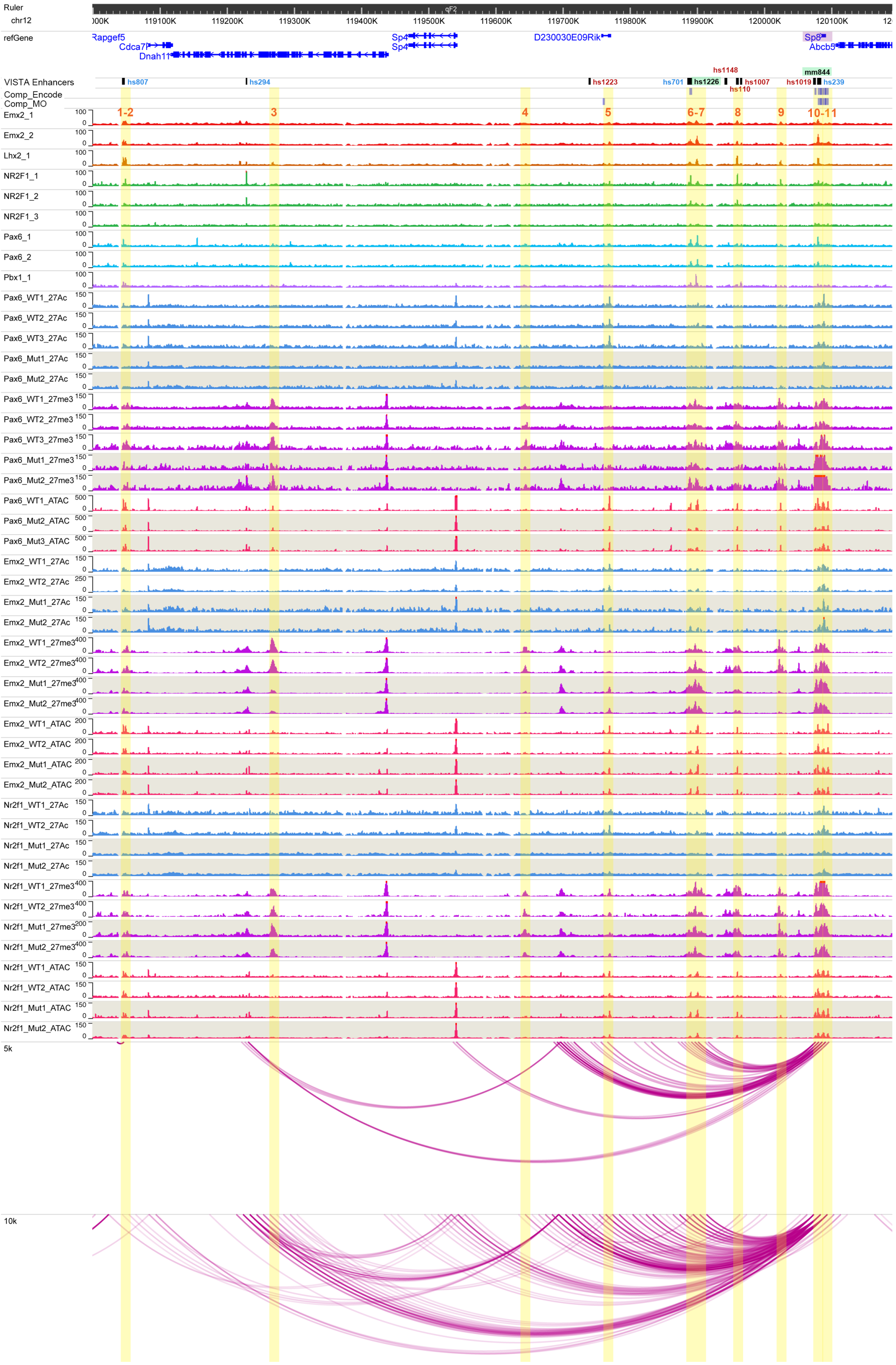

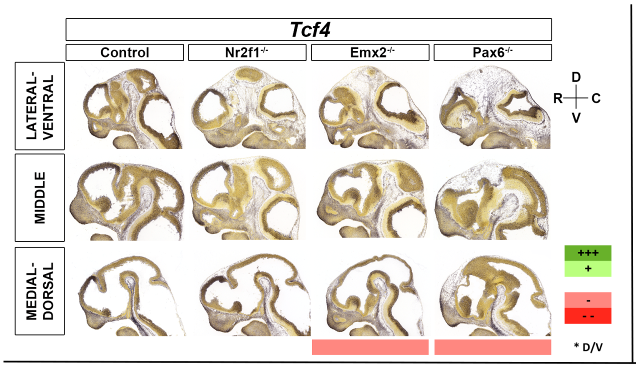


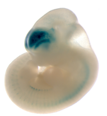


| | | Motifs | | | | | TF ChIP Seq | | | | | Differential epigenomic marks in WT vs. Mutant | | | | | | | | | Plac-Seq | Computation | Genomic features |
|---|---|---|---|---|---|---|---|---|---|---|---|---|---|---|---|---|---|---|---|---|---|---|---|
| | | | | | | | | | | | | Pax6 mutant | | | Emx2 mutant | | | Nr2f1 mutant | | | | | |
| | | Emx2 HB | Nr2f1 | Pax6 Hb | Pax6 P | Pbx1 | Emx2 | Lhx2 | Nr2f1 | Pax6 | Pbx1 | 27Ac | 27me3 | ATAC | 27Ac | 27me3 | ATAC | 27Ac | 27me3 | ATAC | | | |
| Tcf4 | chr18:68600000-69947621 | | | | | | | | | | | | | | | | | | | | | | |
| 1 | chr18:69357344-69358848 | | | | | | | | x | | | | Same | Same | | Gain | Loss | | Same | Same | x | x | |
| 2 | chr18:69503409-69507465 | | | | | | x | | | | | Loss | | Same | Loss | | Same | Loss | | Same | | x | TCF4 TSS |
| 3 | chr18:69658711-69661831 | x | | x | x | | x | x | x | x | x | Loss | | Same | Loss | | Loss | Loss | | Same | x | | hs376 |
| 4 | chr18:69676248-69678881 | | | | | | x | | | | x | Same | | Same | Loss | | Loss | Same | | Same | x | x | |
| 5 | chr18:69721590-69724223 | x | | | | x | x | x | x | x | x | Loss | | Same | Loss | | Same | Same? | | Same | x | | hs1537 |
| 6 | chr18:69751317-69754326 | x | | x | x | | x | x | x | x | x | Loss | | Same | Loss | | Same | Same | | Same | x | x | |
| 7 | chr18:69765992-69770130 | x | | x | | x | x | x | x | x | x | Same | | Same | Loss | | Loss | Same | | Same | x | x | hs1561 |

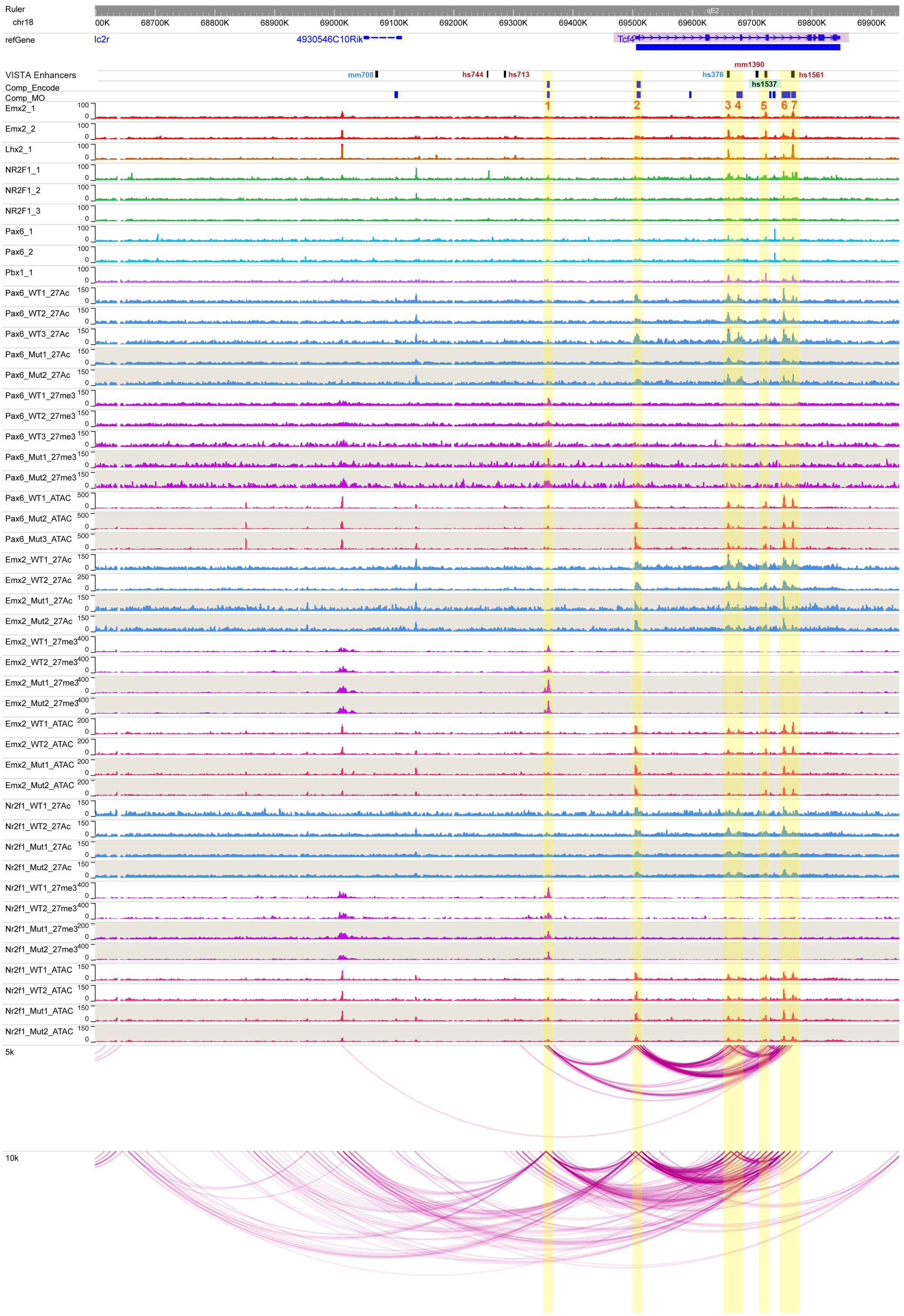

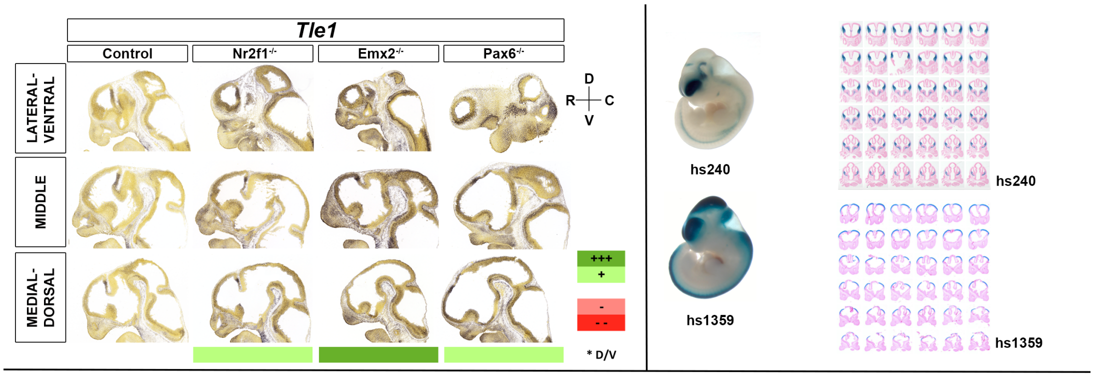


| | | Motifs | | | | | TF ChIP Seq | | | | | Differential epigenomic marks in WT vs. Mutant | | | | | | | | | Plac-Seq | Computation | Genomic features |
|---|---|---|---|---|---|---|---|---|---|---|---|---|---|---|---|---|---|---|---|---|---|---|---|
| | | | | | | | | | | | | Pax6 mutant | | | Emx2 mutant | | | Nr2f1 mutant | | | | | |
| | | Emx2 HB | Nr2f1 | Pax6 Hb | Pax6 P | Pbx1 | Emx2 | Lhx2 | Nr2f1 | Pax6 | Pbx1 | 27Ac | 27me3 | ATAC | 27Ac | 27me3 | ATAC | 27Ac | 27me3 | ATAC | | | |
| Tle1 | chr4:71000000-72800000 | | | | | | | | | | | | | | | | | | | | | | |
| 1 | chr4:71138280-71141616 | | | | | | | | | | | | Loss | | | Loss | | | Same | | x | | |
| 2 | chr4:71145257-71148745 | x | | x | | | x | x | x | x | x | | Same | Same | | Loss | Same | | Same | Same | x | | hs240 |
| 3 | chr4:71594251-71595160 | x | | x | x | | x | x | | | | | | Same | | | Loss | | | Same | x | | |
| 4 | chr4:71710362-71711705 | | | | | | | | | x | | | | Loss | | | Same | | | Same | | | |
| 5 | chr4:71806612-71808583 | | | | | | x | | x | x | | | | Loss | Same | | Same | Same | | Same | | | |
| 6 | chr4:71820673-71822175 | x | | x | x | | x | x | | x | | | | Loss | | | Same | | | Same | | | |
| 7 | chr4:71859657-71862582 | | | | | x | x | | x | | x | Loss | Loss | Same | Gain | Loss | Same | Loss | Same | Same | | x | Tle1 TSS |

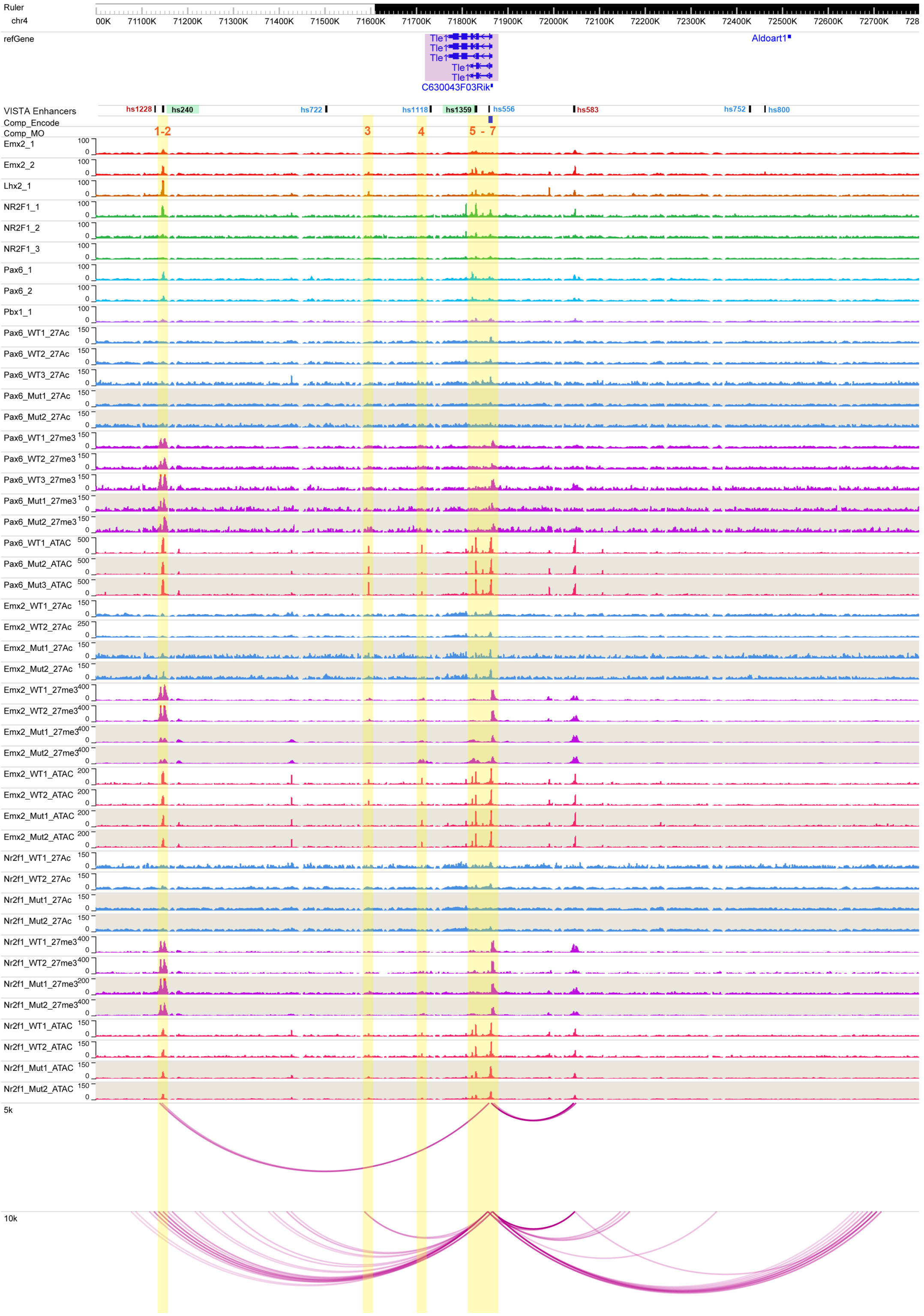

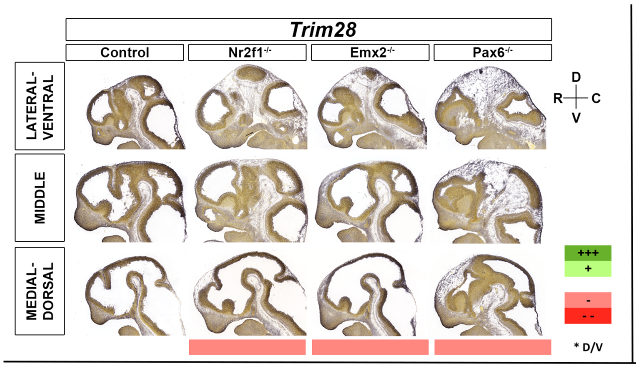


| | | Motifs | | | | | TF ChIP Seq | | | | | Differential epigenomic marks in WT vs. Mutant | | | | | | | | | Plac-Seq | Computation | Genomic features |
|---|---|---|---|---|---|---|---|---|---|---|---|---|---|---|---|---|---|---|---|---|---|---|---|
| | | | | | | | | | | | | Pax6 mutant | | | Emx2 mutant | | | Nr2f1 mutant | | | | | |
| | | Emx2 HB | Nr2f1 | Pax6 Hb | Pax6 P | Pbx1 | Emx2 | Lhx2 | Nr2f1 | Pax6 | Pbx1 | 27Ac | 27me3 | ATAC | 27Ac | 27me3 | ATAC | 27Ac | 27me3 | ATAC | | | |
| Trim28 | chr7:13480000-13716381 | | | | | | | | | | | | | | | | | | | | | | |
| 1 | chr7:13594337-13595650 | | | x | x | x | | | | x | x | Same | | Same | Same | | Same | Loss | | Same | | | Zbtb45 TSS |
| 2 | chr7:13608554-13611167 | | | | | | | | | | | Loss | | Same | Same | | Same | Same | | Same | | | Trim28 TSS |
| 3 | chr7:13622814-13623925 | | | x | | | | | x | | | Loss | | Same | Same | | Same | Loss | | Same | | | Ube2m TSS |

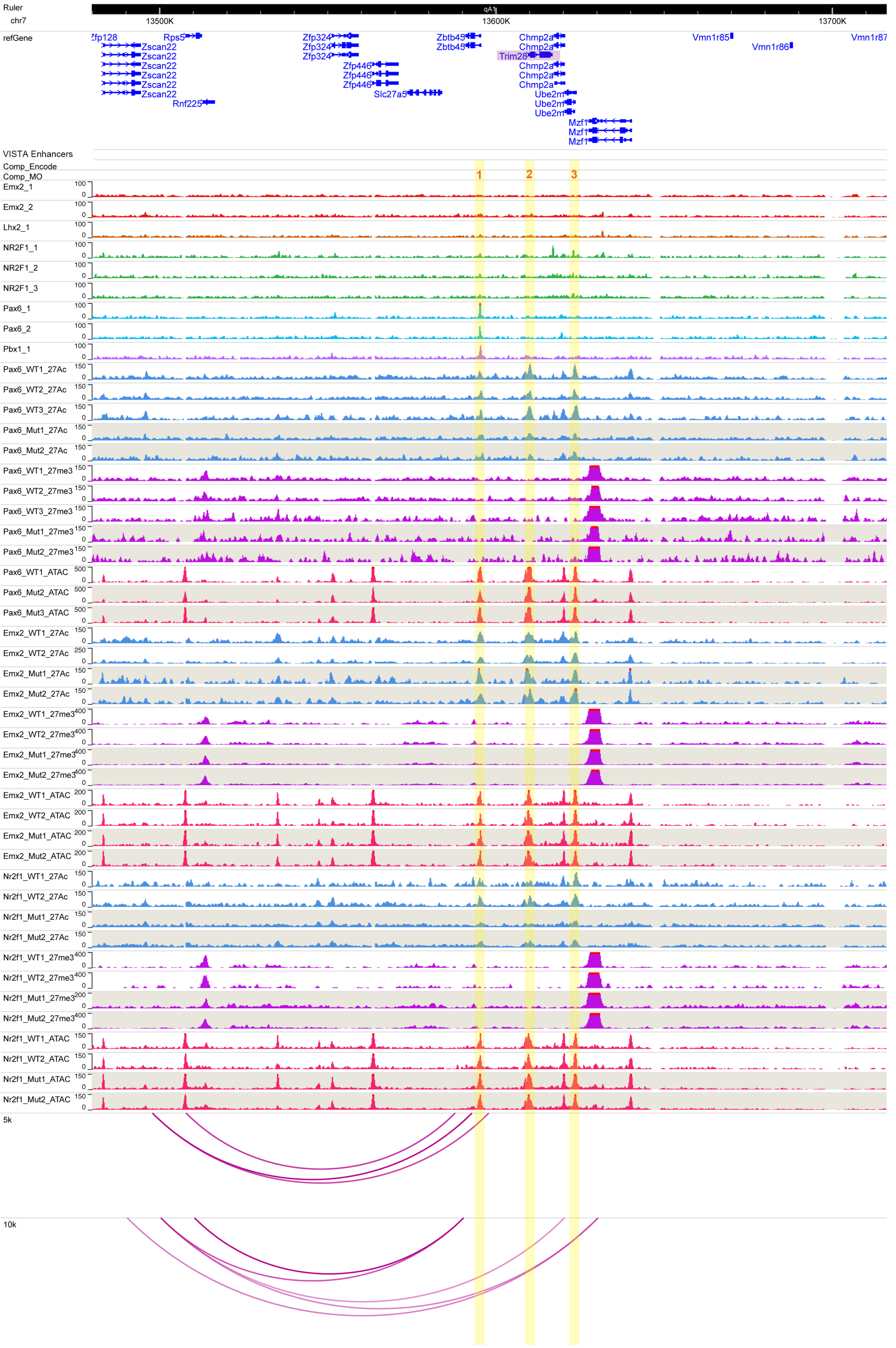

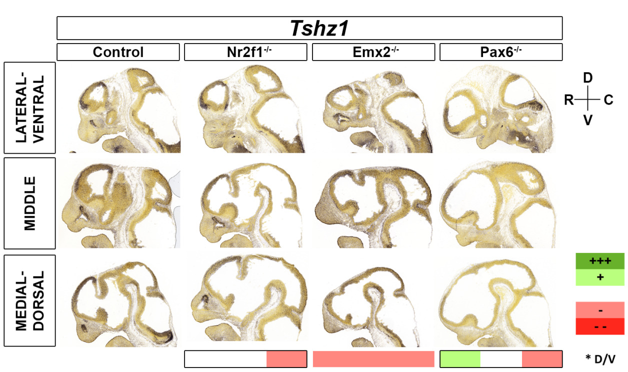


| | | Motifs | | | | | TF ChIP Seq | | | | | Differential epigenomic marks in WT vs. Mutant | | | | | | | | | Plac-Seq | Computation | Genomic features |
|---|---|---|---|---|---|---|---|---|---|---|---|---|---|---|---|---|---|---|---|---|---|---|---|
| | | | | | | | | | | | | Pax6 mutant | | | Emx2 mutant | | | Nr2f1 mutant | | | | | |
| | | Emx2 HB | Nr2f1 | Pax6 Hb | Pax6 P | Pbx1 | Emx2 | Lhx2 | Nr2f1 | Pax6 | Pbx1 | 27Ac | 27me3 | ATAC | 27Ac | 27me3 | ATAC | 27Ac | 27me3 | ATAC | | | |
| **Tshz1** | chr18:83200000-84800000 | | | | | | | | | | | | | | | | | | | | | | |
| 1 | chr18:83568794-83569877 | | | | | | | | x | | | | | Loss | | | Same | | | Same | x | | |
| 2 | chr18:83806342-83808046 | x | | x | | | x | x | | x | | Same | | Same | Loss | | Loss | Same | | Same | x | | |
| 3 | chr18:83982198-83985551 | x | | | | | x | | | | | | Same | Small Gain | | Loss | | | Same | | x | | |
| 4 | chr18:84258352-84261915 | | | | | | | | | | | | Loss | | | Gain | | | Loss | | | x | Zadh2 Gene Body |
| 5 | chr18:84757934-84760102 | | | | | | | | | | | Loss | | Same | Same | | Same | Loss | | Same | x | | Zfp407 TSS |

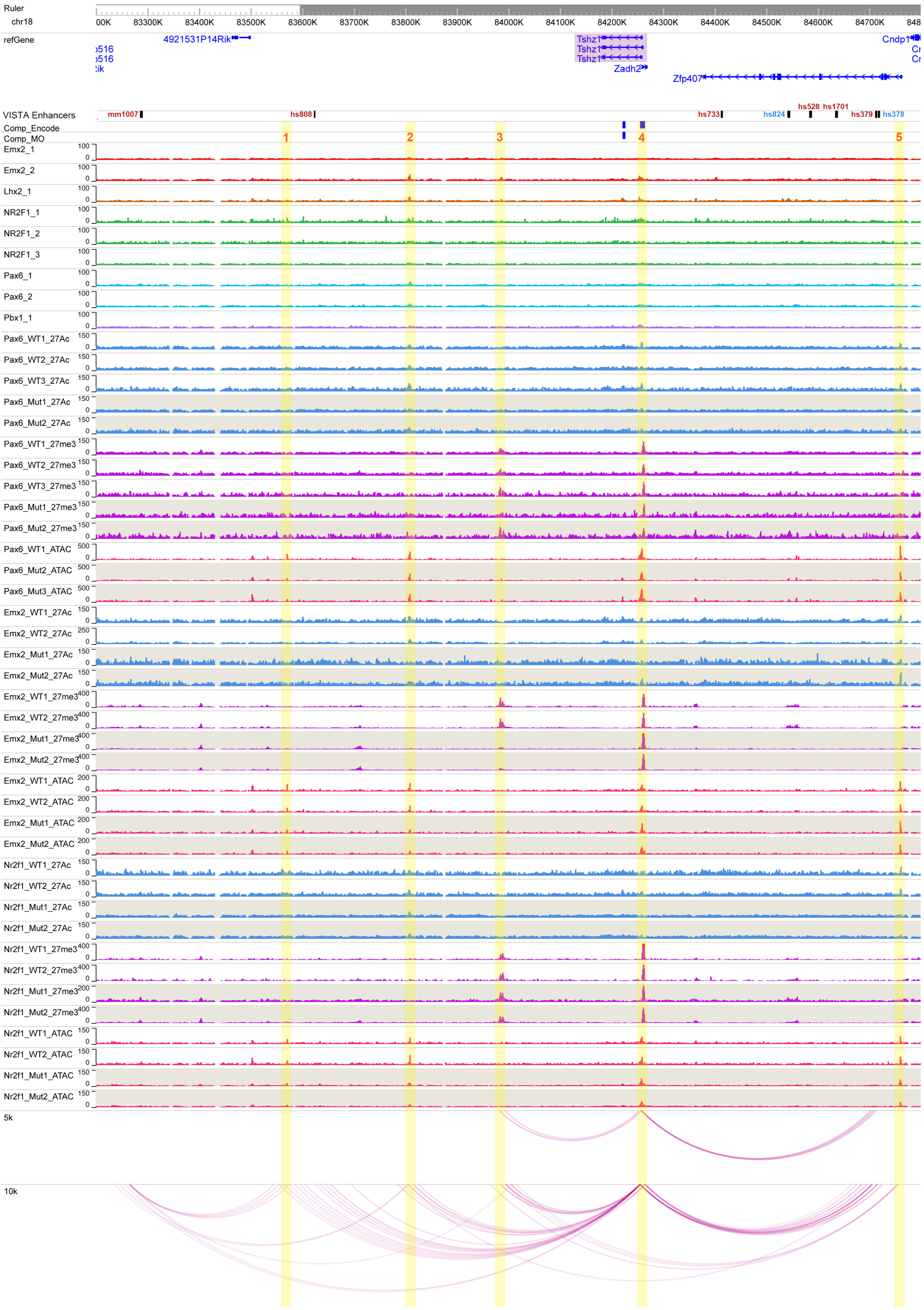

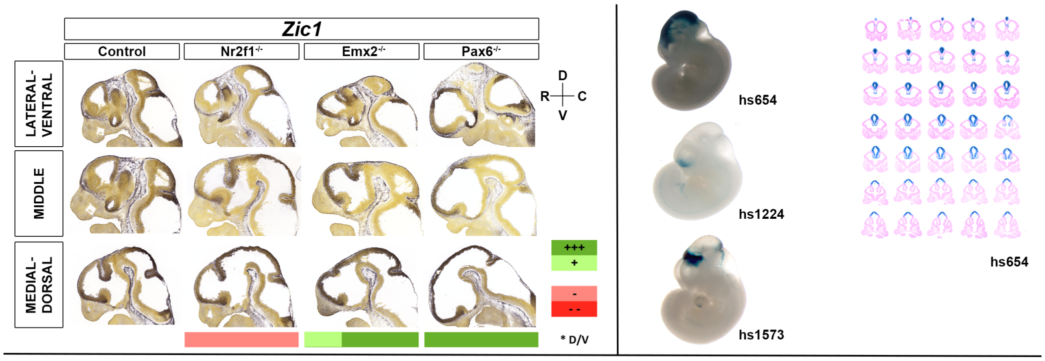


| | | Motifs | | | | | TF ChIP Seq | | | | | Differential epigenomic marks in WT vs. Mutant | | | | | | | | | Plac-Seq | Computation | Genomic features |
|---|---|---|---|---|---|---|---|---|---|---|---|---|---|---|---|---|---|---|---|---|---|---|---|
| | | | | | | | | | | | | Pax6 mutant | | | Emx2 mutant | | | Nr2f1 mutant | | | | | |
| | | Emx2 HB | Nr2f1 | Pax6 Hb | Pax6 P | Pbx1 | Emx2 | Lhx2 | Nr2f1 | Pax6 | Pbx1 | 27Ac | 27me3 | ATAC | 27Ac | 27me3 | ATAC | 27Ac | 27me3 | ATAC | | | |
| Zic1 | chr9:90500000-91900000 | | | | | | | | | | | | | | | | | | | | | | |
| 1 | chr9:90587642-90589755 | x | | | | | x | x | x | x | | | Loss | Same | | Same | Same | | Same | Same | x | x | hs654 |
| 2 | chr9:90595267-90596439 | x | | | | | x | x | | x | | | | Loss | Gain | | Same | | | Same | x | | hs739 |
| 3 | chr9:90625721-90627316 | | | | | | | | | x | | | | Loss | | | Gain | | | Same | x | | hs1549 |
| 4 | chr9:91243690-91252943 | | | | | | | | | | | | Same | Same | Gain | Loss | Gain | | Same | Gain | | x | |
| 5 | chr9:91254220-91260123 | | | | | | | | | | | Same | Loss | Same | Gain | Loss | Gain | Loss | Loss | Same | | x | Zic1 Gene body |
| 6 | chr9:91261560-91266186 | | x | | | | | | x | | | Same | Same | Loss | Gain | Loss | Gain | Same | Loss | Same | | | hs1043 + Zic4 TSS |
| 7 | chr9:91271293-91279908 | | | | | | | | | x | | Same | Same | Same | Gain | Loss | Same | Same | Same | Same | | x | Zic4 gene body |
| 8 | chr9:91297461-91302087 | x | | x | | | | | | x | | | Same | Same | Gain | Loss | Same | | Loss | Same | | | |

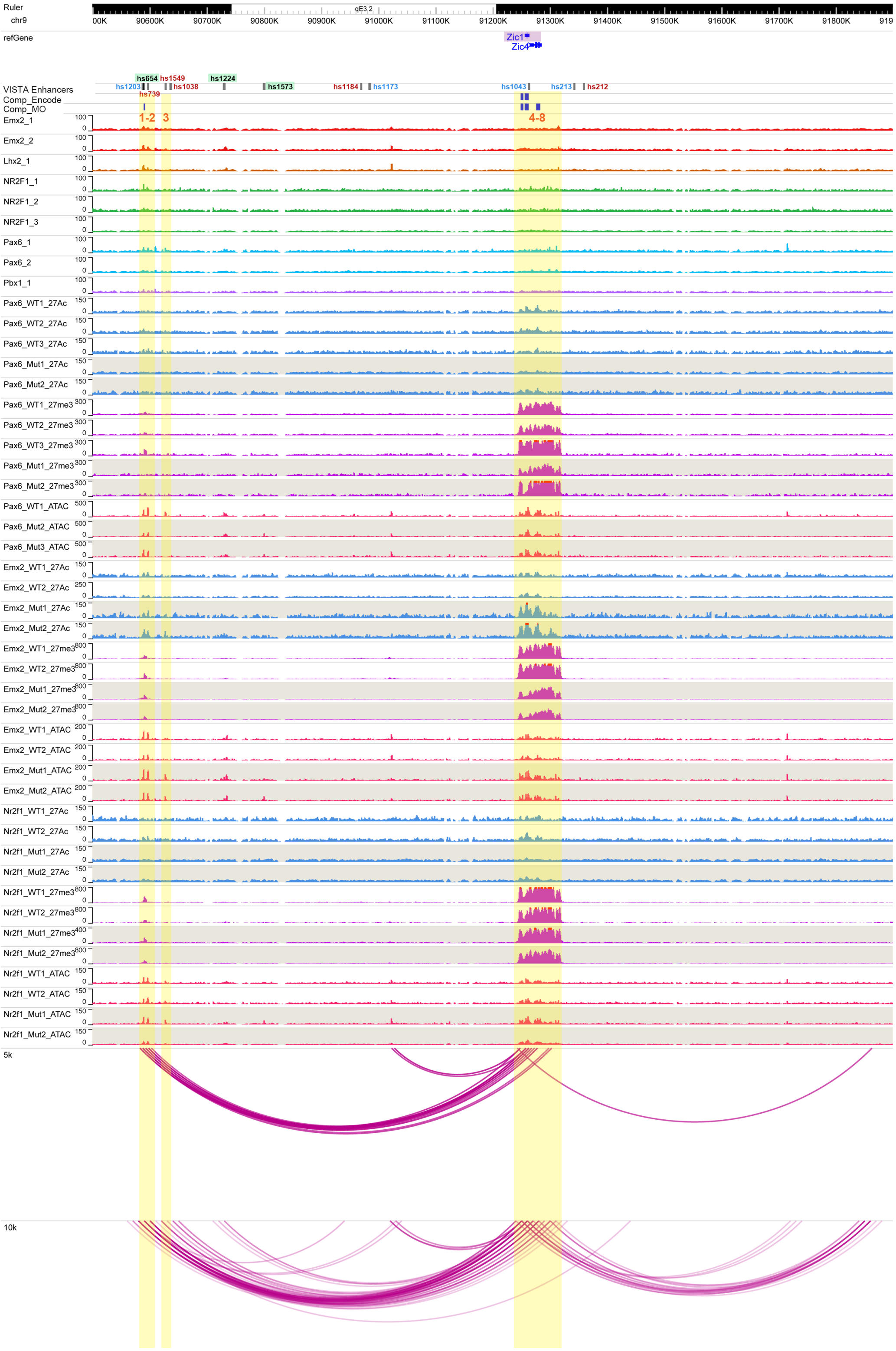

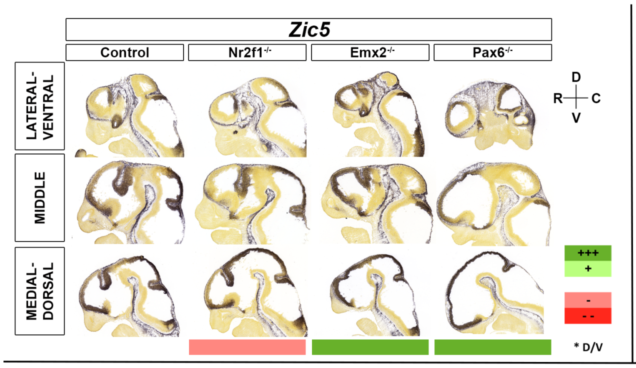


| Zic5 | chr14:122500000-123750000 | Motifs: Emx2 HB | Motifs: Nr2f1 | Motifs: Pax6 Hb | Motifs: Pax6 P | Motifs: Pbx1 | TF ChIP Seq: Emx2 | TF ChIP Seq: Lhx2 | TF ChIP Seq: Nr2f1 | TF ChIP Seq: Pax6 | TF ChIP Seq: Pbx1 | Pax6 mutant 27Ac | Pax6 mutant 27me3 | Pax6 mutant ATAC | Emx2 mutant 27Ac | Emx2 mutant 27me3 | Emx2 mutant ATAC | Nr2f1 mutant 27Ac | Nr2f1 mutant 27me3 | Nr2f1 mutant ATAC | Plac-Seq | Computation | Genomic features |
|---|---|---|---|---|---|---|---|---|---|---|---|---|---|---|---|---|---|---|---|---|---|---|---|
| 1 | chr14:122625471-122629125 | | | x | x | | x | | x | | | | | | Gain | | | | | | x | x | |
| 2 | chr14:122645738-122648063 | x | | x | x | x | x | x | | | x | | | Same | | Small loss | Small loss | | | Same | x | | |
| 3 | chr14:122772536-122775105 | x | | | | | x | | x | | x | | | Same | Gain | | Same | | | Same | x | | |
| 4 | chr14:122861504-122864073 | | | | | | | | x | | | Loss | | Same | Gain | | Same | Same | | Same | | | Zic5 TSS |
| 5 | chr14:122872971-122876529 | | | | | | x | | | | | Loss | | Same | Gain | | Same | Loss | | Same | | | Zic2 TSS |
| 6 | chr14:123381259-123383393 | | | | | | | | x | | x | Loss | | Same | Gain | | Same | Loss | | Same | x | | |

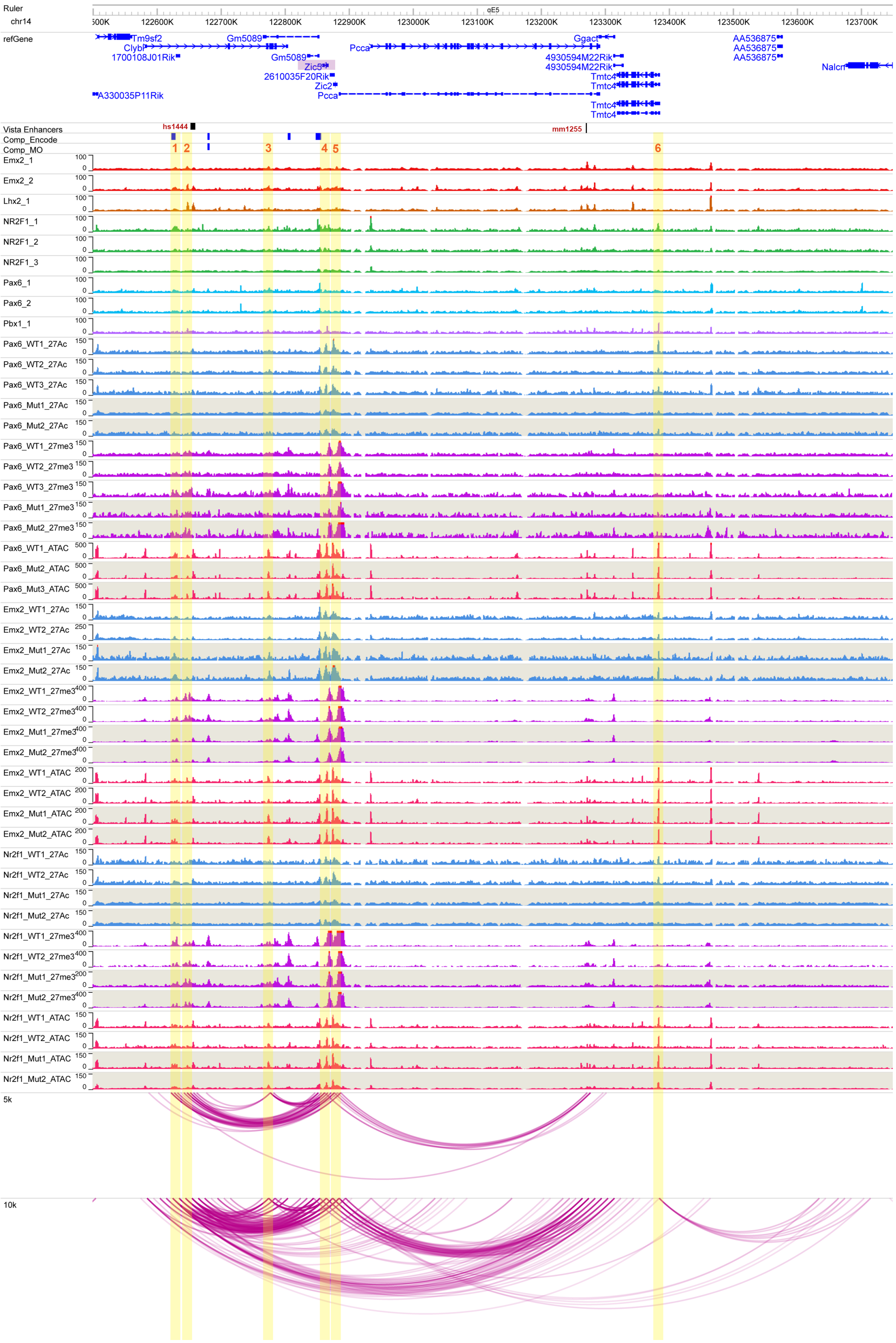

#### *Arx*

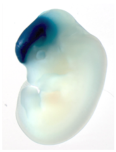
hs122

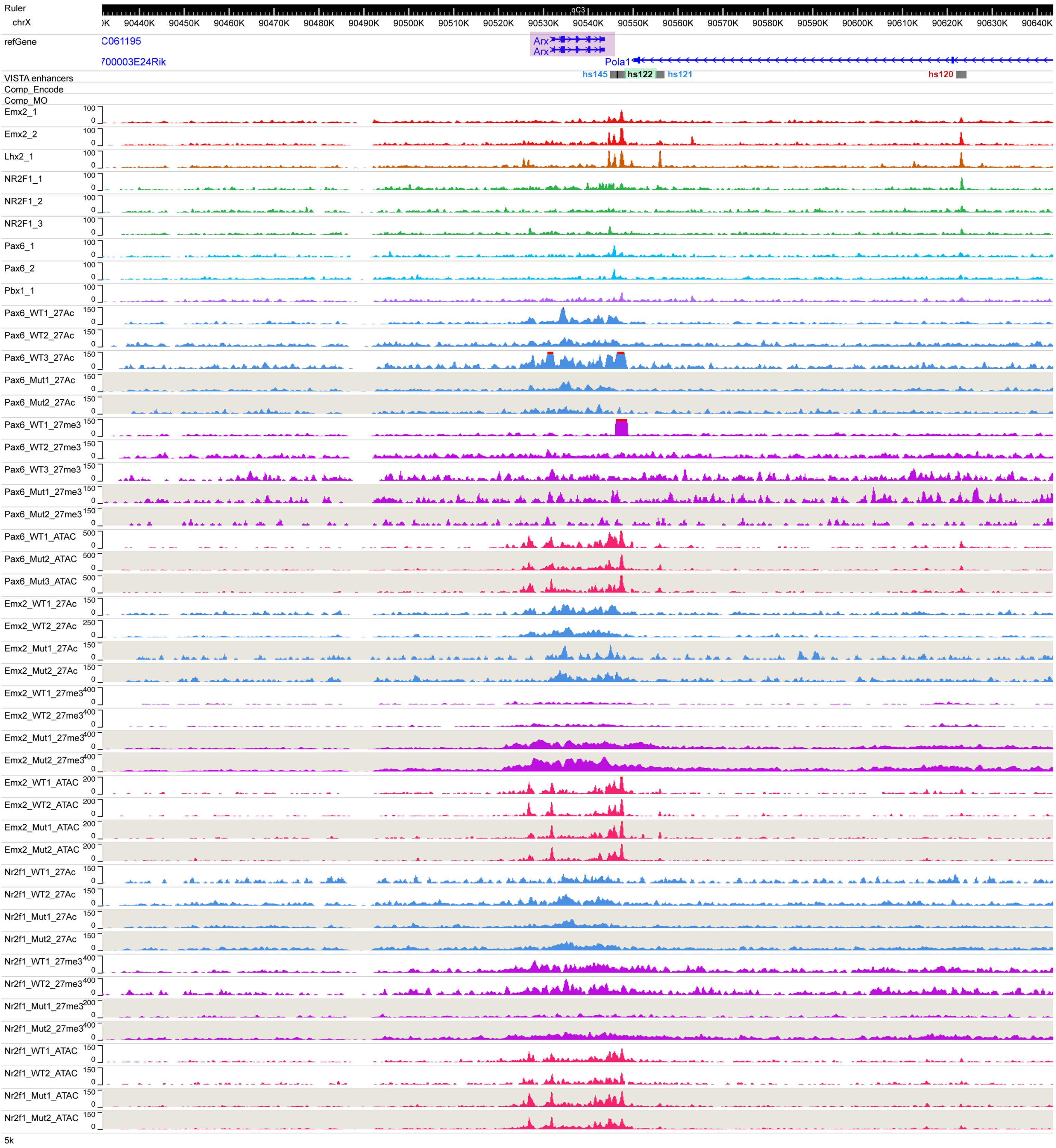
Ruler
chrX
90440K
90450K
90460K
90470K
90480K
90490K
90500K
90510K
90520K
90530K
90540K
90550K
90560K
90570K
90580K
90590K
90600K
90610K
90620K
90630K
90640K
refGene
C061195
700003E24Rik
Arx
Arx
Pola1
VISTA enhancers
hs145
hs122
hs121
hs120
Comp_Encode
Comp_MO
Emx2_1
Emx2_2
Lhx2_1
NR2F1_1
NR2F1_2
NR2F1_3
Pax6_1
Pax6_2
Pbx1_1
Pax6_WT1_27Ac
Pax6_WT2_27Ac
Pax6_WT3_27Ac
Pax6_Mut1_27Ac
Pax6_Mut2_27Ac
Pax6_WT1_27me3
Pax6_WT2_27me3
Pax6_WT3_27me3
Pax6_Mut1_27me3
Pax6_Mut2_27me3
Pax6_WT1_ATAC
Pax6_Mut2_ATAC
Pax6_Mut3_ATAC
Emx2_WT1_27Ac
Emx2_WT2_27Ac
Emx2_Mut1_27Ac
Emx2_Mut2_27Ac
Emx2_WT1_27me3
Emx2_WT2_27me3
Emx2_Mut1_27me3
Emx2_Mut2_27me3
Emx2_WT1_ATAC
Emx2_WT2_ATAC
Emx2_Mut1_ATAC
Emx2_Mut2_ATAC
Nr2f1_WT1_27Ac
Nr2f1_WT2_27Ac
Nr2f1_Mut1_27Ac
Nr2f1_Mut2_27Ac
Nr2f1_WT1_27me3
Nr2f1_WT2_27me3
Nr2f1_Mut1_27me3
Nr2f1_Mut2_27me3
Nr2f1_WT1_ATAC
Nr2f1_WT2_ATAC
Nr2f1_Mut1_ATAC
Nr2f1_Mut2_ATAC
5k
